# Asymmetrical Glyco-Engineering of Monoclonal Antibodies Provides Insights into α-Gal Immunogenicity

**DOI:** 10.64898/2025.12.15.694461

**Authors:** Gerlof P. Bosman, Anna Ehlers, Thomas Buckley, Justyna M. Dobruchowska, Javier Sastre Toraño, Arthur E. H. Bentlage, André C. Knulst, Gestur Vidarsson, Geert-Jan Boons

## Abstract

Glycosylation heterogeneity of therapeutic monoclonal antibodies (mAbs) necessitates precise glycoengineering to optimize effector functions, such as those mediated by Fcγ-receptors (FcγRs). However, current enzymatic remodeling methods are limited in scope, producing only a narrow spectrum of glycoforms and unable to achieve complete control over defined asymmetrical structures. Here, we report a multi-step, fully enzymatic platform for the solution-phase synthesis of a comprehensive library of biantennary mAb glycoforms, offering complete control over the α1,3- and α1,6-mannose arm architecture. This strategy uses a single domain of the β-*N*-acetylglucosaminidase StrH (GH20-2) to achieve regioselective hydrolysis of the GlcNAc residue on the α1,3-mannose arm of the intact antibody, thereby introducing asymmetry. Next, the remaining terminal GlcNAc residue at the α1,6-antenna can be enzymatically extended with a β1,4-galactoside and sialoside or α1,3-galactoside. Remarkably, after this extension, recombinant GnT-I can reintroduce the GlcNAc at the α1,3-mannosyl antenna, allowing precise control over glycan branching and composition. This robust, two-stage enzymatic remodeling methodology, demonstrated on infliximab and cetuximab, unlocks a broad chemical space of biantennary Fc glycans for comprehensive functional studies. We employed these well-defined glycovariants to determine the minimum structural requirements for anti-α-Gal IgE binding and to systematically evaluate binding affinities toward FcγRs, providing new insights for the rational design of next-generation immune-therapeutics.

## INTRODUCTION

Monoclonal antibodies (mAbs) are an important class of therapeutic glycoproteins used for the treatment of hematologic malignancies, solid tumors, infections, and inflammatory and autoimmune disorders.^1^ IgG antibodies consist of a variable region that recognizes a target antigen and an Fc region that binds immune cells via Fcγ-receptors (FcγRs) and complement proteins, thereby triggering antibody-dependent cell-mediated cytotoxicity (ADCC) and complement-dependent cytotoxicity (CDC), respectively. The Fc region harbors a conserved glycosylation site at N297 that is predominantly occupied by the biantennary complex-type and can vary in the degree of core-fucosylation, α- and β-galactosylation, sialylation and modification by a bisecting GlcNAc.^2^ The structure of the *N*-glycan of IgG antibodies greatly impacts its stability and function, and in particular affects binding affinities of FcγRs and C1q. Therefore, there is considerable interest to produce specific glycoforms of therapeutic mAbs.^3–6^

Several strategies have been described to modulate glycan composition in mAbs using cells with knock-ins and knock-outs of glycosylhydrolases and glycosyltransferases.^7–13^ Also inhibitors of glycosyltransferases and supplementation with additives to skew production towards a desired glycoform have been employed.^10,13–15^ Alternatively, heterogeneous *N*-glycans of mAbs have been modified by enzymatic means to produce homogeneous glycovariants. In one approach, nine glycosylhydrolases and glycosyltransferases were employed to sequentially trim and extend the glycans of an immobilized mAb.^16^ Nevertheless, these approaches have only provided a small number of glycoforms, and in particular naturally occurring asymmetrical glycans in which each antenna of a biantennary glycan has a different epitope, have been difficult to produce.

Transglycosylation is an attractive alternative approach to homogenize the *N*-glycan of a mAb. This approach employs a wild-type endo-β-*N*-acetylglucosaminidase to hydrolyze the glycosidic bond of the chitobiose core of an *N*-glycan leaving a GlcNAc or GlcNAc-Fuc moiety at the antibody. Subsequently, a mutant endo-β-*N*-acetylglucosaminidase, such as Endo-S D233A/Q or Endo-S2 D184M, performs transglycosylation in combination with a glycan-oxazoline synthesized by chemical and/or enzymatic procedures.^17–20^ The *trans*-glycosylation requires, however, 50-100 molar equivalents of chemically labile glycan-oxazoline. Furthermore, driving the reaction to completion can be difficult due to residual hydrolytic properties of glycosynthases. Here, we report an enzymatic remodeling strategy that can provide, for the first time, readily a wide variety of well-defined natural glycoforms of mAbs including symmetrical and asymmetrical structures. It exploits a single domain of the β-*N*-acetylglucosaminidase StrH (GH20) from *Streptococcus pneumoniae* R6 that selectively cleaves the terminal GlcNAc moiety at the α1,3-antennae of an A2-glycan to give a glycoform with an asymmetrical architecture (Figure 1A). Next, the remaining terminal GlcNAc residue at the α1,6-antenna can be enzymatically extended by a β1,4-galactoside and sialoside or α1,3-galactoside. Remarkably, after this extension, recombinant GnT-I can reintroduce the GlcNAc at the α1,3-mannosyl antenna, allowing precise control over glycan branching and composition. The methodology was employed to generate a comprehensive library of glycovariants of the therapeutic mAbs, infliximab and cetuximab (Figure 1B). It includes glycoforms that have different patterns of α1,3-linked galactosylation that have never been synthesized as single glycoforms. To assess the functional consequences of α1,3-galactosylation, we determined FcγR binding by SPR and the structural threshold required to trigger anti-α-Gal IgE binding using ELISA.

**Figure 1.**
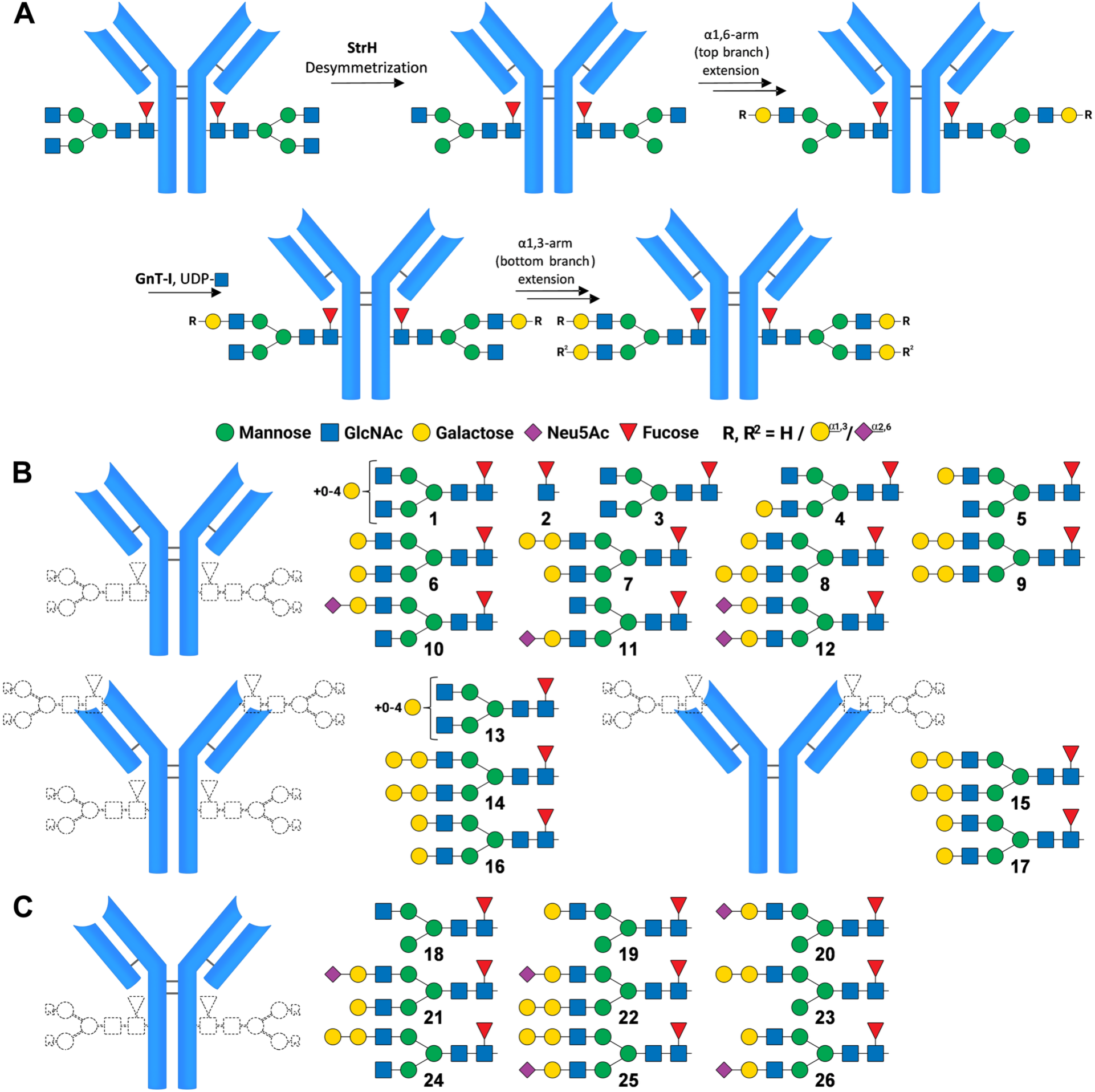
(A) General overview of the Fc region glycoremodeling strategy used to prepare the glycoforms. (B) Remodeled Fc glycans of infliximab **2**-**12** and Fc and Fab glycans of cetuximab **13**-**17** generated in this study. (C) Infliximab intermediates **18**-**26** formed during synthesis. These were characterized by LC-MS, but excluded from subsequent FcγR and IgE binding studies. Note: The right panels in (B) and (C) show the glycans detached for clarity. In reality, in the intact antibody, these identical glycan structures are symmetrically bound to both sides of the Fc or Fab region.

The α1,3-galactosyltransferase (GGTA1) is responsible for installing α1,3-linked galactosides at terminal β-galactosides on *N*-glycans.^21^ Humans possess an inactive GGTA1 gene, resulting in loss of α-Gal biosynthesis.^22^ Exposure to α-Gal through consumption of meat induces IgG antibodies that constitute approximately 1% of the total circulating IgG pool.^23^ In some individuals, sensitization *via* tick bites leads to an IgE-mediated response to α-Gal, giving rise to systemic allergic reactions to red meat. In a delayed fashion, potentially through epitope presentation on a lipid scaffold, cross-linking of IgE bound by the high-affinity FcεRI receptor with the α-Gal epitope causes degranulation of basophils and mast cells. In patients suffering from α-Gal syndrome, severe systemic allergic reactions can also occur after intravenous administration of α-galactosylated glycoproteins such as the mAbs infliximab and cetuximab.^24–28^

Cell lines commonly employed to express mAbs, such as murine myeloma (SP2/0) and CHO cells, express functional GGTA1.^21^ Infliximab and cetuximab are chimeric human–mouse IgG1 monoclonal antibodies targeting tumor necrosis factor-α (TNF-α) and the epidermal growth factor receptor (EGFR), respectively. Infliximab can carry an α-Gal epitope on its Fc glycan, whereas cetuximab possesses an additional *N*-glycosylation site in the Fab region that may also bear α-Gal. In some patients who received treatment with either infliximab or cetuximab, first-dose anaphylaxis, an IgE-mediated response, occurred.^29–40^ Notably, allergic reactions have also been reported in patients without a known history of tick bites or dietary meat allergy.^41^ The structural requirement of an α-Gal-bearing *N*-glycan to bind to IgE has not been established. Furthermore, the influence of α-Gal on binding to FcγRs has not been determined. The glycoforms developed in this study have addressed these deficiencies.

## RESULTS AND DISCUSSION

### Glycoremodeling Model Studies Exploiting Substrate Specificities of StrH and GnT-I

The first critical step in the glycoengineering approach is the selective hydrolysis of the terminal GlcNAc moiety at the α1,3-antenna of an A2-glycan (Figure 2A). We have found that a single domain of the 138 kDa β-*N*-acetylglucosaminidase StrH (GH20) can accomplish this transformation. The surface-exposed *N*-acetylhexosaminidase StrH from *Streptococcus pneumoniae R6* contributes to virulence by removing β(1,2)-linked GlcNAc from host defense molecules following cleavage of sialosides and galactosides by neuraminidase and β-galactosidase, respectively. While most GlcNAcases from the GH20 family act on β1,4-and β1,6-linkages, StrH strictly hydrolyses β1,2-glycosidic bonds. Furthermore, it is the only GH20 enzyme that contains a tandem repeat of two catalytic domains with 53% sequence identity, designated GH20-1 and GH20-2, respectively. X-ray crystallography confirmed the presence of catalytic domains that differ in substrate specificity. Notably, it revealed that a catalytically inactive mutant of the GH20-2 binding domain of StrH binds preferentially to the terminal GlcNAcβ1,2-Man motif located on the α1,3-mannosyl arm of complex *N*-glycans. In contrast, the GH20-1 domain binds to terminal GlcNAcβ1,2-Man motifs on both the α1,3-and α1,6-mannosyl antennae.^42,43^

**Figure 2.**
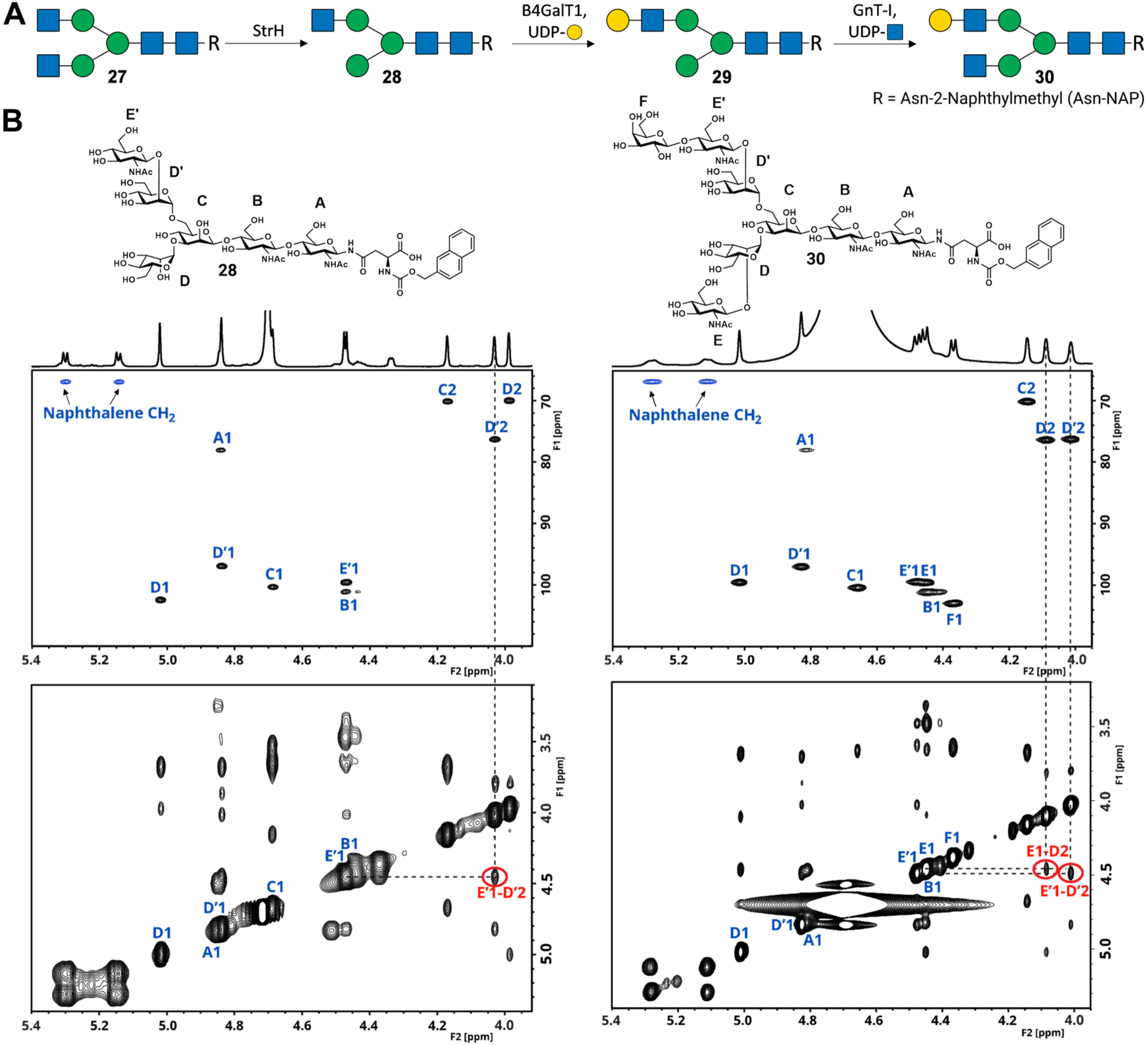
Confirmation of the branch selectivity of StrH. (A) The A2-glycan **27** is subjected to StrH treatment yielding glycan **28**. GnT-I can reinstall the GlcNAc that was earlier removed by StrH. To maintain asymmetry, glycan **28** is galactosylated by B4GalT1, yielding glycan **29**, which is subsequently GlcNAcylated by GnT-I forming glycan **30**. (B) NMR analyses of asymmetrical glycans **28** and **30**. Left panel shows (partial) HSQC spectrum of glycan **28**. Right panel: HSQC spectrum of glycan **30**. Representative NOESY spectra displayed below HSQC. Additional anomeric peaks noted E1 and F1 indicate installation of two more glycosides. Installation of GlcNAc by GnT-I on glycan **30** induces a downfield shift in the α1,3 Man-D H2 (D2). NOESY spectra indicate a single through-space correlation between GnT-II GlcNAc-E’ H1 and the Man-D’ H2 (E’1 → D’2). Reinstallation of GnT-I GlcNAc-E indicates a new additional through-space correlation between GlcNAc-E H1 and Man D H2 (E1 → D2). See for full analyses Figures S2, S4, and S5.

To exploit GH20-2’s substrate specificity, the gene encoding the GH20-2 domain was cloned and the corresponding enzyme was produced. The GH20-2 domain (41 kDa) (GH20-2, from here on referred to as “StrH”) was recombinantly expressed in *E. coli* as a fusion protein with maltose-binding protein (MBP). After sequential purification via tandem affinity chromatography (Ni-NTA and Strep-Tactin) and final size-exclusion polishing, we used the pure enzyme for subsequent evaluations.

To establish the glycoengineering approach, we first performed model studies using *N*-glycan **27**, which was derived from a sialylglycopeptide (SGP) isolated from egg yolk powder.^44^ Glycan **27** was treated with recombinant StrH in sodium acetate buffer, and reaction progress was monitored by HILIC-MS. Analysis revealed the formation of a single product, glycan **28**, lacking a single GlcNAc residue. The product was fully characterized by LC-MS and NMR spectroscopy, including ^1^H–^13^C HSQC, TOCSY, NOESY, and COSY characterization (Figures 2B and S1–S5), which collectively demonstrated the presence of a single glycoform. These analyses confirmed the selective hydrolysis of the GlcNAc-E residue from the α1,3-mannosyl antenna, while the corresponding GlcNAc-E’ residue on the α1,6-mannose arm remained intact. Consistent with this assignment, the ^1^H–^13^C HSQC spectrum lacked an anomeric assignment corresponding to the GlcNAc-E residue on the α1,3 branch (Figure 2B, left upper panel). Furthermore, diagnostic NOEs were observed between H1 of the GlcNAc-E’ residue on the α1,6 antenna and H2 of the corresponding Man-D’ residue (δ 4.47–4.03, E′1→D′2), confirming retention of the GlcNAc (E’) on the α1,6-arm (Figure 2B, left lower panel). COSY spectra enabled assignment of the H2 resonances of the individual mannose residues, including H2 of the Man-C (δ 4.17), H2 of Man-D (δ 3.99), and H2 of Man-D’ (δ 4.03) (Figures S1-S5). Glycan **28** was subsequently galactosylated using B4GalT1 to install a β1,4-linked galactose residue, affording glycan **29**. LC-MS and NMR analyses indicated complete and regioselective conversion to a single asymmetrical glycoform (Figures S2 and S3).

α1,3-Mannosyl-glycoprotein 2-β-*N*-acetylglucosaminyltransferase (GnT-I) is the gatekeeper enzyme for converting high-mannose glycans into hybrid and complex *N*-glycans. Canonically, GnT-I installs a β1,2-GlcNAc residue on the α1,3-mannosyl arm of the Man5GlcNAc2 core, thereby enabling subsequent processing by Golgi α-mannosidase II and GnT-II, ultimately leading to the formation of complex N-glycan structures.^45,46^ Several studies have demonstrated that GnT-I exhibits relaxed acceptor specificity. For example, Man6-based hybrid glycans have been observed in cells lacking Golgi α-mannosidase IA, IB, and IC activity, implying that GnT-I can act on non-canonical substrates.^47^ In addition, GnT-I has been reported to modify Man3 structures.^48^ These observations prompted us to investigate whether recombinant human GnT-I could reinstall a β1,2-GlcNAc residue at the terminal α1,3-mannosyl branch of glycan **29**. Gratifyingly, incubation of glycan **29** with UDP-GlcNAc and recombinant human GnT-I resulted in quantitative formation of glycan **30**, as determined by LC-MS analysis (Figure S4). The structure of glycan **30** was confirmed by LC-MS and NMR spectroscopy. Installation of the GlcNAc-E residue on the α1,3-mannose branch resulted in a characteristic downfield shift of the C2/H2 of Man-D(δC-2 70.0, δH-2 3.97 → δC-2 76.4, δH-2 4.09), consistent with glycosylation at O-2 and in agreement with previously reported data for structurally related glycans.^49^ Additional support for the assignment was provided by the appearance of a diagnostic NOEs between H1 of the newly installed GlcNAc-E residue and H2 of the Man-D residue (δ 4.45–4.09, E1→D2), confirming formation of the β1,2 linkage on the Man-D arm (Figure 2B). The ^1^H–^13^C HSQC spectrum displayed a single set of branch-specific anomeric shifts, indicating the presence of a single glycoform. Collectively, these results demonstrate that the combination of selective GlcNAc removal by StrH and subsequent reinstallation by GnT-I, provides an effective strategy for generating structurally defined asymmetrical *N*-glycans. Full spectra, peak assignments, and annotations for glycans **27**–**30** are provided in Figures S1–S4 and the Experimental Section.

### Glycoremodeling of Infliximab

Next, we explored the scope of the asymmetrical glycan remodeling strategy for preparing a wide range of glycoforms of therapeutic mAbs. For this purpose, we employed infliximab, which is a chimeric human–mouse IgG1 monoclonal antibody used to treat autoimmune diseases by neutralizing TNFα. It is commonly employed for rheumatoid arthritis, Crohn’s disease, ulcerative colitis, ankylosing spondylitis, psoriasis, and psoriatic arthritis. Despite its clinical efficacy, infliximab is immunogenic and has been associated with infusion-related hypersensitivity reactions, including IgE-mediated responses. First, infliximab was subjected to a neuraminidase (*C. perfringens*) and a β-galactosidase (BgaA, *S. pneumoniae*)^50^ to yield a homogeneous A2F-glycan at N297 of the Fc region. The latter glycoform **3** was subjected to StrH in 50 mM 3-(*N*-morpholino)propanesulfonic acid (MOPS) buffer (pH 6.5) for 8 h and then isolated by Protein A affinity chromatography. The mAb **18** was released from the resin with 100 mM glycine buffer (pH 3), and after adjusting the pH to neutral, the sample was concentrated by spin filtration. It was found that even after exposure of the mAb to StrH for 72 h, a single GlcNAc remained. The mAb was analyzed by LC-MS using a reverse-phase Acquity 300Å column (Waters) and a Q-TOF mass detector. Prior to analysis, the mAbs were treated by the enzyme IdeS that cleaves below the hinge region,^51^ resulting in a single Fc fragment, confirming removal of one GlcNAc moiety. It showed the expected mass (Figure 3C) and matched the calculated mass in Table S2.

**Figure 3.**
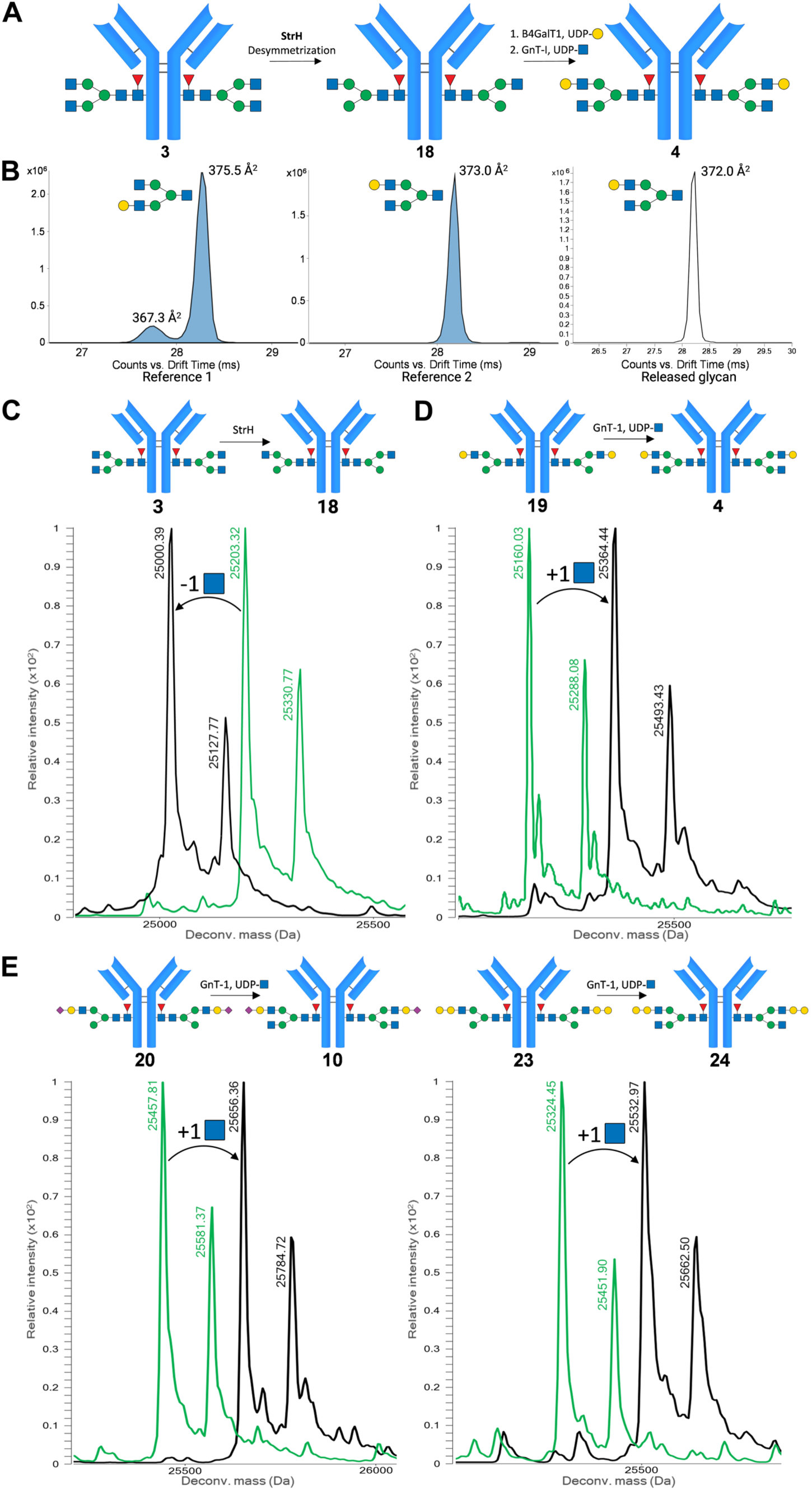
(A) Infliximab was sequentially trimmed with neuraminidase and β-galactosidase to establish a homogeneous A2F-glycan core, mAb **3**. Regioselective removal of a single GlcNAc residue by StrH yielded asymmetrical mAb **18**. Subsequent B4GalT1 catalyzed galactosylation of the α1,6-arm afforded mAb **19**, followed by GnT-I-mediated restoration of the α1,3-branch to complete the synthesis of asymmetrical mAb **4**. (B) The released Fc glycan was 2AA-labelled, subjected to IM-MS analysis, which showed only the A2G1(ɑ1,6)-glycan. The PGC-MS trace of the released Fc glycan, and the tryptic glycopeptide LC-MS analyses are shown in Figures S6 and S7 (C) LC-MS trace of IdeS-released Fc fragment of mAbs **3** and **18**, a single GlcNAc is released. Full spectra are provided in Figures S14 and S29. (D) Installation of GlcNAc by GnT-I on mAb **19** carrying an GlcNAcacted and galactosylated α1,6-branch. (E) Capping the α1,6-branch through installation of a Neu5Ac or α-Gal does not hinder GnT-I’s conversion of **20** and **23** and thus, in general, capping does not interfere with reactivation of the α1,3-mannose arm. Observed masses agreed with the theoretical masses, as shown in Table S2. Full deconvoluted spectra of mAbs **4**, **10**, **19**, **20**, **23**, and **24** are shown in Figures S15, S21, S30, S31, S34, and S35, respectively. Note: Each spectrum displays two distinct peaks per glycoform representing variants with and without the C-terminal lysine residue. This truncation is driven by naturally occurring carboxypeptidases during production and is frequently observed in therapeutic antibodies like infliximab and cetuximab.

Next, the asymmetrical glycan of mAb **18** was galactosylated using B4GalT1 and UDP-Gal to install a β1,4-galactoside, affording mAb **19**, followed by reinstallation of GlcNAc at the α1,3-mannosyl branch using recombinant GnT-I to yield mAb **4** (Figure 3A). After each step, we purified the mAb by Protein A affinity chromatography, then subjected it to IdeS, followed by LC-MS analysis of the Fc fragment. In each step, the formation of a single glycoform was observed with the correct mass.

We confirmed the structural integrity of the glycan of mAb **4** by ion mobility mass spectrometry (IM-MS). In IM-MS, ions are introduced into a drift tube and separated under an electric field based on their mobility, which depends on the charge and size of the ions. The drift tube is filled with a buffer gas (nitrogen) that collides with the ions during migration through the drift tube, providing a separation based on ion shape and a unique arrival time at the end of the drift tube. Separation by shape can be an effective way to separate and identify isomeric structures. The arrival time at the end of the drift tube is directly related to the rotationally averaged collision cross section (CCS), which is an intrinsic value of the ion that can be used for identification. Recently, we have shown that glycan isomers have specific gas-phase conformers that can be separated by ion mobility. The unique arrival time distributions (ATDs) of the conformers can be used as a fingerprint for unambiguous identification of exact structures.^52^ By employing appropriate standards, isomeric identification can be made. Thus, the *N*-glycan of mAb **4** was released by wild-type Endo-S2 and then labeled with 2-AA. The LC-MS analysis using a Porous Graphitic Carbon (PGC) column run showed one glycan peak, indicative of a single, homogeneous, released glycan (Figure S6). ATDs were compared to those of a collection of well-defined synthetic *N*-glycan standards (Figure 3B). The ATD of the A2G1(α1,3)-glycan has a typical shoulder, whereas this signal was not observed for the A2G1(1,6)-glycan (see SI for full structures and glycan nomenclature). The ATD of the released glycan matched that of A2G1(α1,6)-glycan confirming the regioselectivity of StrH. Tryptic glycopeptide LC-MS analysis (Figure S7 and Table S1) confirmed that mAb **4** contains no other *N*-glycan than the A2G1F-glycan.

Glycoform **19** is a key intermediate for the preparation of a wide variety of glycoforms (Figure S10). We envisaged that the terminal β(1,4)-galactoside of **19** can be modified by a 2,6-sialoside or α1,3-galactoside to give mAbs **20** and **23**, respectively (Figure 4). These modifications block further enzymatic extension, allowing reinstallation of GlcNAc at the α1,3-mannosyl branch by recombinant GnT-I, which can be followed by further enzymatic extension of the resulting GlcNAc moiety. At an appropriate stage of the synthesis, the terminal 2,6-sialoside or α1,3-Gal can be removed by enzymatic hydrolysis to give further glycoforms.

**Figure 4.**
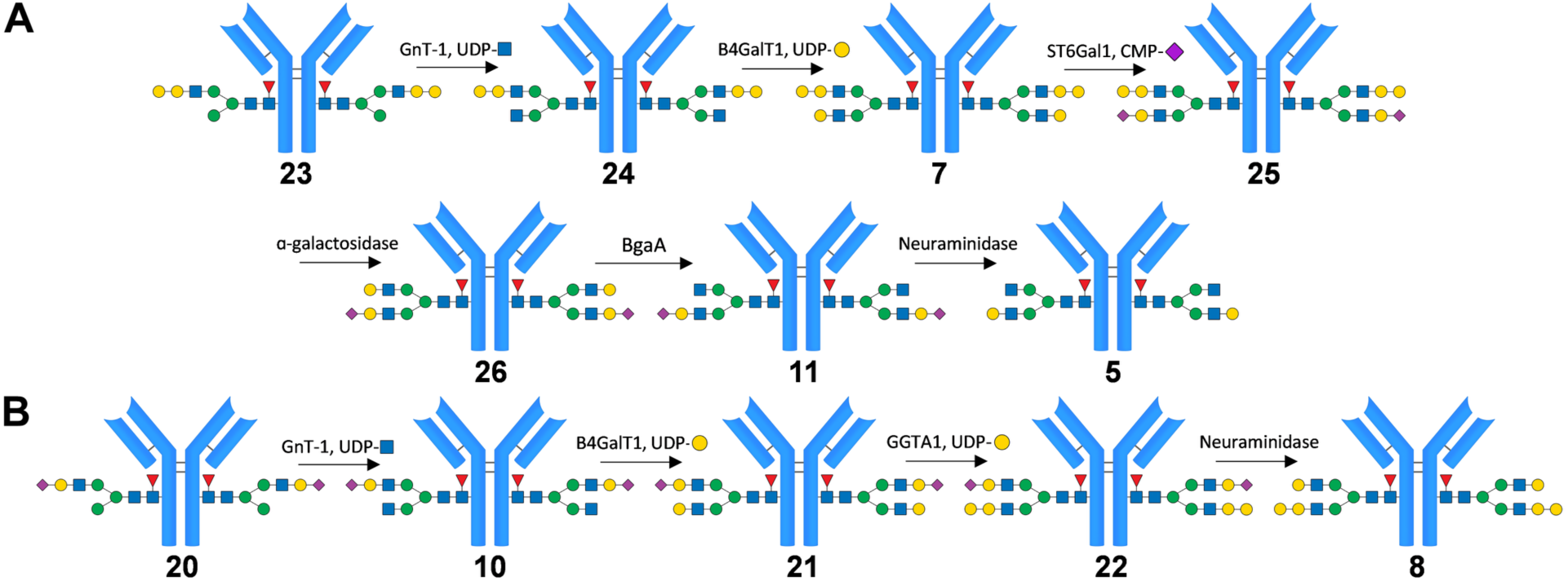
Asymmetrical glycoremodeling step-by-step procedure. The terminal β1,4-galactoside on the α1,6-arm of **19** is selectively capped using mouse-derived recombinant GGTA1 to introduce an α1,3-galactoside yielding **23,** or human ST6Gal1 to introduce a 2,6-sialoside providing **20**. (A) The α1,6-branch α-Gal strategically blocks further enzymatic extension on the α1,6-arm, enabling recombinant GnT-I to reinstall β1,2-GlcNAc at the open α1,3-mannosyl branch. Downstream iterations of β4GalT1 galactosylation and ST6Gal1 sialylation yield **7** and **25**. Enzymatic trimming by an α-galactosidase, BgaA and a neuraminidase results in intermediate **26** and final products **11** and **5**. (B) The α1,6-branched sialoside in **20**, similar to compound **23**, does not hinder GnT-I from reattaching the β1,2-GlcNAc. Subsequently, β4GalT1 catalyzes galactosylation, and ST6Gal1 adds sialic acid, producing compounds **21** and **22**, respectively. Final treatment with neuraminidase completes this synthesis, resulting in compound **8**.

As the gene for a functional version of the enzyme ɑ1,3-galactosyltransferase GGTA1 is lacking in humans, we retrieved the GGTA1 gene from the mouse (*M. musculus*) genome. The luminal portion of GGTA1 was cloned and produced as a Green Fluorescent Protein (GFP) fusion protein in HEK cells.^21,53^ After extraction and purification, the enzyme showed high activity in the *in vitro* modification of mAb glycans and efficiently converted mAb **19** to mAb **23** (Figure 4A).

mAb **19** could also be converted into the corresponding 2,6-sialoside by treatment with recombinant human ST6Gal1 in the presence of Calf Intestinal Alkaline Phosphatase (CIAP) and cytidine monophosphate (CMP)-Neu5Ac to give **20** (Figure 4B). Three consecutive daily additions of fresh ST6Gal1, CIAP, and CMP-Neu5Ac were needed to drive the sialylation to completion. It is well-documented that ST6Gal1 prefers the α1,3-mannose antenna, providing a rationale for the rather slow modification of **19** by this enzyme. Alternative sialyl transferases were explored including the bacterial, wildtype Pd26ST and Psp26ST A366G/A235M.^54–56^ LC-MS analysis showed, however, inferior performance of these enzymes (Figures S8 and S9).

Gratifyingly, GnT-I exhibits the required substrate tolerance and can reinstall the β1,2-GlcNAc moiety regardless of whether the β-galactoside at the α1,6-mannose is capped by Neu5Ac or α-Gal (Figure 3E). As a result, the glycoforms **20** and **10** could readily be converted into **23** and **24**, respectively. The latter two mAbs were incubated with B4GalT1 in the presence of UDP-Gal, resulting in the facile formation of **21** and **7**, respectively (Figure 4).

At this point, the glycoremodeling of **21** and **7** diverged. The α1,3-arm of **7** could readily be capped with Neu5Ac by treatment with ST6Gal1 and CMP-Neu5Ac to produce **25** (Figure 4B). Subsequently, the epitope at the α1,6-arm could be trimmed by first incubation with an α-galactosidase to give **26**, which was further exposed to the β-galactosidase, BgaA, generating mAb **11**. Finally, removal of the sialoside from the α1,3-arm of **11** by a neuraminidase yielded glycoform **5**. Alternatively, α-galactosylation of the terminal β-galactoside of **21** by GGTA1 in a one-pot reaction gave **22**. Finally, treatment of **22** with a neuraminidase provided **8**.

The workflow was straightforward for generating symmetrical glycoforms. Figure S11 shows the synthesis of infliximab-based compounds **2**, **3**, **6**, **9**, and **12**. Compound **3** was obtained in one step by incubating with the galactosidase BgaA, yielding the homogeneous A2F-glycan. Incubation of unmodified infliximab (**1**) with B4GalT1 afforded the fully galactosylated mAb **6** in a single step. Subsequent modification of **6** with ST6Gal1 or GGTA1 generated the fully sialylated or α-galactosylated mAbs **12** and **9**, respectively. The (partially) deglycosylated **2** was obtained after treating **1** with wild-type Endo-S2 and served as a control.

The complete procedure starting from the unmodified mAb **1** is outlined in Figure S10. All reactions were monitored by LC-MS; the deconvoluted spectra (Figures S12–S37) agreed with the theoretical masses (Tables S2 and S3).

### Enzymatic Glycoremodeling of the Fc and Fab-Glycans of Cetuximab

To establish the importance of glycan modification of the Fab fragment of a mAbs by α-Gal epitopes, we also made well-defined glycoforms of cetuximab. The latter mAb, which, like infliximab, is also produced in the SP2/0 cell line, carries a Fab *N*-glycan at N88. The incidence of hypersensitivity reactions to cetuximab (**13**) is substantially higher than for infliximab,^29–40^ which may be due to its exposed Fab-glycan. Two cetuximab glycovariants were prepared having well-defined α-Gal content at the Fc and Fab region (**14**) and the Fab fragment only (**15**) (Figures 6 and S11). As control mAb, the glycoforms **16** and **17** were prepared that have a terminal β1,4-linked galactoside. Because of the high Fc sequence homology between infliximab and cetuximab (see Tables S2 and S3), the impact of glycan variations can be compared.

Unmodified cetuximab **13**, in which at least 80% of Fab glycans contain α-Gal (Figure S24), was incubated with α-galactosidase and B4GalT1 and UDP-Gal, yielding glycovariant **16**, which has only β-linked terminal galactosides. Another batch of unmodified cetuximab **13** was incubated with B4GalT1 and GGTA1 to obtain fully α-galactosylated cetuximab **14** (Figure 5).

**Figure 5.**
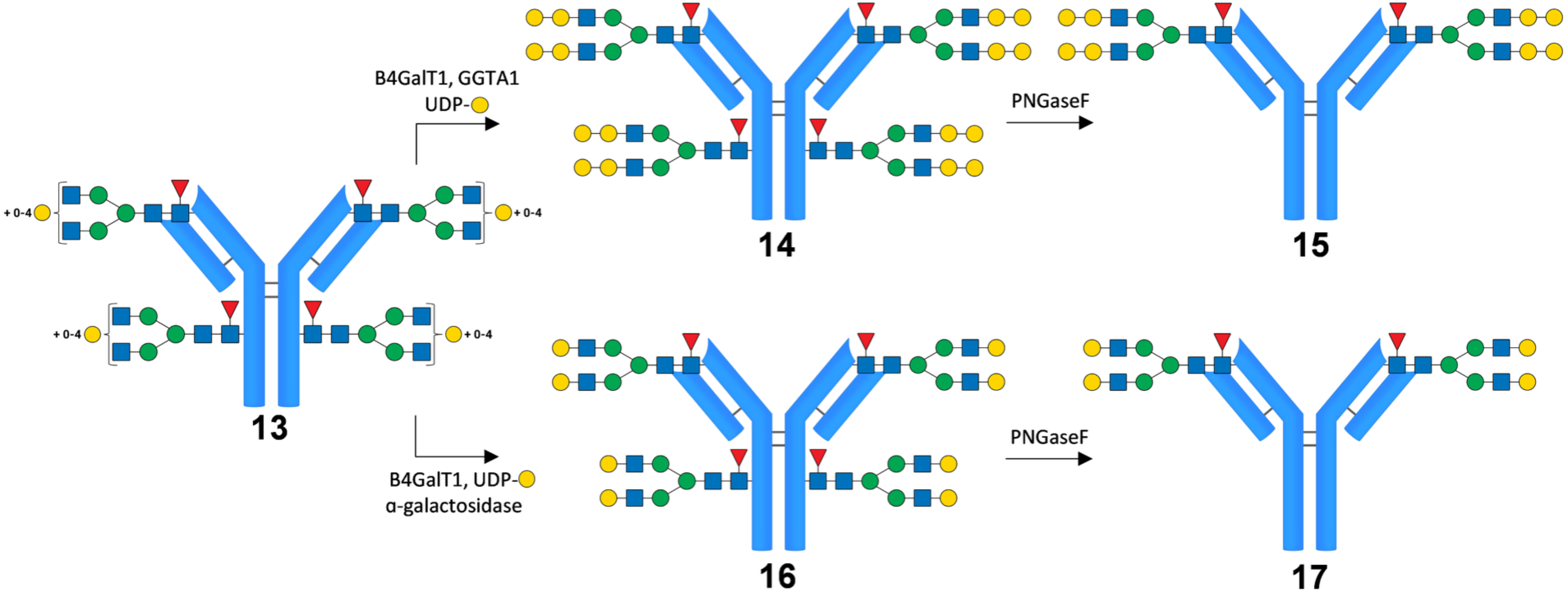
Symmetrical cetuximab variants with defined Fc and Fab glycosylation. mAb **13** is unmodified cetuximab and contains some degree of α-galactosylation (Figure S24). Selective removal of the Fc glycan was achieved by incubation with PNGase-F.

To determine the effect of the α-Gal epitope on the Fab-glycan, we sought to selectively remove the Fc glycan from **14** and **16**. We evaluated a panel of enzymes for selective Fc glycan removal. In test reactions, mAb **16** was treated with Endo-F3 WT, Endo-S WT, Endo-S2 WT, and PNGase-F. The enzymes Endo-F3 WT, Endo-S WT, Endo-S2 WT inadequately removed the Fc glycan selectively from **16**, and in all cases both Fab and Fc glycans were partially removed (data not shown). However, PNGase-F was capable of selective removal of the Fc glycan of cetuximab and, even after prolonged incubation times, left the Fab-glycan untouched. Therefore, PNGase-F was used to modify **16**, yielding a mAb with only a Fab-glycan and no α-Gal epitope (**17**). Similarly, PNGase-F digestion of **14** afforded **15**, a cetuximab variant retaining only the Fab-linked α-Gal epitope.

### IgE Binding Profiling of α-Gal-Containing Infliximab and Cetuximab Variants

Alpha-Gal syndrome is an allergic disorder, most commonly triggered by tick bites, characterized by a severe, potentially life-threatening IgE-mediated hypersensitivity to α-Gal epitopes, typically manifesting after consumption of red meat. These structures are naturally present in most non-primate mammalian tissues and their derived products.^32,57^ Both Chinese hamster ovary (CHO) and murine myeloma cell lines, which are commonly used to express biotherapeutics, including monoclonal antibodies, can produce proteins containing α-Gal epitopes.^21,58^ Administration of therapeutic glycoproteins having an α-Gal epitope can lead to an IgE-mediated anaphylaxis response. The commercial antibody cetuximab (Erbitux) contains N-glycans in both its Fc and Fab regions. Because the drug is manufactured in a murine myeloma cell line, these glycans contain the α-Gal moiety, which is responsible for most anaphylactic reactions in treated patients.^31^

Little is known about the structural requirements of α-Gal-containing glycans for binding to IgE. In this study, we generated seventeen structurally defined glycoforms of infliximab and cetuximab, including α-Gal-containing glycans in the Fc and Fab regions. The panel of mAbs was employed to examine binding to recombinant and patient-derived IgE to establish the structural requirements for recognition. To evaluate IgE reactivity, the infliximab and cetuximab variants were coated onto 96-well plates. Infliximab variants **1–6** and **10–12**, along with cetuximab variants **16** and **17**, lack the α-Gal epitope and served as negative controls (Figure 6). The plates were then incubated with sera from red-meat-allergic individuals, while an anti-α-Gal IgE monoclonal antibody (clone m86; Absolute Antibody) was used as a positive control. Bound IgE was subsequently detected using an HRP-conjugated goat anti-human IgE secondary antibody. Final IgE binding was evaluated by measuring the optical density and normalizing these values against the mean of the negative controls.

**Figure 6.**
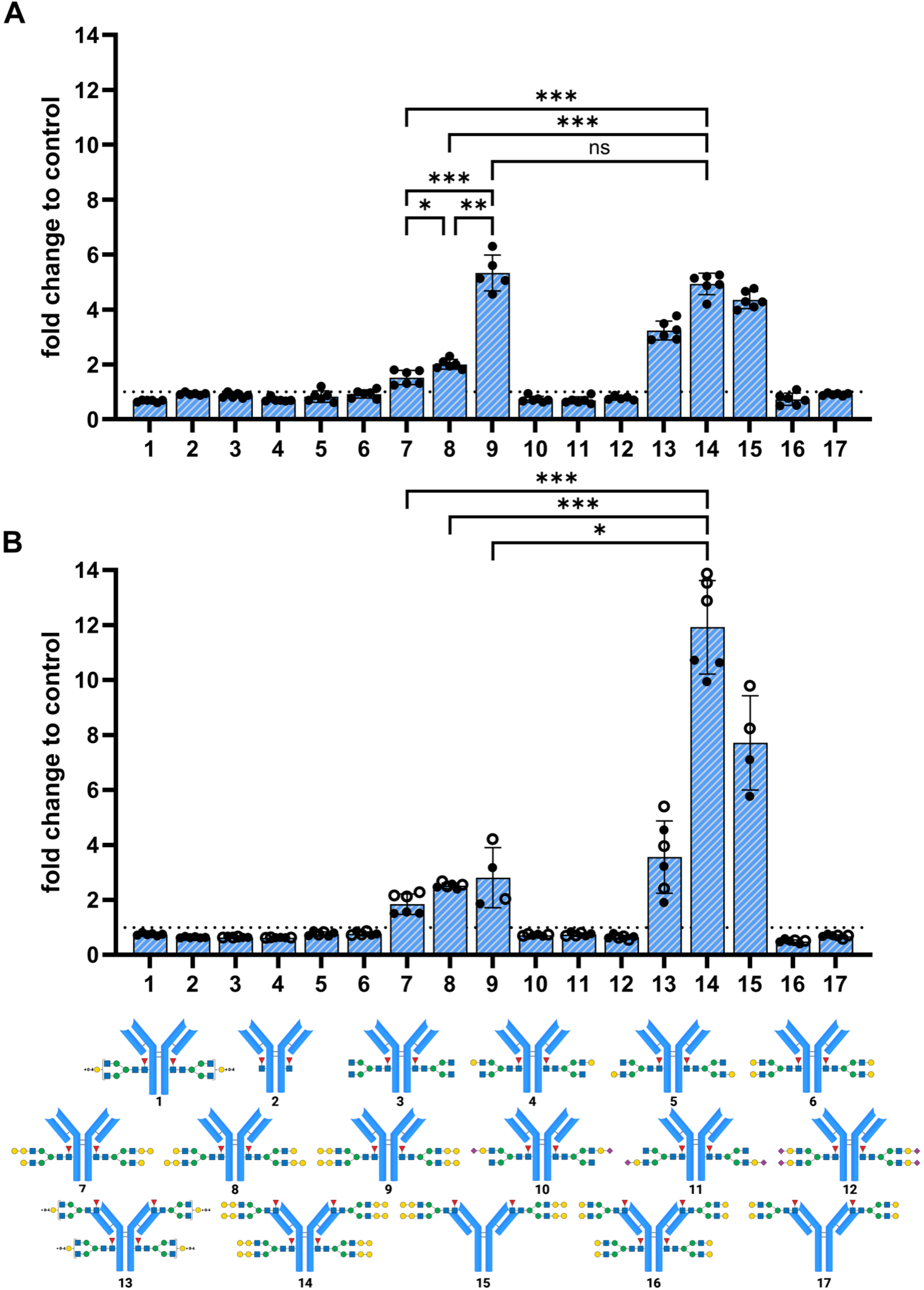
Evaluation of recombinant and patient-derived IgE binding to engineered α-Gal-containing antibody glycoforms. Normalized ELISA values (OD450 nm) displaying the recognition profiles of recombinant anti-α-Gal IgE (clone m86, n = 6, triplicate measurements on two consecutive days) and sera from red meat allergic patients (n = 2, triplicate measurements, duplicate measurements for **9** and **15**) across various infliximab and cetuximab variants. (A) Recombinant IgE exhibits a clear preference for bis-α-galactosylated structures over mono-α-galactosylated structures, regardless of whether they reside on the Fc or Fab regions. (B) Patient-derived IgE demonstrates a stringent structural preference for Fab-linked α-Gal epitopes over Fc-linked epitopes. Control variants lacking the α-Gal epitope (infliximab 1–6, 10–12; cetuximab 16, 17) showed no baseline binding. The same symbol represents the same serum. All data are shown as mean with standard deviation. Statistical analysis was performed with mixed-effects analysis, and the Tukey test was used for correction for multiple testing. Significant differences between specific variants are indicated (***p < 0.001, **p < 0.01, *p < 0.05), and a restricted number of comparisons were selected for visual presentation.

The recombinant IgE monoclonal antibody showed markedly higher responsiveness to infliximab variants **7**, **8**, and **9** containing either mono-or bis-α-galactosylated glycans than to the controls. Furthermore, bis-α-galactosylated glycoform **9** gave significantly higher responses than mAbs with mono-α-galactosylated glycans **7** and **8**. The presentation of the α-Gal residue at either the α1,3-(**7**) or α1,6-antennae (**8**) of the Fc glycan did not greatly impact binding. Interestingly, the cetuximab variants **14** and **15**, having either bis-α-galactosylated glycans in the Fc and Fab regions (**14**) or Fab alone (**15**), respectively, gave a similar response to the infliximab variant having only a bis-α-galactosylated glycan in the Fc region. These observations indicate that binding is not impacted by the presentation of the α-Gal epitope at either the Fc or Fab region.

Next, we assessed binding of IgE from sera of two patients with red meat allergy (serum-specific IgE against α-Gal: 89.9 kU/L and >100 kU/L). In contrast to the recombinant IgE, patient-derived IgE displayed a clear preference for α-Gal on the Fab-linked glycan. Structures bearing only Fc-linked α-Gal (**7** and **8**) bound significantly weaker than **14** and **15**, which have an α-Gal epitope on the Fab fragment (p < 0.001). Even variant **9**, fully α-galactosylated at the Fc, was outperformed by Fab-α-galactosylated variants **14** and **15** (p < 0.05). Both patients showed strikingly similar binding patterns, with only variants **7** and **14** differing significantly (p < 0.001 and p < 0.01, respectively). Thus, patient-derived IgE preferentially binds to α-Gal epitopes at Fab region.

### FcγR Binding Evaluation of Engineered Glycovariants

FcγRs are specialized receptor proteins on the surface of immune cells that bind to the Fc region of antibodies. They trigger immune responses such as phagocytosis, inflammation, and ADCC to destroy pathogens or infected cells. The conserved *N*-glycan at N297 of the CH2 domain of the Fc fragment of IgG modulates the binding of FcγRs.^3–6^ For example, the absence of core fucose of the *N*-glycan greatly increases the binding affinity to FcγRIIIa, thereby enhancing ADCC activity. Furthermore, sialic acid generally promotes anti-inflammatory properties, while lower levels are associated with pro-inflammatory states.

Glycovariants **4**-**12** were evaluated for binding to a panel of FcγRs, including FcγRI, FcγRIIa-131H, FcγRIIa-131R, FcγRIIb, FcγRIIIa-158V and -158F, and FcγRIIIb-NA1 and -NA2, by surface plasmon responses (SPR), and their affinities were compared to **3**, which is the A2F-glycan bearing reference glycoform (Table 1). Additional controls included unmodified infliximab (**1)** and the deglycosylated variant of infliximab (**2**). Compounds **14**-**17** are compared with the unmodified cetuximab (**13**).

**Table 1.**
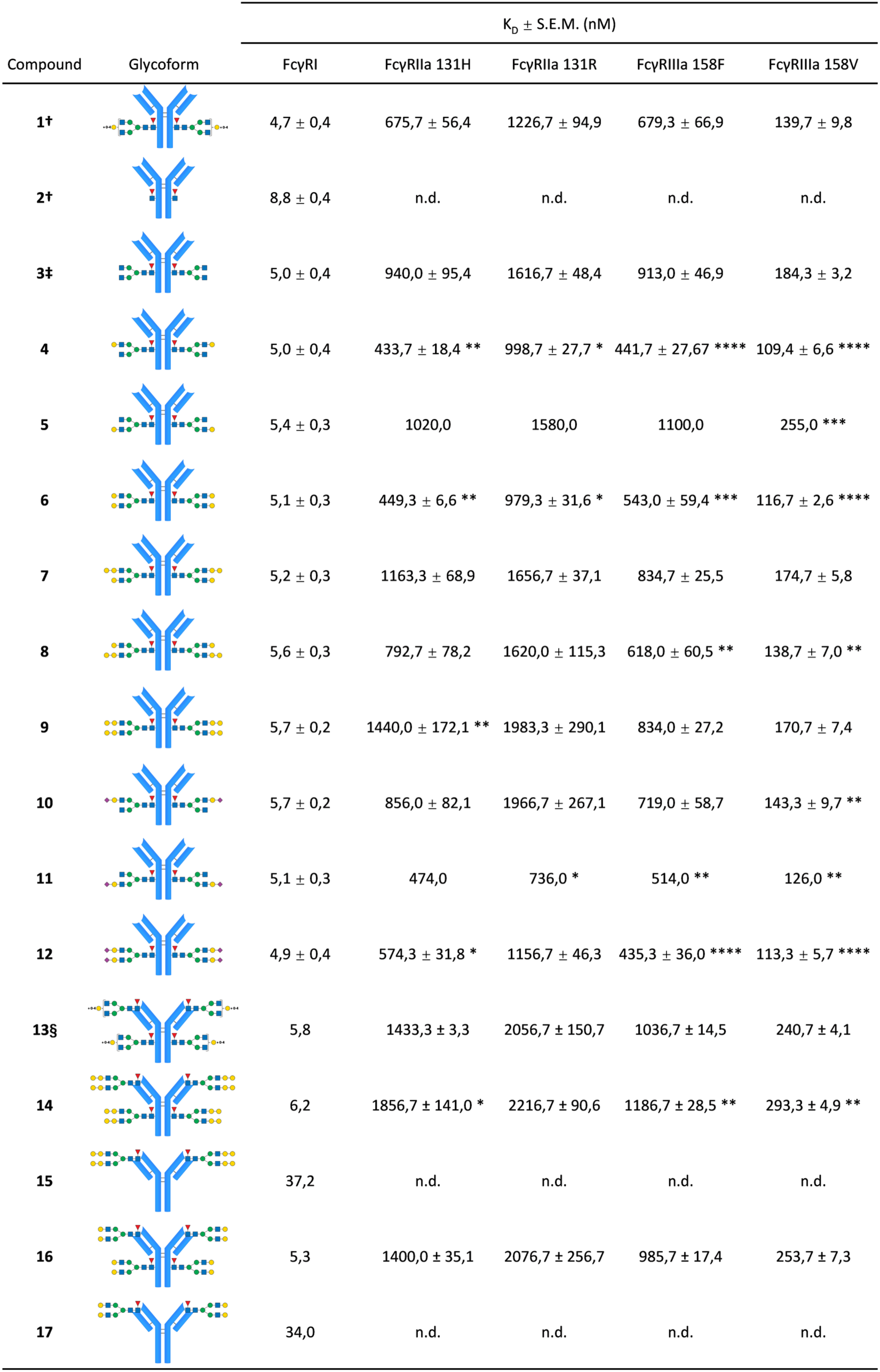
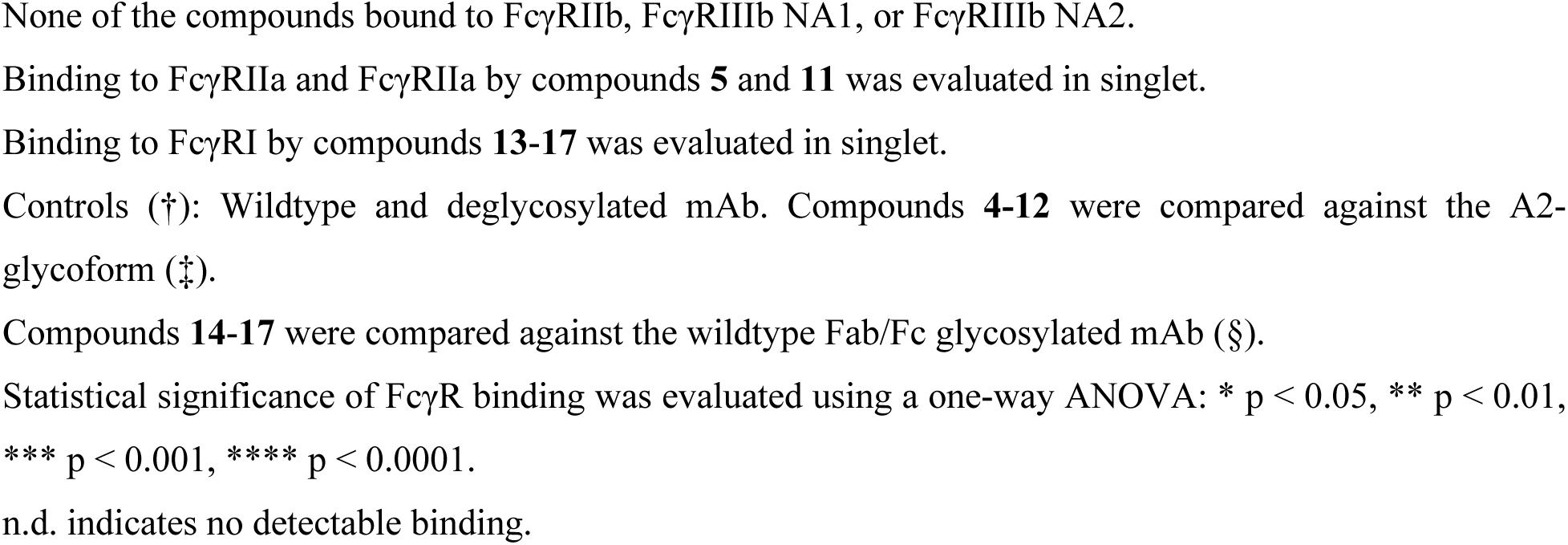
Determined KD values for the interaction between compounds 1-17 and FcγRs.

| Compound | Glycoform | $K_D \pm \text{S.E.M. (nM)}$ | | | | |
| --- | --- | --- | --- | --- | --- | --- |
|  |  | FcγRI | FcγRIIa 131H | FcγRIIa 131R | FcγRIIIa 158F | FcγRIIIa 158V |
| **1†** |  | 4,7 ± 0,4 | 675,7 ± 56,4 | 1226,7 ± 94,9 | 679,3 ± 66,9 | 139,7 ± 9,8 |
| **2†** |  | 8,8 ± 0,4 | n.d. | n.d. | n.d. | n.d. |
| **3‡** |  | 5,0 ± 0,4 | 940,0 ± 95,4 | 1616,7 ± 48,4 | 913,0 ± 46,9 | 184,3 ± 3,2 |
| **4** |  | 5,0 ± 0,4 | 433,7 ± 18,4 ** | 998,7 ± 27,7 * | 441,7 ± 27,67 **** | 109,4 ± 6,6 **** |
| **5** |  | 5,4 ± 0,3 | 1020,0 | 1580,0 | 1100,0 | 255,0 *** |
| **6** |  | 5,1 ± 0,3 | 449,3 ± 6,6 ** | 979,3 ± 31,6 * | 543,0 ± 59,4 *** | 116,7 ± 2,6 **** |
| **7** |  | 5,2 ± 0,3 | 1163,3 ± 68,9 | 1656,7 ± 37,1 | 834,7 ± 25,5 | 174,7 ± 5,8 |
| **8** |  | 5,6 ± 0,3 | 792,7 ± 78,2 | 1620,0 ± 115,3 | 618,0 ± 60,5 ** | 138,7 ± 7,0 ** |
| **9** |  | 5,7 ± 0,2 | 1440,0 ± 172,1 ** | 1983,3 ± 290,1 | 834,0 ± 27,2 | 170,7 ± 7,4 |
| **10** |  | 5,7 ± 0,2 | 856,0 ± 82,1 | 1966,7 ± 267,1 | 719,0 ± 58,7 | 143,3 ± 9,7 ** |
| **11** |  | 5,1 ± 0,3 | 474,0 | 736,0 * | 514,0 ** | 126,0 ** |
| **12** |  | 4,9 ± 0,4 | 574,3 ± 31,8 * | 1156,7 ± 46,3 | 435,3 ± 36,0 **** | 113,3 ± 5,7 **** |
| **13§** |  | 5,8 | 1433,3 ± 3,3 | 2056,7 ± 150,7 | 1036,7 ± 14,5 | 240,7 ± 4,1 |
| **14** |  | 6,2 | 1856,7 ± 141,0 * | 2216,7 ± 90,6 | 1186,7 ± 28,5 ** | 293,3 ± 4,9 ** |
| **15** |  | 37,2 | n.d. | n.d. | n.d. | n.d. |
| **16** |  | 5,3 | 1400,0 ± 35,1 | 2076,7 ± 256,7 | 985,7 ± 17,4 | 253,7 ± 7,3 |
| **17** |  | 34,0 | n.d. | n.d. | n.d. | n.d. |

None of the compounds bound to FcγRIIb, FcγRIIIb NA1, or FcγRIIIb NA2.
Binding to FcγRIIa and FcγRIIa by compounds **5** and **11** was evaluated in singlet.
Binding to FcγRI by compounds **13-17** was evaluated in singlet.
Controls (†): Wildtype and deglycosylated mAb. Compounds **4-12** were compared against the A2-glycoform (‡).
Compounds **14-17** were compared against the wildtype Fab/Fc glycosylated mAb (§).
Statistical significance of FcγR binding was evaluated using a one-way ANOVA: \* $p < 0.05$ , \*\* $p < 0.01$ , \*\*\* $p < 0.001$ , \*\*\*\* $p < 0.0001$ .
n.d. indicates no detectable binding.

Minimal or no engagement of the mAbs with FcγRIIb, FcγRIIIb-NA1 and -NA2 was observed (the corresponding sensorgrams are shown in Figure S39). High-affinity FcγRI bound all glycovariants with similar affinity, except deglycosylated variant **2**, which showed a small reduction in KD. Thus, Fc glycosylation is of minor importance for FcγRI binding. Deglycosylated **2** lost binding to all other FcγRs, indicating the importance of Fc glycosylation for these receptors. For the low-affinity receptors FcγRIIa and FcγRIIIa, comparison of **3**, **4**, **5**, and **6** revealed that galactosylation of the α1,6-arm increased binding, irrespective of the presence of a galactoside on the α1,3-antenna, whereas the galactoside on the α1,3-arm alone had no detectable effect. These observations agree with previous findings.^59,60^ This positive effect of ɑ1,6-antenna galactosylation on the affinity of mAb **4** was negated when this branch was capped by Neu5Ac or α-Gal, such as in mAbs **7** and **10**. Conversely, modification of the galactoside at the α1,3-antenna by α-Gal in mAb **8** (compared with **5**) resulted in modest improvement in binding to both FcγRIIIa allotypes. This effect was lost when the α1,6-antennae also carried an αGal cap, such as in **9**. A similar affinity increase was observed when the α1,3-arm was capped with Neu5Ac, however disialylation resulted in even stronger binding to FcγRIIIa. In addition to galactosylation, sialylation positively correlates with FcγRIIIa binding, while for FcγRIIa, sialylation of the α1,3-arm, the preferred branch of ST6Gal1, is sufficient for optimal binding.

Comparing α1,6-galactosylation of **4** with the α-galactosylated mAb **7**, binding to FcγRIIa 131H and FcγRIIIa 158F is significantly impacted by the presence of α-Gal, worsening binding by a factor of ∼2.5 and ∼2, respectively. This effect is not observed when β-galactosylation and α-Gal are present on the α1,3-branch, as in **5** and **8**. Bis-β-galactosylated and bis-α-galactosylated structures **6** and **9** show a similar pattern as **4** and **7**, indicating α-galactosylation of the top branch is the primary culprit.

The contribution of Fab-glycosylation to the affinity for the FcγRs appeared marginal. Solely the removal of the Fc glycan abrogated the binding to all FcγRs, below detection for the low-affinity receptors, while severely affecting binding to FcγRI (Table 1), for example in the case of **15** and **17** relative to **13**.

## CONCLUSIONS

In this study, we present a versatile methodology to precisely glycol-remodel the Fc glycan of mAbs. Previous research had indicated that asymmetrical structures, in which just a single branch of the biantennary glycan is modified, can modulate effector functions. For example, galactosylation of the α1,6-antennae was shown to benefit the binding of FcγRIIIa.^17,59–61^ Our generated glycoforms confirm the effect of galactosylation on FcγRIIIa binding. In addition, we found that sialylation of only the ɑ1,3-arm has superior binding to FcγRIIa. It is well established that sialylation has a profound role in instigating an anti-inflammatory effect, for instance in the case of treatment with IVIg. In contrast, galactosylation of the α1,3-antennae decreased binding to both FcγRIIa and FcγIIIa, consistent with previous reports.^17,59^ Although increased overall Fc galactosylation is generally reported to enhance FcγR binding and effector function, our arm-resolved data reveal that this relationship is site-specific: galactosylation of the α1,3-antenna alone decreases FcγRIIa/FcγRIIIa binding, opposite to the enhancing effect attributed to bulk galactosylation. This indicates that the two antennae of the Fc glycan can exert functionally opposing effects on FcγR engagement, an important consideration when evaluating antibody effector function in relation to glycosylation. Some reports indicate that full sialylation is not a requirement for improved IVIg-based therapy outcomes.^62,63^ Site-specific sialylation had arm-dependent effects on binding to the classical activating FcγRs: sialylation of the α1,3-antenna in **11** alone was sufficient to recapitulate most of the affinity gain seen with full sialylation of **12** for FcγRIIa (KD 474.0 and 736.0 nM for the 131H and 131R allotypes, respectively, versus 856.0 and 1966.7 nM for α1,6-arm sialylation (**10**) alone) and FcγRIIIa (KD 514.0 and 126.0 nM for the 158F and 158V allotypes, versus 719.0 and 143.3 nM for α1,6-arm sialylation alone). In contrast, minimal or no engagement with FcγRIIb suggests that anti-inflammatory effects previously attributed to IgG sialylation are not mediated through these inhibitory FcγRs. DC-SIGN engagement was not evaluated in this study; however, prior work has reported that the DC-SIGN sialylated Fc interaction is dispensable for the anti-inflammatory activity of sialylated IgG on human dendritic cells, leaving Siglecs as a plausible candidate lectin-type receptor mediating the anti-inflammatory effects of (mono-)sialylated glycans.^64–66^ The modularity of our approach provides an ideal platform to investigate the contribution of a single Neu5Ac per glycan either on the ɑ1,3-or the ɑ1,6-mannose arm.

Engineering asymmetrical α-galactosylated structures enabled us to determine the minimal requirement for IgE recognition, and apparently, at minimum, one α-Gal epitope in the Fc glycan, so as few as 2 α-galactosylated epitopes per mAb were enough to elicit IgE binding *in vitro* (as compound **7** barely makes it above the background as depicted in Figure 6A). A slight increase in IgE binding is seen when the α-Gal epitope is located at the opposing α1,6-branch, comparable to the improved binding to FcγRIIIa and C1q when only the α1,6-branch was galactosylated. However, the most striking enhancement occurred when both branches of the Fc glycan were α-galactosylated. In contrast to FcγRIIIa, where additional galactose beyond A2G1F(α1,6) only marginally improves affinity, IgE binding increased synergistically with α-galactosylation of the α1,3-arm. This difference likely reflects the mode of recognition: whereas FcγR engagement perturbs the conformation of the α1,6-arm indirectly, IgE interacts directly with the α-gal epitopes.^67^ We speculate that α-Gal on the α1,3-branch may increase the accessibility or favorable orientation of both terminal epitopes, thereby amplifying IgE affinity when the Fc glycan is fully α-galactosylated.

Using sera from patients with red meat allergy, we observed that Fab glycosylation plays a far more prominent role in IgE-mediated recognition than Fc-associated α-Gal. Rather than IgE binding independently to underlying structures (as our α-Gal-negative controls showed no baseline reactivity), this moderate response may reflect the structural constraints of the Fc region. Endogenous, polyclonal anti-α-Gal IgE may require an extended conformational epitope where terminal α-Gal is presented in a specific orientation alongside the underlying LacNAc or core mannose. The sterically restricted Fc cavity may prevent optimal docking of such IgE clones, whereas enhanced Fab accessibility facilitates higher avidity binding. Additionally, serum components could augment IgE-glycan interactions in ways that remain unclear.

Several studies have indicated that anti-α-Gal IgE is not the dominant driver of infliximab hypersensitivity. Some patients generate anti-drug IgE targeting peptide epitopes rather than against glycosidic epitopes,^68^ and red meat allergy sera were previously shown to bind cetuximab, but not infliximab, via α-Gal-specific IgE.^69^ In agreement, unmodified infliximab (mAb **1**) did not bind either recombinant or patient-derived IgE in our assays. However, comparing engineered non-α-galactosylated infliximab variants (mAbs **2**-**6**, **10**-**12**) with α-galactosylated counterparts (mAbs **7**-**9**) revealed clear α-Gal-dependent binding, showing that infliximab can elicit IgE engagement when α-gal is present, even though its native α-gal content is extremely low. This aligns with quantitative analyses showing that Remsima® and related biosimilars carry about 1-1.5% α-galactosylated glycans.^70^

Conversely, cetuximab shows high levels of Fab α-galactosylation: lectin arrays detect strong binding to α-Gal-recognizing lectins,^71^ and MS analyses report that up to 89% of Fab-glycans contain α-Gal.^51^ Its Fc region, nearly identical to infliximab’s, remains minimally α-galactosylated. These observations indicate that Fab-glycans are far more accessible substrates for GGTA1 than Fc glycans. Although GGTA1 readily modified both Fab and Fc regions during our *in vitro* glycoremodeling procedures, this does not reflect the constraints of cellular biosynthesis.

In conclusion, the enzymatic generation of asymmetrical mAb glycoforms provides a framework to dissect arm-specific contributions of defined glycan structures. Using this approach, we confirm the key role of α1,6-terminal galactose in FcγRIIIa binding and identify α1,3-specific sialylation as a determinant of FcγRIIa engagement. Furthermore, we demonstrate that α-galactosylated infliximab variants elicit α-gal-dependent IgE binding, with synergistic enhancement when both Fc glycan branches carry the α-gal epitope. These results highlight the importance of precise glycan topology in shaping immune recognition and point to the potential of arm-specific glycoengineering to unravel more functions of individual monosaccharides and epitopes within the mAb glycome.

## ASSOCIATED CONTENT

### Supporting Information

The Supporting Information containing experimental procedures, analytical data, additional figures and tables, and MS and NMR spectra (PDF) is available free of charge at https://pubs.acs.org/doi/

## AUTHOR INFORMATION

### Notes

The authors declare no competing financial interests.

## ACKNOWLEDGMENTS

The research was supported by Utrecht University (to G.B.), grants from NWO TA PPS fund (ENPPS.TA.019.008 to G.J.B. and A.C.K.) and the National Institute of Allergy and Infectious Diseases, National Institutes of Health (Award Number R01 AI165692 to G.J.B.).

