## Supplementary material for "Asymmetrical Glyco-Engineering of Monoclonal Antibodies Provides Insights into α-Gal Immunogenicity": SI

#### Table of contents


Glycan nomenclature ........ S4

Experimental procedures ........ S5

NMR settings ........ S5

Instrument settings mass spectrometry ........ S5

HILIC-MS analyses of glycans **27-30** ........ S5

Preparation of glycan **27** ........ S6

Preparation of glycan **28** ........ S7

Preparation of glycan **29** ........ S8

Preparation of glycan **30** ........ S9

Ion-mobility MS characterization of released glycans from mAb **4** ........ S10

Glycopeptide analysis of mAb **4** ........ S10

Expression of Endo-S2, BgaA, StrH, PNGase-F, Pd26ST, and Psp26ST A366G/A235M in *E. coli* ........ S10

Expression of GnT-I, B4GalT1, ST6Gal1 and mmGGTA1 in HEK293F suspension cells ........ S11

General procedure for Ni-NTA purification of *E. coli* and HEK produced enzymes ........ S12

General procedure for STREP-tag purification of *E. coli* expressed enzymes and mmGGTA1 ........ S12

General procedure for SEC purification of *E. coli* and HEK produced enzymes ........ S12

General procedure for StrH glycan remodeling of an intact mAb ........ S12

General procedure for terminal GlcNAc installation on an intact mAb ........ S12

General procedure for terminal β1,4-galactose installation on an intact mAb ........ S13

General procedure for terminal α1,3-galactose installation on an intact mAb ........ S13

General procedure for Neu5Ac installation on an intact mAb ........ S13

General procedure for α-Gal removal on an intact mAb ........ S13

Deglycosylation of mAbs with Endo-S2 WT or PNGase-F ........ S14

Protein A purification of mAbs ........ S14

LC-ESI-MS analysis of glycoprotein fragments ........ S14

IgE binding experiments ........ S14

SPR binding studies ........ S15

References ........ S16

Figures and Tables ........ S17

Figure S1. LC-MS and NMR analysis of glycan **27** ........ S17

Figure S2. LC-MS and NMR analysis of glycan **28** ........ S21

Figure S3. LC-MS and NMR analysis of glycan **29** ........ S26

Figure S4. LC-MS and NMR analysis of glycan **30** ........ S31

Figure S5. NMR linkage analysis of glycans **28** and **30** ........ S36

Figure S6. PGC-MS chromatogram of released glycan mAb **4** ........ S37

Figure S7. Glycopeptide analysis of mAb **4** .................... S38
Table S1. Detected glycopeptides of mAb **4** tryptic digest .................... S39
Figure S8. Sialylation optimization using mAb **4** .................... S40
Figure S9. Sialylation optimization using mAb **19** .................... S41
Figure S10. Complete enzymatical, asymmetrical glycoremodeling route .................... S42
Figure S11. Symmetrical glycoremodeling routes of infliximab and cetuximab .................... S43
Table S2. Theoretical masses versus observed masses infliximab .................... S44
Table S3. Theoretical masses versus observed masses cetuximab .................... S45
Figure S12. Deconvoluted MS spectrum mAb **1**.................... S47
Figure S13. Deconvoluted MS spectrum mAb **2**.................... S48
Figure S14. Deconvoluted MS spectrum mAb **3**.................... S49
Figure S15. Deconvoluted MS spectrum mAb **4**.................... S50
Figure S16. Deconvoluted MS spectrum mAb **5**.................... S51
Figure S17. Deconvoluted MS spectrum mAb **6**.................... S52
Figure S18. Deconvoluted MS spectrum mAb **7**.................... S53
Figure S19. Deconvoluted MS spectrum mAb **8**.................... S54
Figure S20. Deconvoluted MS spectrum mAb **9**.................... S55
Figure S21. Deconvoluted MS spectrum mAb **10**.................... S56
Figure S22. Deconvoluted MS spectrum mAb **11**.................... S57
Figure S23. Deconvoluted MS spectrum mAb **12**.................... S58
Figure S24. Deconvoluted MS spectrum mAb **13**.................... S59
Figure S25. Deconvoluted MS spectrum mAb **14**.................... S61
Figure S26. Deconvoluted MS spectrum mAb **15**.................... S63
Figure S27. Deconvoluted MS spectrum mAb **16**.................... S65
Figure S28. Deconvoluted MS spectrum mAb **17**.................... S67
Figure S29. Deconvoluted MS spectrum intermediate mAb **18**.................... S69
Figure S30. Deconvoluted MS spectrum intermediate mAb **19**.................... S70
Figure S31. Deconvoluted MS spectrum intermediate mAb **20**.................... S71
Figure S32. Deconvoluted MS spectrum intermediate mAb **21**.................... S72
Figure S33. Deconvoluted MS spectrum intermediate mAb **22**.................... S73
Figure S34. Deconvoluted MS spectrum intermediate mAb **23**.................... S74
Figure S35. Deconvoluted MS spectrum intermediate mAb **24**.................... S75
Figure S36. Deconvoluted MS spectrum intermediate mAb **25**.................... S76
Figure S37. Deconvoluted MS spectrum intermediate mAb **26**.................... S77
Figure S38. SDS-PAGE gel – Coomassie stain of mAbs **1-17**.................... S78
Figure S39. SPR FcγR binding sensorgrams. .................... S79

#### Glycan nomenclature

| Glycan | Assigned name(s) |
| --- | --- |
| 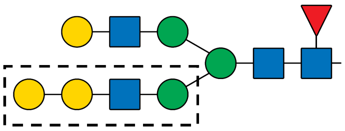 | α1,3-mannose arm, α1,3-arm, arm processed by **GnT-I**. Also referred to as bottom branch. Example structure. |
| 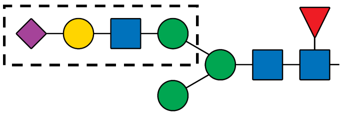 | α1,6-mannose arm, α1,6-arm, arm processed by **GnT-II**. Also referred to as the top branch. Example structure. |
| 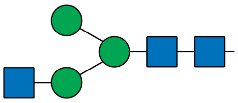 | A1(α1,3)-glycan |
| 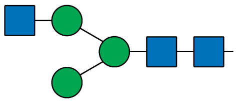 | A1(α1,6)-glycan |
| 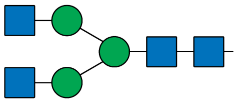 | A2-glycan |
| 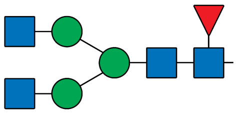 | A2F-glycan |
| 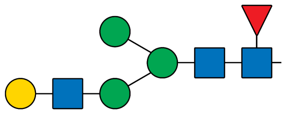 | A1G1F(α1,3)-glycan |
| 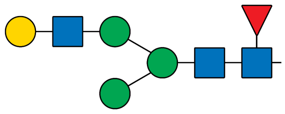 | A1G1F(α1,6)-glycan |
| 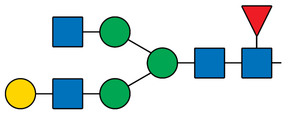 | A2G1F(α1,3)-glycan |
| 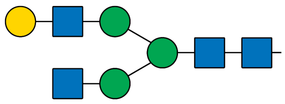 | A2G1F(α1,6)-glycan |

### Experimental procedures

#### NMR settings

Resolution-enhanced 1D/2D $^1H$ NMR (600 MHz) and $^{13}C$ NMR (150 MHz) spectra were recorded in $D_2O$ using a Bruker Avance Neo spectrometer equipped with a TCI Prodigy CryoProbe™ (Bruker, Billerica, MA, USA) at a probe temperature of 298 K. Prior to analysis, samples were exchanged twice in $D_2O$ (99.9 atom % D, Cambridge Isotope Laboratories, Inc., Andover, MA) with intermediate lyophilization, and then dissolved in 0.5 mL $D_2O$. Chemical shifts ($\delta$) are expressed in ppm by reference to internal acetone ($\delta$ 2.225 for $^1H$ and 31.07 for $^{13}C$). Suppression of the HOD signal was achieved by applying a WEFT pulse sequence for 1D experiments and by a pre-saturation of 1 s during the relaxation delay in 2D experiments. 2D TOCSY spectra were recorded using an MLEV-17 mixing sequence with spin-lock times of 40–150 ms. 2D NOESY experiments were performed with a mixing time of 300 ms. Natural abundance 2D $^1H$-$^{13}C$ HSQC experiments were recorded without decoupling during acquisition of the $^1H$ FID.

#### Instrument settings mass spectrometry

For IMS analysis of the released glycan of **4**, an Agilent Technologies 6560 Ion mobility Q-TOF LC/MS system was used. For analysis of the 2-AA labelled glycans the MS source parameters were; nitrogen drying gas heated to 300 °C at 8 L/min and sheath gas heated to 350 °C at 11 L/min, nebulizer pressure of 40 psig, capillary voltage of 3500 V and nozzle voltage of 1000 V. The IMS measurements were conducted using an IM transient rate of 16 transients/frame, a trap fill time of 3900 µs, a trap release time of 250 µs, a drift tube entrance voltage of -1700, a drift tube exit voltage of -250 V, nitrogen as drift tube gas type, a drift tube pressure of 3.95 Torr, a trap funnel pressure of 3.80 Torr and a multiplexing pulsing sequence length of 4 bit. The IM-MS was used in negative polarity mode with a mass range of m/z 100-1700. The data was demultiplexed using PNNL Preprocessor v4.0 (Pacific Northwest National Laboratory, Richland WA) using an interpolation of 3 drift bins and a 5 point moving average smoothing in the IM dimension. Features were identified with Masshunter IMS browser software v10.0 (Agilent Technologies) using an unbiased isotope model. High-resolution arrival-time distributions (ATDs) were obtained using Agilent Technologies HRdm v2.0 software with an *m/z* width multiplier of 12, saturation check of 0.40 and an IF multiplier of 1.125 with SSS and Post QC enabled.

For analysis of protein fragments only the QTOF functionality of the Agilent Technologies 6560 Ion mobility Q-TOF LC/MS was used, the Q-TOF source settings were optimized for high-resolution protein characterization; nitrogen gas temperature was 350 °C with a flow of 8 L/min, the pressure of the nebulizer gas was 45 psig, sheath gas of 400 °C with a flow of 11 L/min was pumped into the chamber. The fragmentor was set at 380 V, the capillary voltage was 5500 V and the nozzle voltage was 2000 V.

#### HILIC-MS analyses of glycans 27-30

For analysis of glycans **27**-**30**, a Shimadzu LC-ESI-IT-TOF was used in combination with a Waters XBridge BEH, Amide Column (3.5 µm, 2.1 x 150 mm). Over a linear gradient of 80-50% B vs A in 18 min at 25 °C (A: 10 mM $NH_4HCOO$ in ultrapure water, pH3.5, B: ACN) with a flow rate of 0.2 mL/min, glycans and byproducts could be separated. After the 18-min elution phase, a washing step consisting of 6 min at 25% B and subsequently, an equilibration phase of 6 min at 80% B followed.

**Preparation of glycan 27**

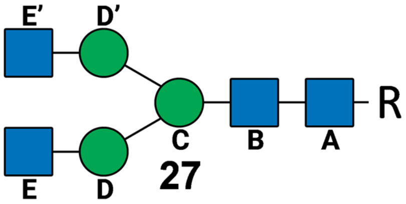


Glycan **27** with a NAP-protected (R = naphtylmethyl) asparagine was prepared according to previous literature procedures.[1] First, SGP was extracted as previously described.[2] Briefly, SGP was truncated to a single asparagine using Pronase from *S. griseus* and further desialylated and degalactosylated using sialidase from *C. perfringens* and galactosidase from *A. niger* to yield the glycosyl asparagine. To a solution of the glycosyl asparagine (500 mg, 3.5 mM) in DMF was added 2-napthylmethyl-*p*-nitrophenol carbonate (4 eq.) and $NEt_3$ (8 eq.) and stirred overnight at 45 °C. Reaction completeness was determined by LC-MS. Once all starting material was consumed, the solvent was evaporated *in vacuo* and the residue was reconstituted in water. The slurry was filtered through a Buchner filter frit and rinsed with water. The filtrate was further concentrated in vacuo and lyophilized overnight. The crude material was further purified by a C18 silica plug, eluting the product with 20% aq. ACN. Lastly, Bio-Gel P2 Extra-Fine size-exclusion chromatography was used to further purify the material, yielding the naphthyl protected glycosyl asparagine, Glycan **27** as a fluffy white solid (253 mg, 45.8%). NMR spectrum is provided in Figure S1.

| Residue | H1<br>C1 | H2<br>C2 | H3<br>C3 | H4<br>C4 | H5<br>C5 | H6a;H6b<br>C6 | NHAc |
|---|---|---|---|---|---|---|---|
| **A** | 4.83<br>78.0 | 3.66<br>53.6 | 3.53<br>72.8 | 3.45<br>78.4 | 3.23<br>76.0 | 3.54; 3.41<br>59.7 | 1.79<br>22.0 |
| **B** | 4.46<br>101.1 | 3.71<br>54.8 | 3.66<br>72.0 | 3.64<br>79.3 | 3.50<br>74.4 | 3.83; 3.53<br>61.6 | 2.00<br>22.2 |
| **C** | 4.68<br>100.4 | 4.16<br>70.2 | 3.68<br>80.5 | 3.72<br>65.4 | 3.54<br>74.3 | 3.88; 3.69<br>65.6 | - |
| **D** | 5.03<br>99.5 | 4.10<br>76.4 | 3.81<br>69.4 | 3.41<br>67.3 | 3.66<br>73.5 | 3.77; 3.64<br>59.9 | - |
| **D'** | 4.84<br>96.9 | 4.03<br>76.3 | 3.81<br>69.4 | 3.41<br>67.3 | 3.68<br>72.0 | 3.77; 3.68<br>60.0 | - |
| **E'** | 4.46<br>99.6 | 3.62<br>55.2 | 3.47<br>73.3 | 3.37<br>69.8 | 3.35<br>75.8 | 3.83; 3.67<br>60.5 | 1.95<br>22.2 |
| **E** | 4.46<br>99.6 | 3.62<br>55.2 | 3.47<br>73.3 | 3.37<br>69.8 | 3.35<br>75.8 | 3.83; 3.67<br>60.5 | 1.95<br>22.2 |

**Preparation of glycan 28**

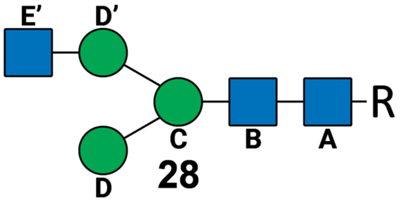


The A2-glycan **27** (20 mg, 12.4 µmol) was dissolved in 1.2 mL of 100 mM NaOAc buffer pH 6.0, supplemented with 5 mM $CaCl_2$. Next, StrH (0.1% wt/wt of A2-glycan) was added to the solution and left to incubate overnight at 37 °C. Reaction completeness was confirmed using LC-MS (Figure S2). The reaction was heated at 100 °C in a heat-block for 5 min before centrifugation to remove precipitated protein. The supernatant was loaded onto a C18 silica plug and rinsed with water and eluted with 20% ACN and lyophilized. Glycan **28** was obtained as a fluffy white solid (14.1 mg, 81%). NMR spectrum is provided in Figure S2.

| Residue | H1<br>C1 | H2<br>C2 | H3<br>C3 | H4<br>C4 | H5<br>C5 | H6a;H6b<br>C6 | NHAc |
|---|---|---|---|---|---|---|---|
| **A** | 4.84<br>78.06 | 3.68<br>53.5 | 3.47<br>72.8 | 3.26<br>78.4 | 3.26<br>76.1 | 3.56; 3.43<br>59.7 | 1.79<br>22.0 |
| **B** | 4.47<br>101.1 | 3.71<br>54.8 | 3.66<br>72.0 | 3.64<br>79.5 | 3.50<br>74.4 | 3.83; 3.54<br>61.6 | 2.00<br>22.2 |
| **C** | 4.68<br>100.3 | 4.17<br>70.2 | 3.68<br>80.5 | 3.72<br>65.4 | 3.54<br>74.3 | 3.89; 3.70<br>65.8 | - |
| **D** | 5.02<br>102.4 | 3.99<br>69.9 | 3.81<br>69.4 | 3.55<br>66.8 | 3.73<br>73.4 | 3.78; 3,65<br>59.9 | - |
| **D’** | 4.84<br>96.9 | 4.03<br>76.2 | 3.81<br>70.3 | 3.41<br>67.3 | 3.67<br>71.9 | 3.83; 3.66<br>61.0 | - |
| **E’** | 4.47<br>99.6 | 3.63<br>55.2 | 3.46<br>73.4 | 3.39<br>69.8 | 3.35<br>75.8 | 3.84; 3.68<br>60.6 | 1.97<br>22.2 |

**Preparation of glycan 29**

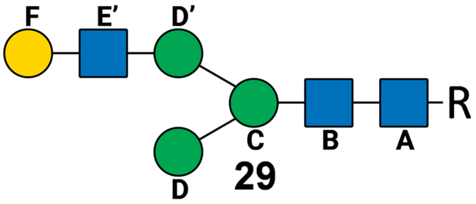


For preparation of glycan **29** (R = naphtylmethyl), 14.1 mg of glycan **28** (10 µmol) was dissolved in 1 mL of 100 mM TRIS pH 7.5, supplemented with 10 mM $MnCl_2$, 15 mM UDP-Galactose (MU06699, Carbosynth) and B4GalT1 (*H. sapiens*) (1% wt/wt glycan 2) and 20 U of calf intestine alkaline phosphatase (CIAP) (M1821, Promega) and was incubated overnight at 37 °C with gently shaking. Reaction completeness was confirmed using LC-MS (Figure S3). The reaction was heated at 100 °C in a heat-block for 5 min before centrifugation to remove precipitated protein. The supernatant was loaded onto a C18 silica plug and rinsed with water and eluted with 20% ACN and lyophilized. Glycan **29** was obtained as a fluffy white solid (9.6 mg, 61%). NMR spectrum is provided in Figure S3.

| Residue | H1<br>C1 | H2<br>C2 | H3<br>C3 | H4<br>C4 | H5<br>C5 | H6a;H6b<br>C6 | NHAc |
|---|---|---|---|---|---|---|---|
| **A** | 4.83<br>78.0 | 3.67<br>53.6 | 3.54<br>72.9 | 3.47<br>78.4 | 3.23<br>76.0 | 3.55; 3.42<br>59.7 | 1.79<br>22.0 |
| **B** | 4.47<br>101.1 | 3.71<br>54.8 | 3.66<br>72.0 | 3.64<br>79.3 | 3.49<br>74.5 | 3.83; 3.53<br>61.6 | 2.00<br>22.2 |
| **C** | 4.68<br>100.3 | 4.17<br>70.2 | 3.68<br>80.5 | 3.72<br>65.4 | 3.54<br>74.3 | 3.89; 3.70<br>65.8 | - |
| **D** | 5.02<br>102.4 | 3.99<br>70.0 | 3.81<br>69.4 | 3.55<br>66.8 | 3.73<br>73.4 | 3.78; 3,65<br>59.9 | - |
| **D'** | 4.84<br>96.9 | 4.03<br>76.2 | 3.81<br>70.3 | 3.41<br>67.3 | 3.54<br>72.6 | 3.83; 3.66<br>61.0 | - |
| **E'** | 4.50<br>99.5 | 3.68<br>54.7 | 3.45<br>72.8 | 3.65<br>78.5 | 3.35<br>69.8 | 3.84; 3.68<br>60.6 | 1.95<br>22.2 |
| **F** | 4.39<br>103.0 | 3.46<br>70.9 | 3.59<br>72.4 | 3.84<br>68.5 | 3.65<br>75.3 | 3.90; 3.77<br>59.9 | |

##### Preparation of glycan 30

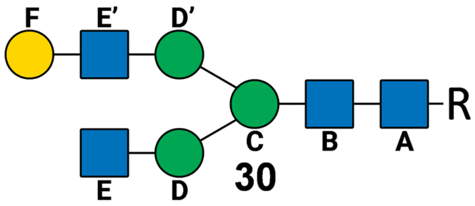


**30**

For preparation of glycan **30** (R = naphtylmethyl), 8.3 mg of glycan **29** (5.3 μmol) was dissolved in 530 μL of 100 mM MES pH 6.5, supplemented with 10 mM $MnCl_2$, 15 mM UDP-Galactose (MU06699, Carbosynth) and GnT-I (*H. sapiens*) (1% wt/wt glycan **29**) and 20 U of CIAP (M1821, Promega) and was incubated overnight at 37 °C with gently shaking. Reaction completeness was confirmed using LC-MS (Figure S4). The reaction was heated at 100 °C in a heat-block for 5 min before centrifugation to remove precipitated protein. The supernatant was loaded onto a C18 silica plug and rinsed with water and eluted with 20% ACN and lyophilized. Glycan **30** was obtained as a fluffy white solid (6.8 mg, 72.3%). NMR spectrum is provided in Figure S4.

| Residue | H1<br>C1 | H2<br>C2 | H3<br>C3 | H4<br>C4 | H5<br>C5 | H6a;H6b<br>C6 | NHAc |
|---|---|---|---|---|---|---|---|
| **A** | 4.81<br>78.0 | 3.65<br>53.6 | 3.52<br>72.9 | 3.45<br>78.4 | 3.21<br>76.0 | 3.53; 3.40<br>59.7 | 1.79<br>22.0 |
| **B** | 4.45<br>101.1 | 3.69<br>54.8 | 3.64<br>72.0 | 3.63<br>79.3 | 3.46<br>74.6 | 3.81; 3.52<br>61.6 | 2.00<br>22.2 |
| **C** | 4.66<br>100.4 | 4.15<br>70.2 | 3.66<br>80.5 | 3.72<br>65.4 | 3.52<br>74.2 | 3.87; 3.69<br>65.6 | - |
| **D** | 5.02<br>99.6 | 4.09<br>76.4 | 3.79<br>69.4 | 3.40<br>67.3 | 3.65<br>73.5 | 3.88; 3.64<br>59.9 | - |
| **D'** | 4.83<br>96.9 | 4.01<br>76.3 | 3.79<br>69.4 | 3.40<br>67.3 | 3.52<br>72.8 | 3.80; 3.65<br>60.6 | - |
| **E** | 4.48<br>99.5 | 3.65<br>54.7 | 3.46<br>73.2 | 3.63<br>78.5 | 3.35<br>69.8 | 3.84; 3.68<br>60.6 | 1.95<br>22.2 |
| **E'** | 4.45<br>99.6 | 3.60<br>55.2 | 3.46<br>73.2 | 3.36<br>69.8 | 3.34<br>75.8 | 3.81; 3.66<br>60.5 | 1.95<br>22.2 |
| **F** | 4.37<br>103.0 | 3.44<br>70.9 | 3.57<br>72.4 | 3.83<br>68.5 | 3.63<br>75.3 | 3.76; 3.74<br>59.9 | - |

**Ion-mobility MS characterization of released glycans from mAb 4**

To verify the glycosylation homogeneity of antibody **4** by ion-mobility-mass spectrometry (IM-MS), 100 µg of the StrH-, B4GalT1-, and GnT-I-treated antibody was incubated with 1 µg of Endo-S2 WT in 150 µL of 50 mM NaOAc buffer (pH 4.5) at 37 °C for 1 h..To separate the proteinaceous content from the released glycans a Vivaspin 500 10 kDa MWCO spin filter (VS0102, Sartorius) was used. To label the released glycans, 120 uL of the spin-filtrate containing the released glycans was derivatized by adding 20 µL undiluted acetic acid, followed by the addition of 40 µL labeling solution consisting of anthranilic acid (2-AA; 30 mg/mL; A89855, Sigma-Aldrich) and sodium cyanoborohydride (75 mg/mL; 156159, Sigma-Aldrich) in DMSO. The mixture was incubated at 68 °C for 2 h and then 1 mL ultrapure water was added. Next, the mixture was purified by solid phase extraction (SPE) using a HyperSep Hypercarb SPE Cartridge (60106-304, Thermo Scientific). Briefly, the SPE cartridge was conditioned with 1 mL acetonitrile (ACN) and washed with 1 mL water. The sample (1180 µL) was loaded on the cartridge and washed with 1 mL 0.05% trifluoro acetic acid (TFA), followed by 1 mL ACN/water/TFA (5%/94.95%/0.05%). Glycans were eluted with 1 mL ACN/0.1% TFA (50%/50%). The eluent was evaporated under a stream of nitrogen and redissolved in 20 µL 90% water/10% ACN of which 5 µL was injected onto a Hypercarb PGC 150 x 4.6 mm (3 µm) column (35003-154630, Thermo Scientific) using an Agilent 1290 Infinity LC, run isocratic for 1 min, using 10% B, then run over a linear gradient of 10-95% B vs A in 39 min at 25 °C (A: 5 mM $NH_4HCOO$ in ultrapure water, B: ACN) with a flow rate of 0.5 mL/min. The detector was an Agilent Technologies 6560 Ion mobility Q-TOF LC/MS system, for IMS settings and processing of data see above.

**Glycopeptide analysis of mAb 4**

Enzymatic digestion was performed by incubating compound **4** (200 µg) with trypsin (10 µg; T4799, Sigma) in 200 µL of 50 mM TRIS buffer (pH 7.5). The reaction mixture was maintained at 37 °C for 18 h under gentle agitation. Next, the sample was cleaned up using an C18 SPE cartridge (WAT054955, Waters). Briefly, 800 µL of 0.1% formic acid (FA) was added to the sample. The column was conditioned with 3 mL methanol/water/FA (90%/9.9%/0.1%) and then washed with 3 mL 0.1% FA in water. Then the sample was loaded and let pass through the column by gravity flow. The column was then washed with 3 mL methanol/water/FA (5%/94.9%/0.1%). The sample was eluted in 500 µL ACN/water (50%/50%) and lyophilized and reconstituted in 20 µL ultrapure water of which 5 µL was injected onto an AdvanceBio Glycan Mapping column, 2.1 × 150 mm, 2.7 µm, (683775913, Agilent) using an Agilent 1290 Infinity LC, run over a linear gradient of 90-50% B vs A in 60 min at 25 °C (A: 0.1% FA in ultrapure water, B: ACN) with a flow rate of 0.3 mL/min. Figure S7 contains the results of the glycopeptide analysis of mAb 4. The detector was an Agilent Technologies 6560 Ion mobility Q-TOF LC/MS system, using the same MS source parameters as for the IMS measurements.

**Expression of Endo-S2, BgaA, StrH, PNGase-F, Pd26ST, and Psp26ST A366G/A235M in *E. coli***

Wildtype (WT) endoglycosidase Endo-S2 WT[3] from *S. pyogenes* (AA38-843) (GenBank: AGU16855.1) was cloned in a custom vector based on pET11a, with a N-terminal STREP-tag (WSHPQFEK), a superfolder GFP (GenBank: AAA27721.1 + S30R/Y39N/N105T/ Y145F/371V/A206V) and a spacer (GGGSGGGSGGS), and a C-terminal HIS-tag (HHHHHHHH). The galactosidase BgaA (AA2-2221) (GenBank: AAK99369.1) from *S. pneumoniae* was cloned in a custom vector based on pET11a with a N-terminal STREP-tag (WSHPQFEK), a mCherry (Uniprot D1MPT3 + N8D/K199N/T200V/D201N) and a spacer (GGGSGGGSGGS), and a C-terminal HIS-tag (HHHHHHHH). The GH20-2 domain of StrH from S. pneumoniae (AA616-977) (Genbank: AAK98861.1) with a C-terminal HIS-tag (HHHHHHHH) and Twin-STREP-tag (WSHPQFEKGGGSGGGSGGSSAWSHPQFEKG) was subcloned in vector pMAL-15x-H (Genscript) (AvaI, EcoRI). PNGase-F was prepared using the vector pOPH6 (40315, Addgene) and the periplasmic enzyme was purified according the previously described protocol.[4] Sialyltransferase Pd26ST WT[5] from *P. damsela* JT0160 (AA15-497) (GenBank: BAA25316.1) with a C-terminal HIS-tag (HHHHHHHH) and Twin-STREP-tag (WSHPQFEKGGGSGGGSGGSSAWSHPQFEKG) was subcloned in vector pMAL-15x-H (Genscript) (AvaI, EcoRI). Sialyltransferase Psp26ST A366G/A235M (AA111-511) (GenBank: BAF92026.1) from *P.sp.* JT-ISH-224 was subcloned in pET52b(+) (SalI/SacI).[6] BL21(DE3) (14527H, New England Biolabs) cells were transformed with each of the individual vectors, and plated on a 2xYT agar (BP97432, Fisher Bioreagents) plate with ampicillin (100 µg/mL) (14417, Cayman Chemical). On the next day, a colony was picked and expanded to a cell culture volume of 500 mL ampicillin containing (100 µg/mL) 2xYT medium (X966, Carl Roth) at 37 °C. The cells were grown for 3-5 h, until the enzyme production was induced at OD600 = 0.6 with isopropyl β-D-1-thiogalactopyranoside (IPTG) (R0393, Thermo Scientific) at a final concentration of 1 mM. After induction, the cells were cultured overnight at 20 °C. Then, the cells were pelleted at 3000 x *g*, resuspended in lysis buffer (TBS (Tris 25 mM, NaCl 150 mM, pH 7.5) with 1 mg/mL lysozyme (62971, Sigma-Aldrich) and 0.1% triton X-100 (T8787, Sigma-Aldrich)) at 5% of the original culture volume. The resuspended cells were incubated at RT for 1 h and then sonicated. Sonication was performed using a Bandelin Sonopuls HD2200, with a MS73 probe, at an amplitude of 50%, 3 times 10 seconds on, 10 seconds off. Supernatant was obtained by removal of cell debris by centrifugation at 10000 x *g*. The supernatant with the protein of interest was further purified by affinity chromatography and size-exclusion chromatography.

**Expression of GnT-I, B4GalT1, ST6Gal1 and mmGGTA1 in HEK293F suspension cells**

The luminal portion of the enzyme human GnT-1 (AA30-445) (GenBank: AAA52563.1), human B4GalT1 (45-398) (GenBank: AAH45773.1) and human ST6Gal1 (AA27-406) (GenBank: AAH31476.1) were cloned into a pGen2-vector resulting in a fusion protein of a N-terminal HIS-tag (HHHHHHHH), a superfolder GFP, a TEV protease site (ENLYFQG) and the enzyme of interest as previously described.[7] The luminal portion of the α1,3 sialyltransferase[8] from *M. musculus* (AA61-394) (GenBank: AAA37657.1) with N-terminal signal peptide from Siglec15 (MEKSIWLLACLAWVLPTGS) and with a HIS-tag (HHHHHHHH), C-terminal TEV protease site (ENLYFQS), and a Twin-STREP-tag (WSHPQFEKGGGSGGGSGGSSAWSHPQFEKG) was subcloned in pcDNA3.1+C-eGFP (Genscript) (NheI, EcoRV).

The enzymes GnT-I, B4GalT1, ST6Gal1 and mmGGTA1 were expressed through transient transfection in wild type HEK293F cells (Freestyle 293F). Briefly, cells were maintained in suspension culture at 1-3 x $10^6$ cells/mL in a humidified $CO_2$ shaker incubator (37 °C, 150 rpm). Transient transfection was performed at a cell density of 3-3.5 x $10^6$ cells/mL in HEK-TF medium (861-0001, Xell) supplemented with GlutaMAX (35050061, Gibco) and Pen-Strep (15070063, Gibco) using predissolved, sterile, transfection grade, linear polyethylenimine (PEI) 25000 g/mol (23966, Polysciences), at a concentration of 9 µg/mL and a vector concentration of 4 µg/mL by directly, dropwise addition of the solutions to the culture medium. The day after the transfection the suspension culture was diluted 1:1 with HEK-TF medium. Right after that, predissolved, sterile valproic acid (VPA) (P4542, Sigma-Aldrich) to a final concentration of 2.2 mM was added. Feed HEK-FS (871-0001, Xell) supplemented with GlutaMAX and Pen-Strep was added at day 1, 3, and 4 post-transfection at 5%, 7.5%, and 10% of the total culture volume respectively. On day 6 the supernatant was collected by centrifugation at 3000 *x g* at 4 °C for 20 min. After centrifugation, the supernatant was filtered over a 3 µm acetate filter to remove any residual cells and debris. The HEK293F cell-free supernatant containing the enzyme of interest was purified using a two-step purification procedure, first by Ni-NTA purification, followed by size-exclusion chromatography.

**General procedure for Ni-NTA purification of *E. coli* and HEK produced enzymes**

The enzymes Endo-S2, BgaA, StrH, PNGase-F, Pd26ST, Psp26ST A366G/A235M, GnT-I, B4GalT1, ST6Gal1 and mmGGTA1 were purified using Ni-NTA affinity chromatography. The clarified supernatant was loaded onto a 4 mL gravity flow Ni-NTA column (17-5318-01, GE healthcare), capable of binding the HIS-tagged protein. A standard buffer consisting of 50 mM TRIS-HCl and 250 mM NaCl pH 8 was made. Imidazole was added to the standard buffer at concentrations of 20 mM, 50 mM or 250 mM to make wash 1, wash 2 and elution buffer (re-adjusted to pH 8 if needed) respectively. Once washed with 10 column volumes (CV) of wash 1, 10 CV of wash 2, the enzyme of interest was eluted in 3 CV of elution buffer.

**General procedure for STREP-tag purification of *E. coli* expressed enzymes and mmGGTA1**

Through the STREP-tag the enzymes Endo-S2, BgaA, StrH, Pd26ST, Psp26ST A366G/A235M and mmGGTA1 were purified using Strep-Tactin Superflow resin (2-1206-025, IBA Lifesciences). The eluate from the Ni-NTA column was directly loaded onto a 4 mL gravity flow Strep-Tactin column. Elution buffer was prepared by addition of 2.5 mM desthiobiotin (D1411, Sigma-Aldrich) to wash buffer (100 mM TRIS, 150 mM NaCl, 1 mM EDTA, pH 8). Once the sample was loaded, unbound material was removed from the resin by washing with 10 CV of wash buffer, followed by elution of the protein of interest with 3-5 CV of elution buffer. Fractions of 2 mL were collected, pure fractions were pooled, concentrated and stored.

**General procedure for SEC purification of *E. coli* and HEK produced enzymes**

Size-exclusion chromatography (SEC) was used to further purify the enzymes Endo-S2, BgaA, StrH, PNGase-F, Pd26ST, Psp26ST A366G/A235M, GnT-I, B4GalT1, ST6Gal1 and mmGGTA1 after Ni-NTA and/or the STREP-tag purification. The eluate from either the Ni-NTA or the STREP-tag purification was concentrated to 10-30 mg/mL using a Vivaspin 6 filter 10 kDa MWCO (VS0602, Sartorius) and further purified on a TBS (25 mM TRIS, 150 mM NaCl, pH 7.5) equilibrated Superdex 200 Increase 10/300 GL column (28990944, GE healthcare) attached to a Shimadzu Nexera system using a flow of 0.45 mL/min collecting one fraction each minute. Detection of protein be done at UV absorbance of 280 nm. Fractions that contained pure enzyme were pooled, concentrated, aliquoted and stored at -20 °C in TBS containing 10% glycerol.

**General procedure for StrH glycan remodeling of an intact mAb**

Infliximab (Celltrion) was diluted to a concentration of 2 mg/mL in 50 mM MOPS pH 6.5, and an appropriate amount of galactosidase BgaA was added, the reaction mixture was incubated overnight at 37 °C with gently shaking. The ratio mAb:BgaA was 50:1 (w/w). Reaction progress was monitored using LC-ESI-MS. Once completed, to the A2F-glycan bearing mAb, TRIS buffer pH 7.5 to a concentration of 100 mM was added and the pH was adjusted to pH 7.5, followed by addition of StrH and incubation overnight at 37 °C while gentle shaking. The ratio mAb:StrH was 50:1 (w/w). After asymmetrization, the mAb was purified according to the protein A purification protocol.

**General procedure for terminal GlcNAc installation on an intact mAb**

The mAb of interest was diluted to a concentration of 2 mg/mL in 50 mM TRIS pH 7.5, supplemented with 10 mM $MnCl_2$, 10 mM UDP-GlcNAc (MU07955, Carbosynth) and hGnT-I (*H. sapiens*) and calf intestine alkaline phosphatase (CIAP) (M1821, Promega) and was incubated overnight at 37 °C with gently shaking. The ratio mAb:hGnT-1 was 100:1 (w/w) and 2 units CIAP per 100 µg mAb were added. Reaction progress was monitored using LC-ESI-MS. If starting material remained, an extra portion of hGnT-1 was added and the reaction mixture was further incubated at aforementioned conditions. Once completed, the mAbs were purified according to the protein A purification protocol.

**General procedure for terminal β1,4-galactose installation on an intact mAb**

The mAb of interest was diluted to a concentration of 2 mg/mL in 50 mM TRIS pH 7.5, supplemented with 10 mM $MnCl_2$, 10 mM UDP-Galactose (MU06699, Carbosynth) and B4GalT1 (*H. sapiens*) and calf intestine alkaline phosphatase (CIAP) (M1821, Promega) and was incubated overnight at 37 °C with gently shaking. The ratio mAb:B4GalT1 was 100:1 (w/w) and 2 units CIAP per 100 µg mAb were added. Reaction progress was monitored using LC-ESI-MS. If starting material remained, an extra portion of hST6Gal1 was added and the reaction mixture was further incubated at the aforementioned conditions. Once completed, the mAbs were purified according to the protein A purification protocol.

**General procedure for terminal α1,3-galactose installation on an intact mAb**

The mAb of interest was diluted to a concentration of 2 mg/mL in 50 mM TRIS pH 7.5, supplemented with 10 mM $MnCl_2$, 10 mM UDP-galactose (MU06699, Carbosynth) and mmGGTA1 (*M. musculus*) and calf intestine alkaline phosphatase (CIAP) (M1821, Promega) and was incubated overnight at 37 °C with gently shaking. The ratio mAb:mmGGTA1 was 100:1 (w/w) and 2 units CIAP per 100 µg mAb were added. Reaction progress was monitored using LC-ESI-MS. If starting material remained, an extra portion of mmGGTA1 was added and the reaction mixture was further incubated at the aforementioned conditions. Once completed, the mAbs were purified according to the protein A purification protocol.

**General procedure for Neu5Ac installation on an intact mAb**

The mAb of interest was diluted to a concentration of 2 mg/mL in 50 mM TRIS pH 7.5, supplemented with 10 mM $MnCl_2$, 20 mM CMP-Neu5Ac (MC04391, Carbosynth) and ST6Gal1 (*H. sapiens*), Pd26ST WT (*P. damselae*) or Psp26ST A366G/A235M (*P. sp.* JT-ISH-224) and calf intestine alkaline phosphatase (CIAP) (M1821, Promega) and was incubated overnight at 37 °C with gently shaking. The ratio mAb:sialyltransferase was 50:1 (w/w) and 2 units CIAP per 100 µg mAb were added. Reaction progress was monitored using LC-ESI-MS. If starting material remained, an extra portion of hST6Gal1 was added and the reaction mixture was further incubated at the aforementioned conditions. Once completed, the mAbs were purified according to the protein A purification protocol.

**General procedure for α-Gal removal on an intact mAb**

Before the α-galactosidase (*A. niger*) (E-AGLANP, Megazyme) was used for the modification of the mAbs glycan, it was purified by size-exclusion chromatography. To do so, 5 mg α-galactosidase powder was dissolved in 1 mL ultrapure water and purified as described in the general procedure for SEC above. To modify the mAb with the purified α-galactosidase, the mAb of interest was diluted to a concentration of 2 mg/mL in 50 mM NaOAc pH 4.5. The ratio mAb:α-galactosidase was 50:1 (w/w). After adding the α-galactosidase, the reaction mixture was incubated overnight at 37 °C with gently shaking. Reaction progress was monitored using LC-ESI-MS. If starting material remained, an extra portion of α-galactosidase was added, and the reaction mixture was further incubated at the aforementioned conditions. Once completed, the mAbs were purified according to the protein A purification protocol.

**Deglycosylation of mAbs with Endo-S2 WT or PNGase-F**

The mAb of interest was diluted to a concentration of 2 mg/mL in 50 mM NaOAc pH 4.5 for treatment with Endo-S2 WT or was diluted to a concentration of 2 mg/mL in 50 mM TRIS pH 7.5 for treatment with PNGase-F. Per 100 µg of mAb, 1 µg of Endo-S2 WT was added and incubated for 2 h at 37 °C with gently shaking. In general, the ratio mAb:Endo-S2 WT of 100:1 (w/w) was sufficient to yield the mAb with a GlcNAc-Fuc glycan. Per 100 µg of mAb, 1 µg of PNGase-F was added and incubated for 2 h at 37 °C with gently shaking. Again, the ratio mAb:PNGase-F of 100:1 (w/w) was sufficient to yield the product, a mAb with the Fc-glycan completely removed. Reaction progress was monitored using LC-ESI-MS. The mAb was purified according to the protein A purification protocol.

**Protein A purification of mAbs**

For purification of the glycomodified mAbs, a gravity flow Protein A resin column was used. The Protein A Sepharose CL-4B resin powder (GE17-0780-01, Cytiva) was left to swell for 1 h in TBS before pouring into the empty gravity flow column. In general, 2 mL of swollen resin was used to purify 2-10 mg mAb. The resin bed was equilibrated by a wash of 10 CV of TBS (25 mM TRIS, 150 mM NaCl, pH 7.5). Then, the protein mixture consisting of the mAb, glycosylhydrolases and -transferases, and other small molecules was loaded after adjusting the buffers pH to 7-8. Subsequently, the resin bed was washed with 10 CV of TBS, followed by elution with 10 CV of 100 mM glycine buffer (pH 3) in fractions of 2 mL. Usually, the mAb was eluted in the fractions 2-5 (after 2-5 CV). The pH of the eluate (10 mL) was directly neutralized by adding 1 mL TRIS 1M pH 8.5 and the eluate was then concentrated and buffer exchanged to ultrapure water using a 30 kDa MWCO Amicon Ultra-4 Centrifugal Filter (UFC8030, Millipore).

**LC-ESI-MS analysis of glycoprotein fragments**

For analysis of the mAb Fc-glycan, samples were pretreated with the enzyme IdeS. This enzyme cleaves below the hinge region (PAPELLG|GPSV), resulting in a single Fc fragment with the glycan at N297 of about ~25 kDa. Firstly, samples were adjusted to pH 7.5, then 10 units IdeS (A0-FR1, Genovis) were added to 10 µg of mAb and incubated at 37 °C for 30 min. A Vivaspin 500 10 kDa MWCO spin filter (VS0102, Sartorius) was used to buffer exchange the reaction mixture to ultrapure water, prior mass spectrometry analysis. Samples were injected with an Agilent 1290 Infinity LC equipped with an Acquity 300 1.7 um, 168 2.1 x 50 mm column (186003685, Waters) over a linear gradient of 20-40% B vs A in 7 min at 70 °C (A: 99.9% ultrapure water, 0.1% formic acid, B: 70% isopropanol, 20% acetonitrile, 9.9% ultrapure water, 0.1% formic acid) with a flow rate of 0.3 mL/min. The detector was an Agilent 6560 Ion Mobility LC/Q-TOF, see above for instrument settings.

**IgE binding experiments**

Binding capacity of IgE antibodies to the generated infliximab/cetuximab glycoforms was evaluated by applying either a monoclonal IgE antibody directed against α-Gal (clone: m86, Ab00532-14.0, Absolute antibody) or sera from two red meat allergic individuals using a direct ELISA. Red meat allergy diagnosed by an experienced allergologist was confirmed by a convincing history of objective symptoms following consumption of lamb, beef, or pork, including urticaria, tongue swelling, and diarrhea, alongside elevated serum specific IgE (sIgE) levels against α-Gal (89.9 kU/L and >100 kU/L). Patient 1 had an 8-year clinical history of red meat allergy (sIgE: lamb 5.0 kU/L, beef 20.1 kU/L, pork 11.2 kU/L), whereas Patient 2 presented with a 2-year clinical history. Ethical approval for the use of sera was granted by the Biobank Committee of the University Medical Center Utrecht (number: 23-094). Briefly, each infliximab/cetuximab variant (0.5 µg/well) was coated overnight in triplicates onto a MaxiSorp 96 well plate (44-2404-21, Invitrogen) after diluting them in phosphate-buffered saline (PBS).

After blocking with 1% bovine serum albumin (BSA) in PBS, either the IgE monoclonal antibody directed against α-gal (1 µg/mL) or sera from red meat allergic individuals (1:10) diluted in blocking buffer (1% BSA in PBS) was incubated for 1 h at RT. Bound IgE antibodies were detected by applying 0.5 µg/mL of a secondary goat anti-human IgE polyclonal antibody coupled with horse-radish peroxidase (clone: GOXHU, A18793, Invitrogen) for 1 h at RT. Visualization was achieved by adding tetramethylbenzidine (TMB) substrate for 15 min in the dark, the reaction was stopped with 1 M hydrochloric acid. The optical density (OD) was measured at 450 nm. IgE antibody binding was evaluated by normalizing OD values against the mean of the negative controls (for infliximab: variants **1**-**6** and **10**-**12**, for cetuximab: variants **16** and **17)** corrected for their standard deviation.

Validation of interday precision was carried out by repeating the experiment on two consecutive days and no significant differences, except within 7 (p=0.0002), was revealed between day one and two. Specificity of IgE was verified by the absence of binding to infliximab variants **1**-**6** and **10**-**12** and cetuximab variants **16** and **17**, all lacking α-Gal epitopes. The data were statistically analyzed with the mixed-effect model using Graphpad Prism (GraphPad Software) and the Tukey test was used to correct for multiple testing.

**SPR binding studies**

Affinity measurements to human FcγRs were performed using an IBIS MX96 (IBIS Technologies) device and a Continuous Flow Microspotter (Wasatch Microfluidics). An array of 4 concentrations of a 3-fold dilution series of C-terminally biotinylated human FcγRs was captured simultaneously on a SensEye G-streptavidin sensor in 1xPBS containing 0.075% (v/v) Tween-80. All FcγRs were from Sino Biological unless stated otherwise. The highest concentrations differed depending on the receptor and was 100 nM for hFcγRIIIa-158V and hFcγRIIIb-NA1 (in-house), 30 nM for hFcγRIIIa-158, hFcγRIIIb-NA2 (in-house), and 10 nM for hFcγRIIa-131H, hFcγRIIa-131R, and hFcγRIIb. A 2-fold dilution series of IgG variants thereof was injected ranging from 0.49 nM to 1000 nM in 1xPBS at pH 7.4 supplemented with 0.075% (v/v) Tween-80.

To measure the C-terminally His-tagged human FcγRI, 4 concentrations of a 3-fold dilutions series of C-terminally biotinylated anti-His mIgG1 (GenScript) was captured on a SensEye G-streptavidin sensor in 1xPBS containing 0.075% (v/v) Tween-80. Then 40 nM hFcγRI was subsequently captured and IgG variants thereof were titrated from 0.41 nM to 100 nM in a 3-fold dilution series in 1xPBS containing 0.075% (v/v) Tween-80. Regeneration of the sensor surface between the cycles was performed by injection of 10 mM Gly-HCl pH 2.1. Affinity calculations were carried out by performing an equilibrium analysis interpolating to an Rmax of 500 RU, and fitting a 1:1 Langmuir binding model, assuming equilibrium to be reached after 360s of IgG containing analyte injections, as described previously.[9] Analysis and calculations were performed using Scrubber Software Version 2 (BioLogic Software), Excel (Microsoft), and Graphpad Prism (GraphPad Software). Sensorgrams are provided in Figure S39.

### Figures and Tables

#### Figure S1. LC-MS and NMR analysis of glycan 27

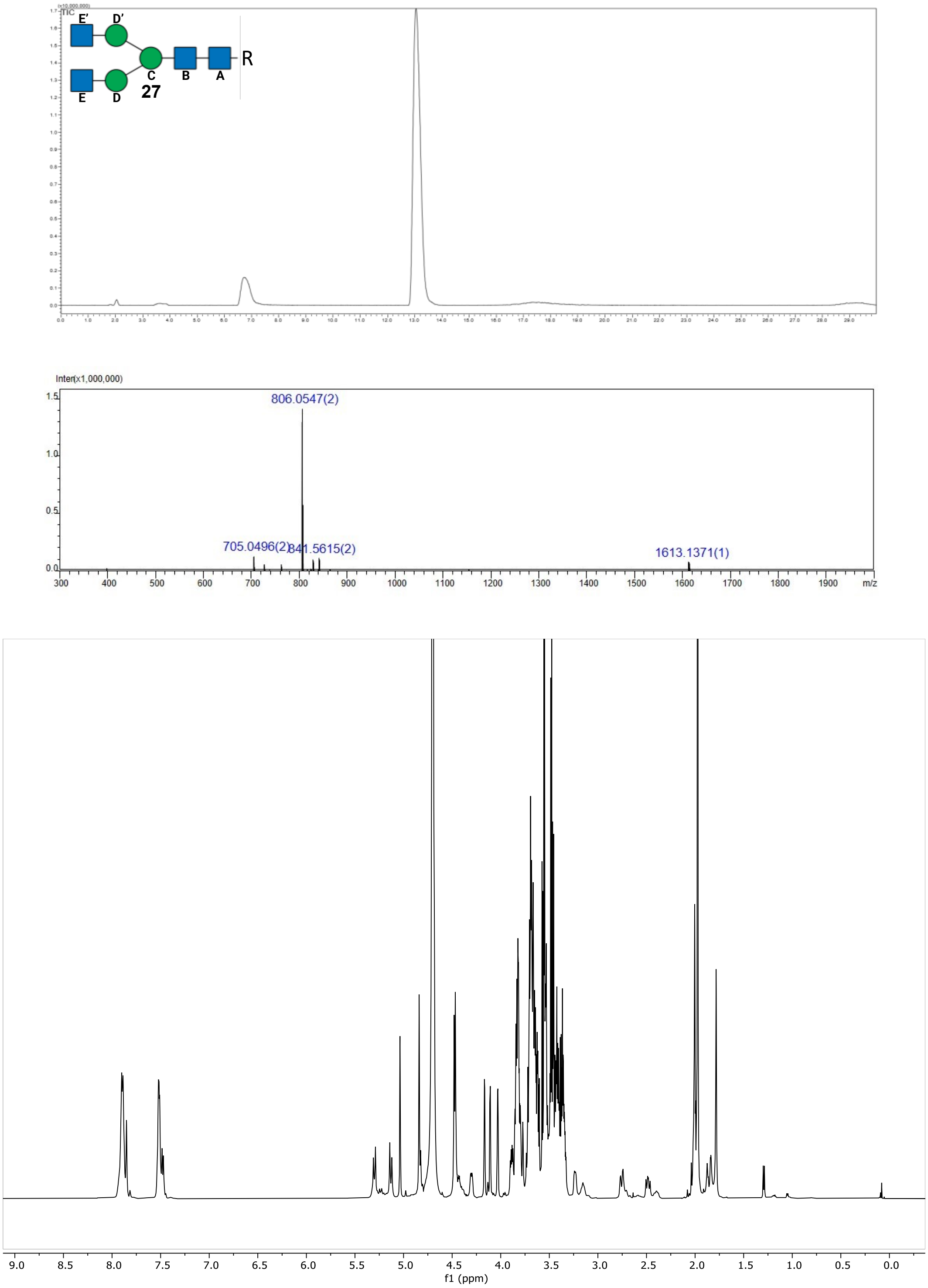


**Figure S1.** LC-MS and NMR analysis of glycan **27**. LC-MS glycan analysis of glycan **27** using a Waters XBridge BEH, Amide Column (3.5 μm, 2.1 x 150 mm). Over a linear gradient of 80-50% B vs A in 18 minutes at 25°C (A: 10 mM NH4HCOO in ultrapure water, pH3.5, B: ACN). Upper panel: TIC. Middle panel: ESI-Scan (+). Lower panel: The 1D $^{1}H$ NMR of glycan **27**, recorded at 298K in $D_2O$. Next pages: 2D $^{13}C$-$^{1}H$ COSY, TOCSY and HSQC spectra of glycan **27**.

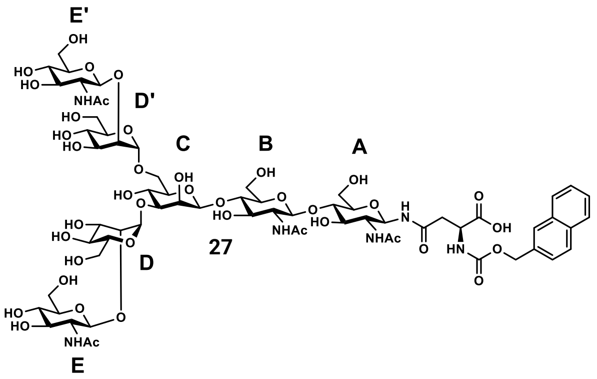


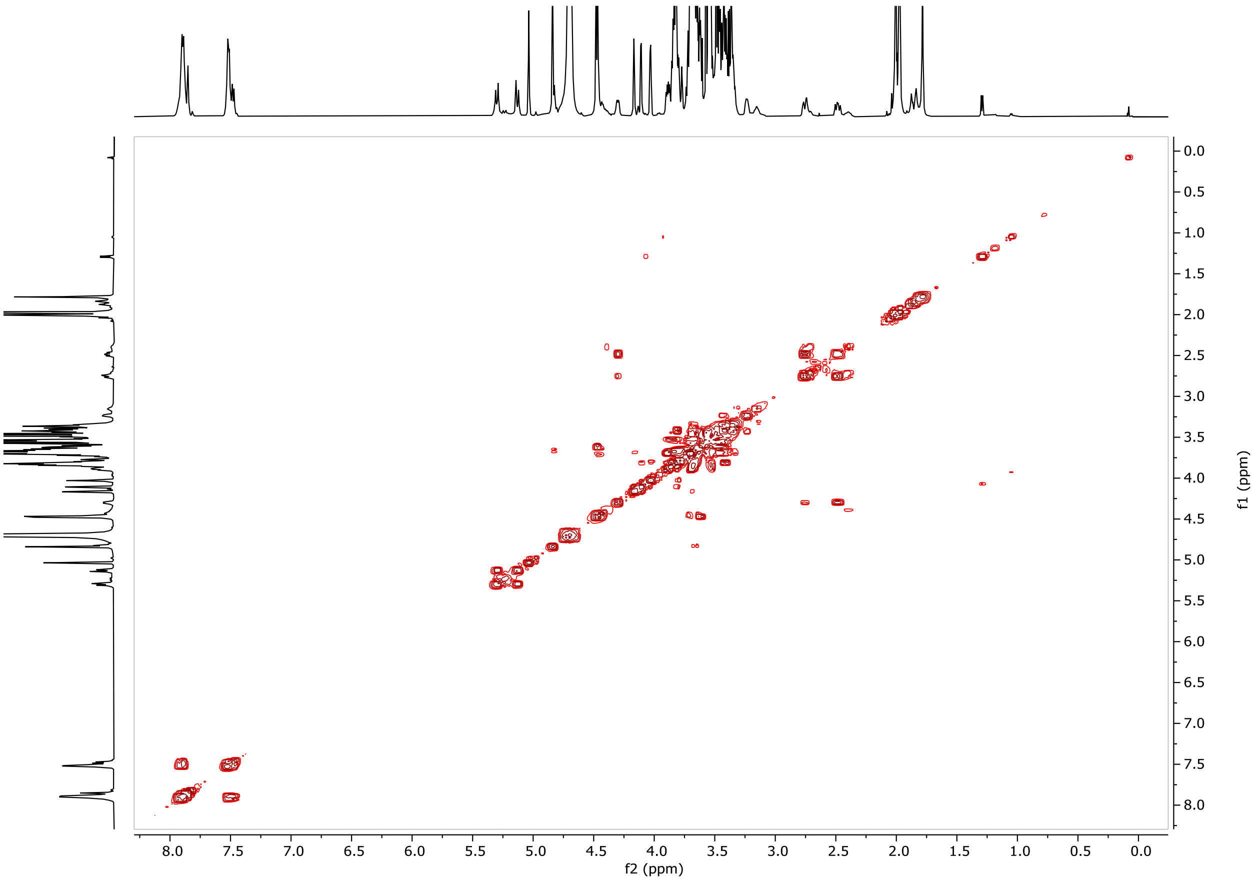


COSY

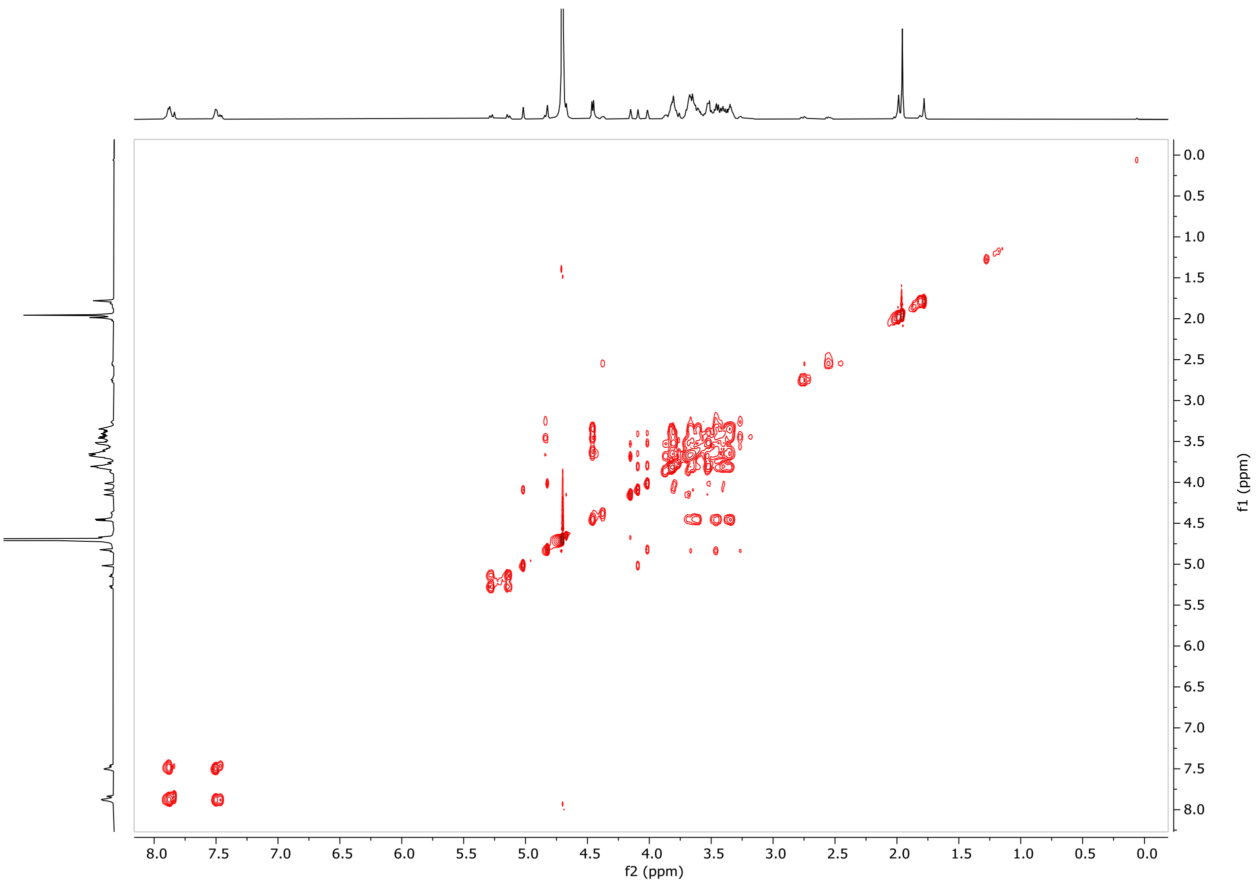


TOCSY

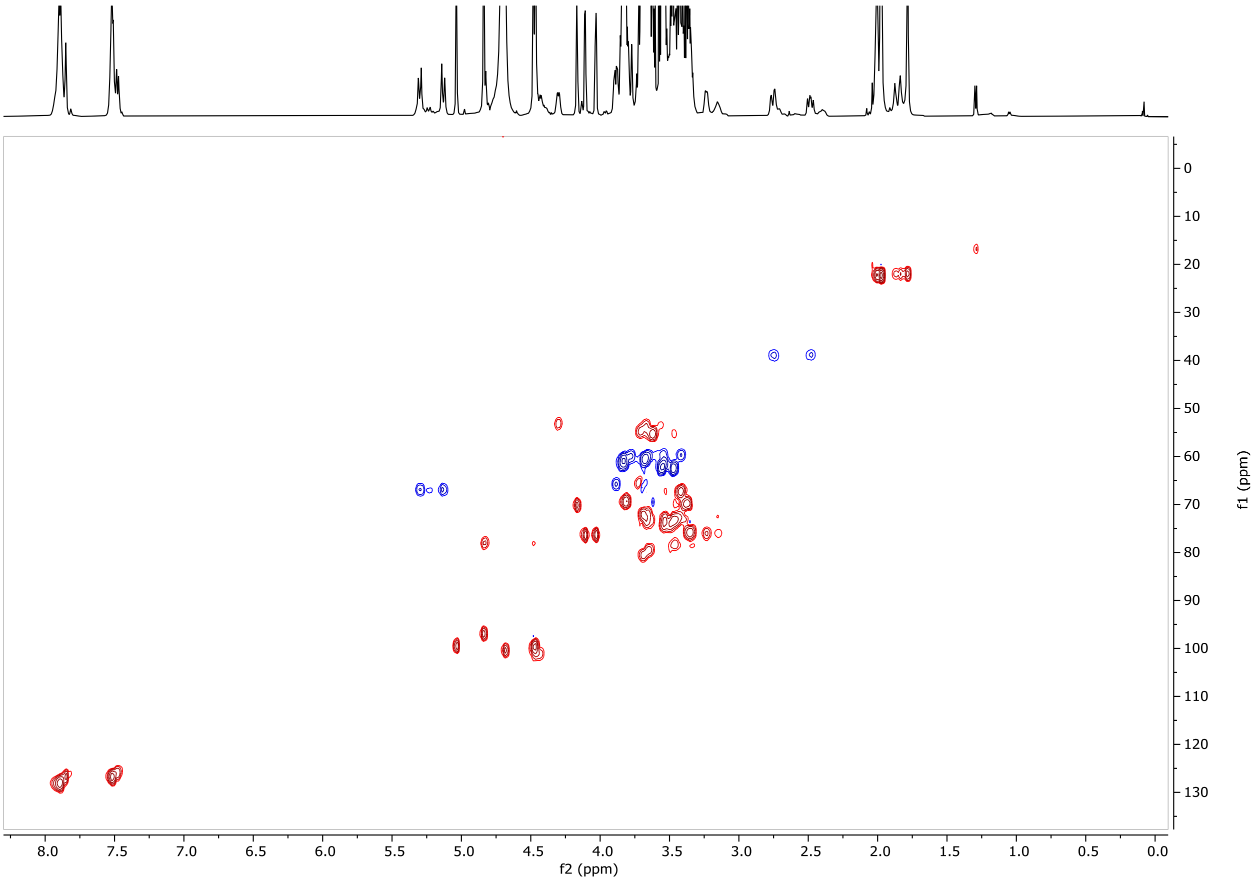

8.0
7.5
7.0
6.5
6.0
5.5
5.0
4.5
4.0
3.5
3.0
2.5
2.0
1.5
1.0
0.5
0.0
f2 (ppm)
0
10
20
30
40
50
60
70
80
90
100
110
120
130
f1 (ppm)


HSQC

#### Figure S2. LC-MS and NMR analysis of glycan 28

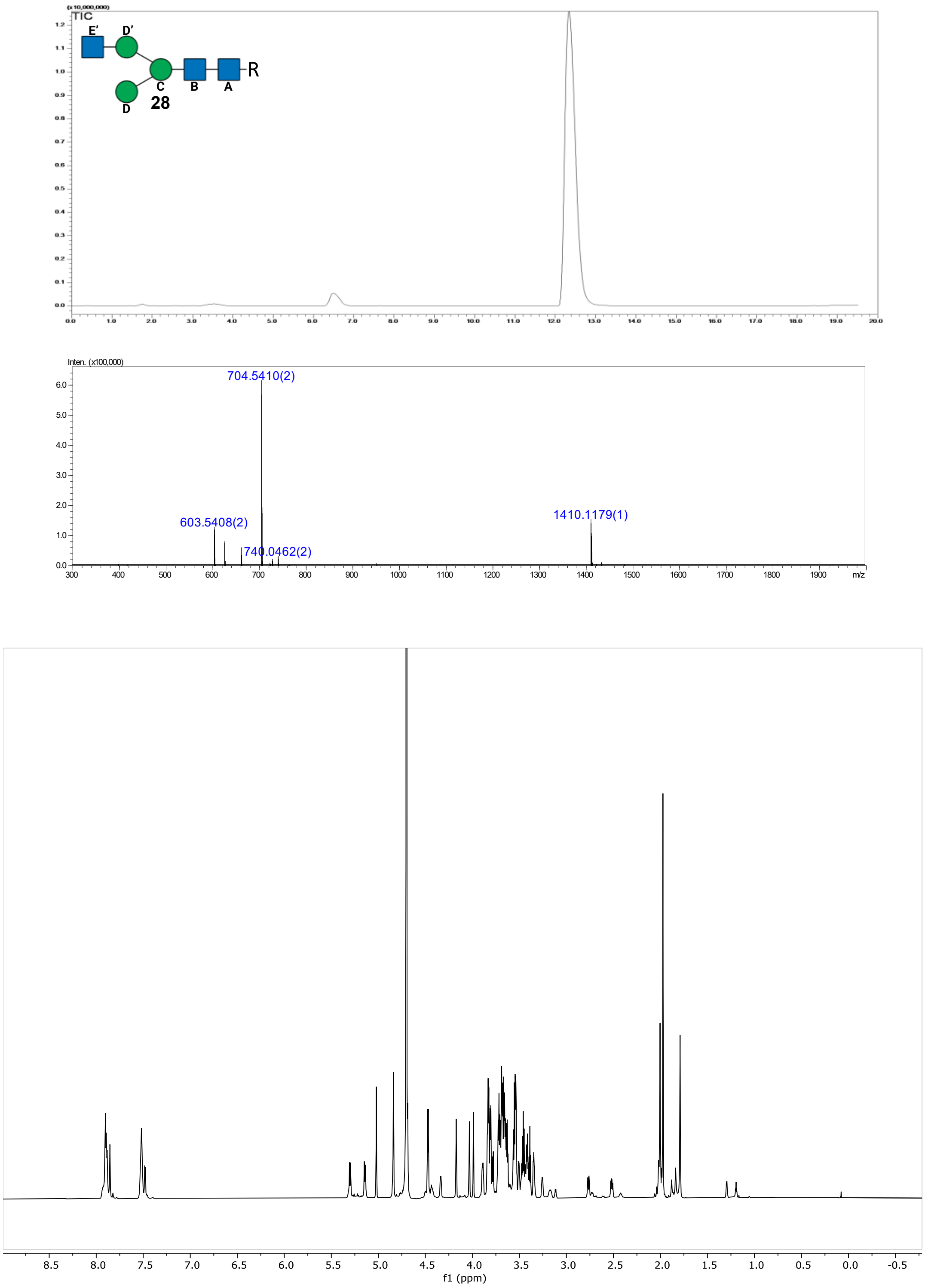


**Figure S2.** LC-MS and NMR analysis of glycan **28**. LC-MS glycan analysis of glycan **28** using a Waters XBridge BEH, Amide Column (3.5 µm, 2.1 x 150 mm). Over a linear gradient of 80-50% B vs A in 18 minutes at 25°C (A: 10 mM NH4HCOO in ultrapure water, pH3.5, B: ACN). Upper panel: TIC. Middle panel: ESI-Scan (+). Lower panel: The 1D $^{1}$H NMR of glycan **28**, recorded at 298K in $D_2O$. Next pages: 2D $^{13}$C-$^{1}$H COSY, TOCSY, HSQC and NOESY spectra of glycan **28**.

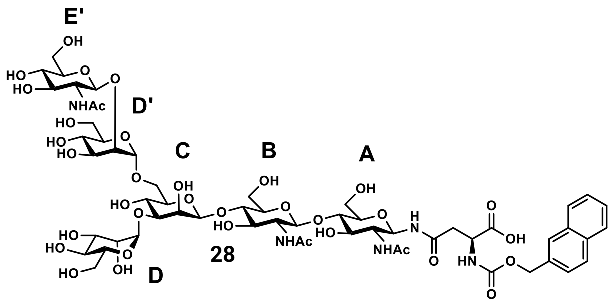


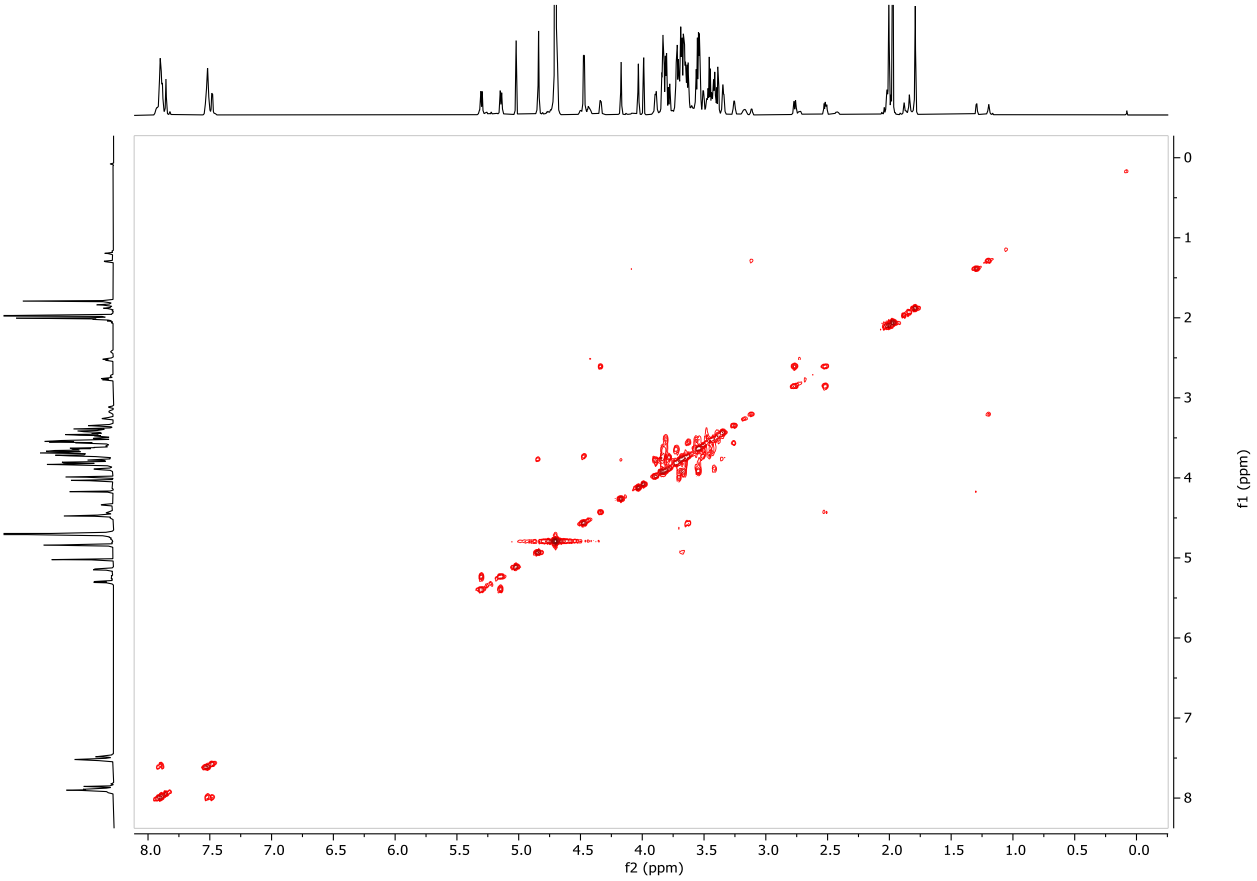


COSY

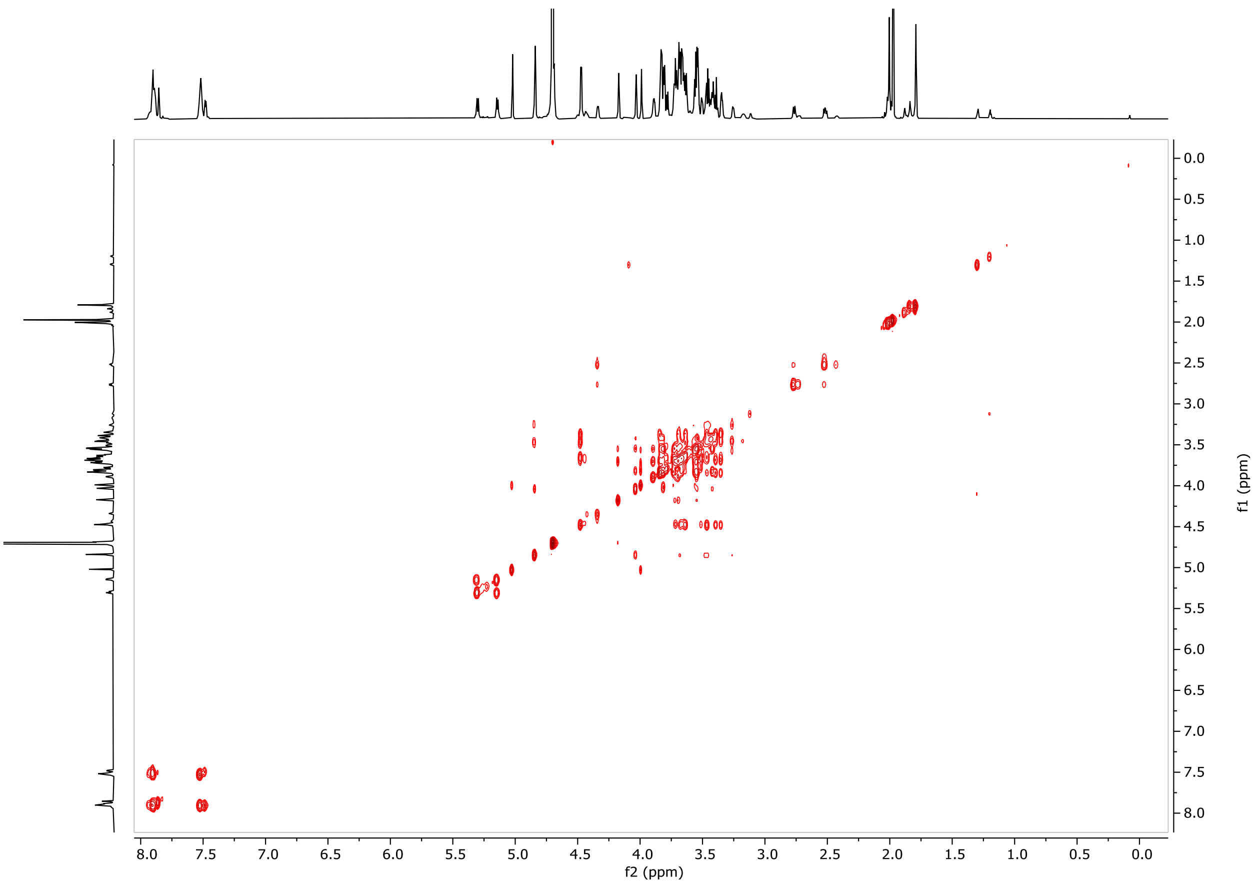


TOCSY

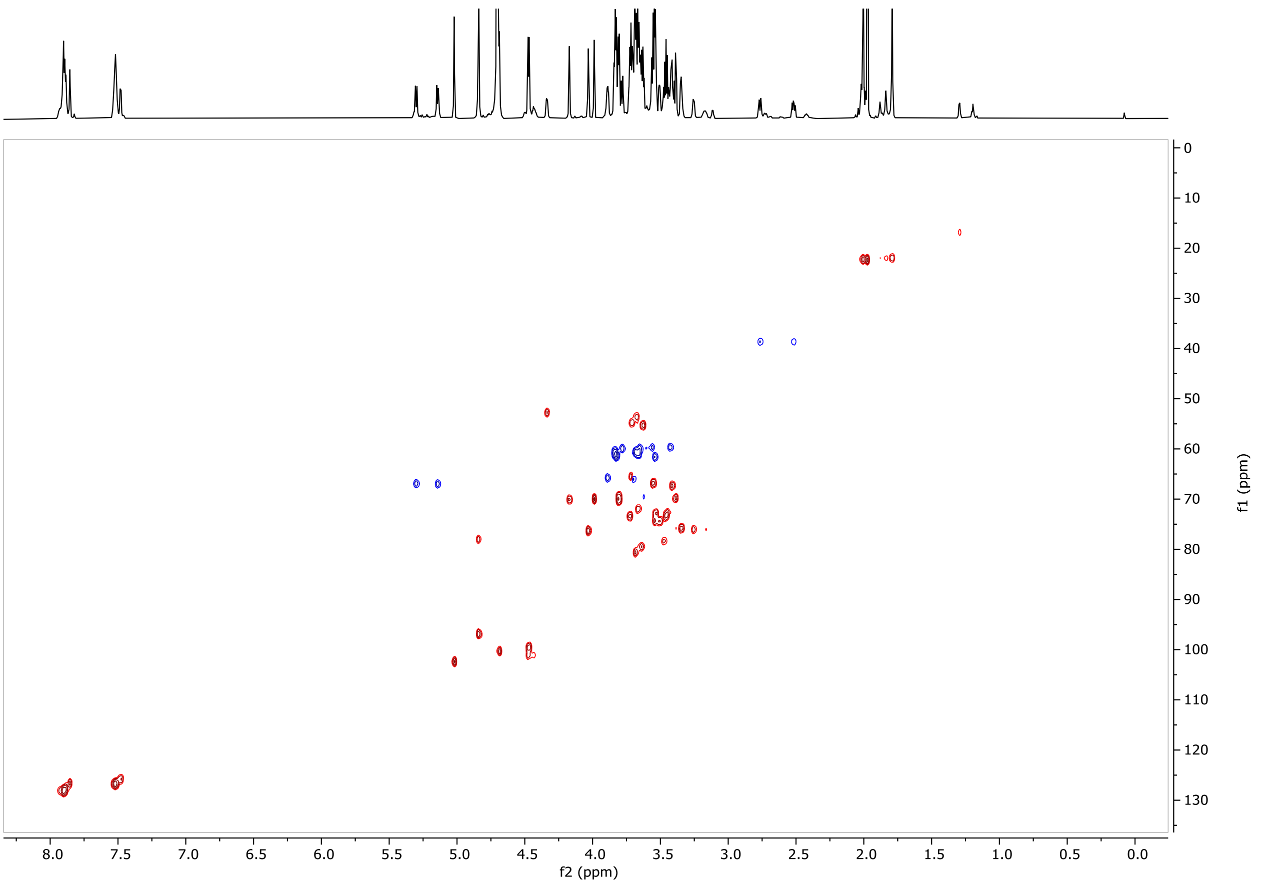


HSQC

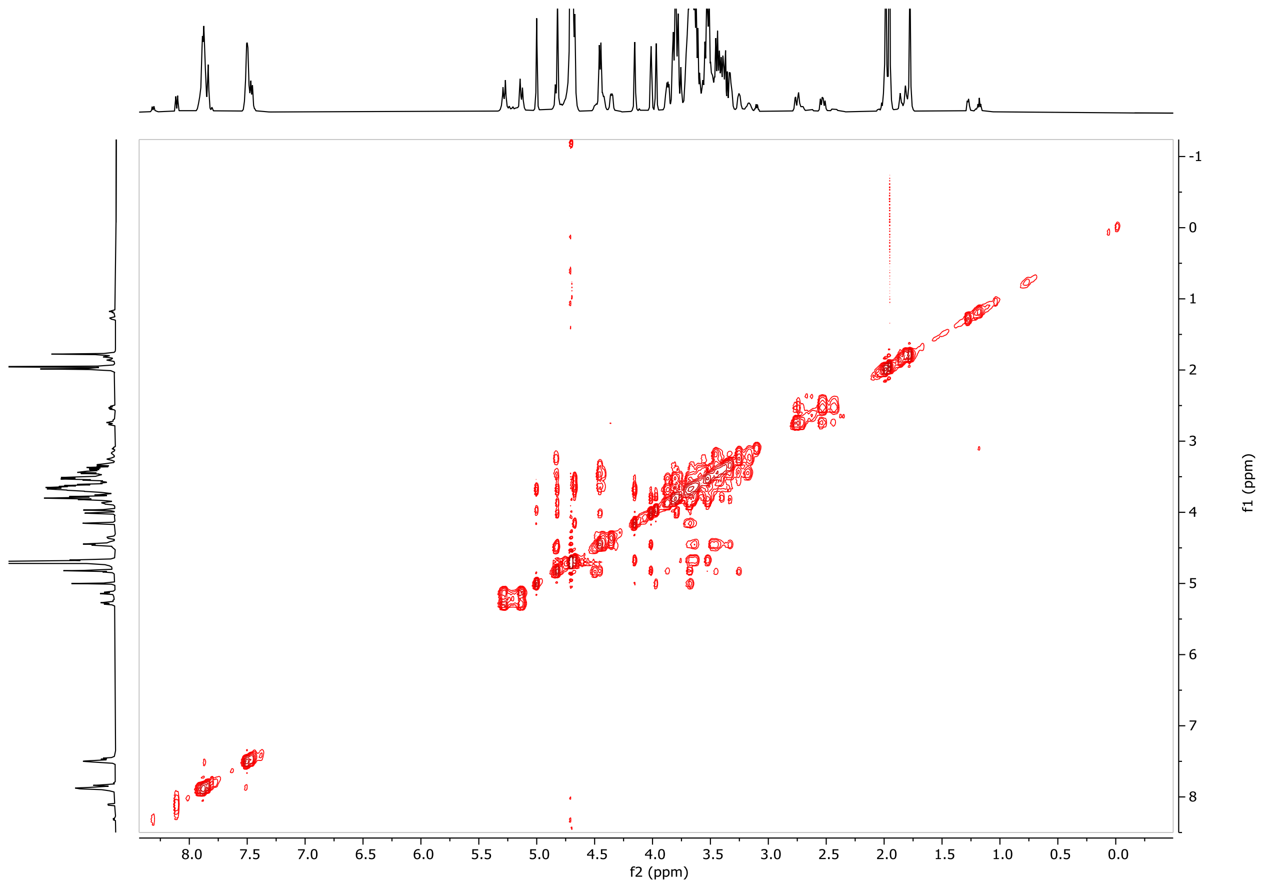


NOESY

### Figure S3. LC-MS and NMR analysis of glycan 29

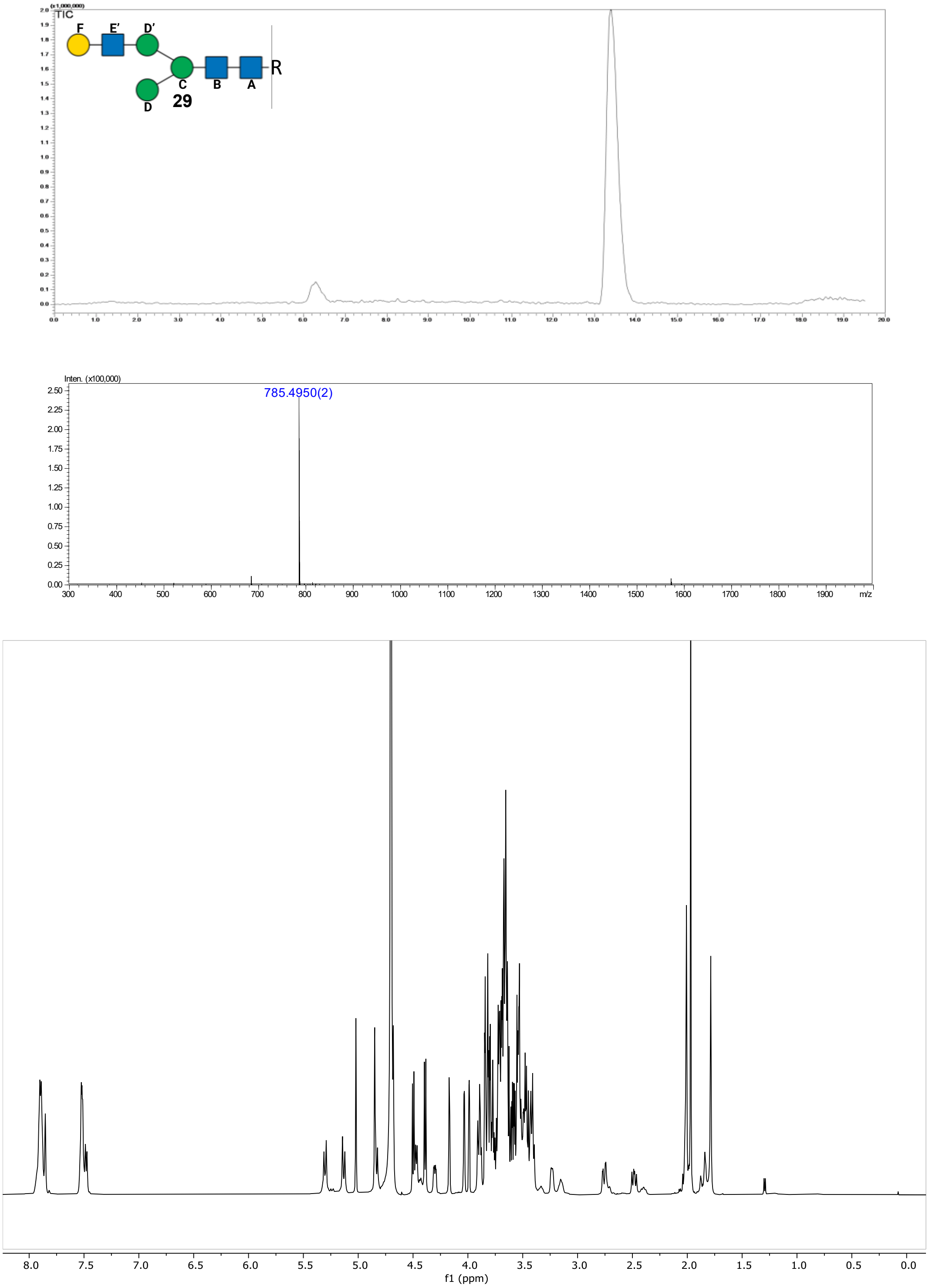


**Figure S3.** LC-MS and NMR analysis of glycan **29**. LC-MS glycan analysis of glycan **29** using a Waters XBridge BEH, Amide Column (3.5 μm, 2.1 x 150 mm). Over a linear gradient of 80-50% B vs A in 18 minutes at 25°C (A: 10 mM NH4HCOO in ultrapure water, pH3.5, B: ACN). Upper panel: TIC. Middle panel: ESI-Scan (+). Lower panel: The 1D $^{1}H$ NMR of glycan **29**, recorded at 298K in $D_2O$. Next pages: 2D $^{13}C$-$^{1}H$ COSY, TOCSY, HSQC and NOESY spectra of glycan **29**.

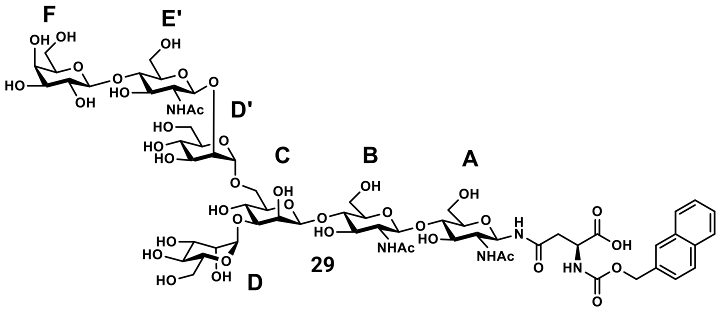


29

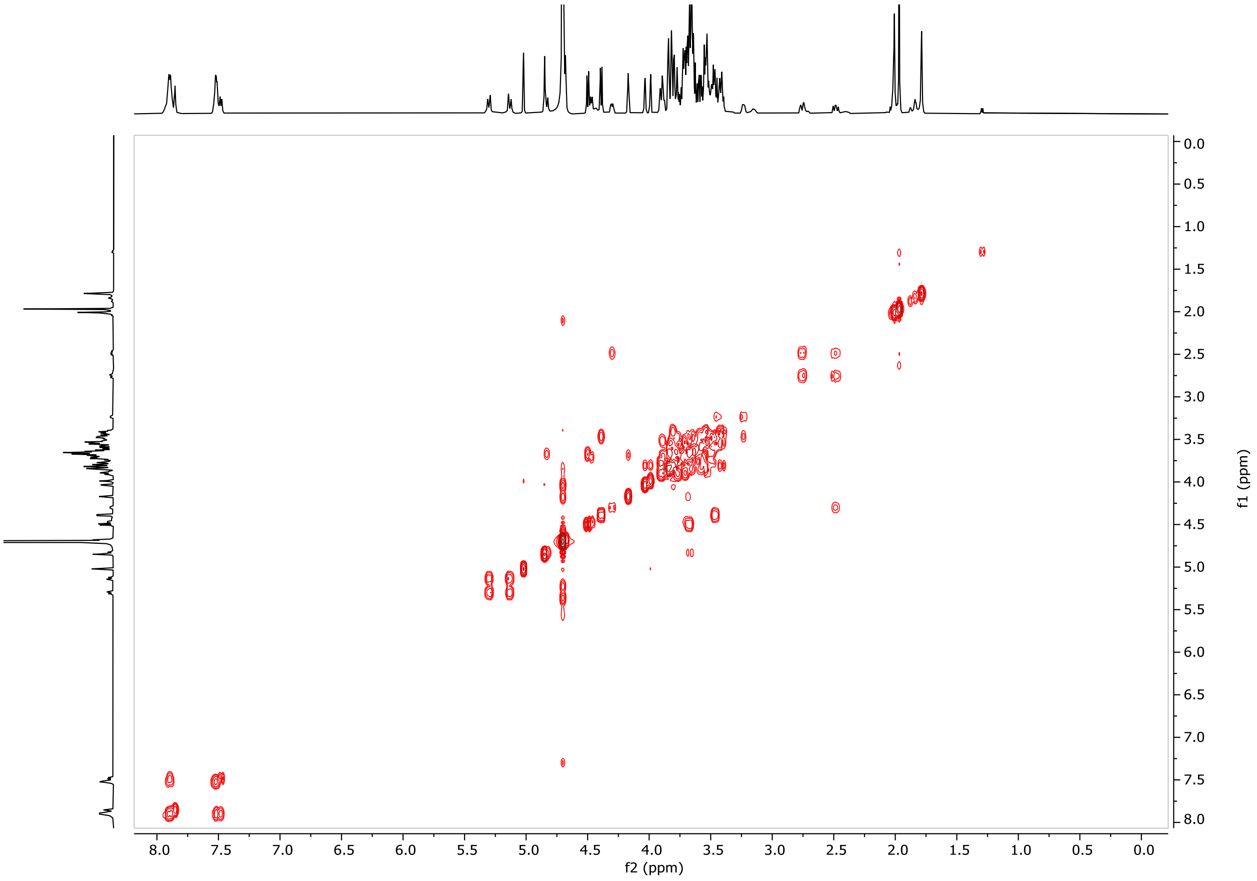


COSY

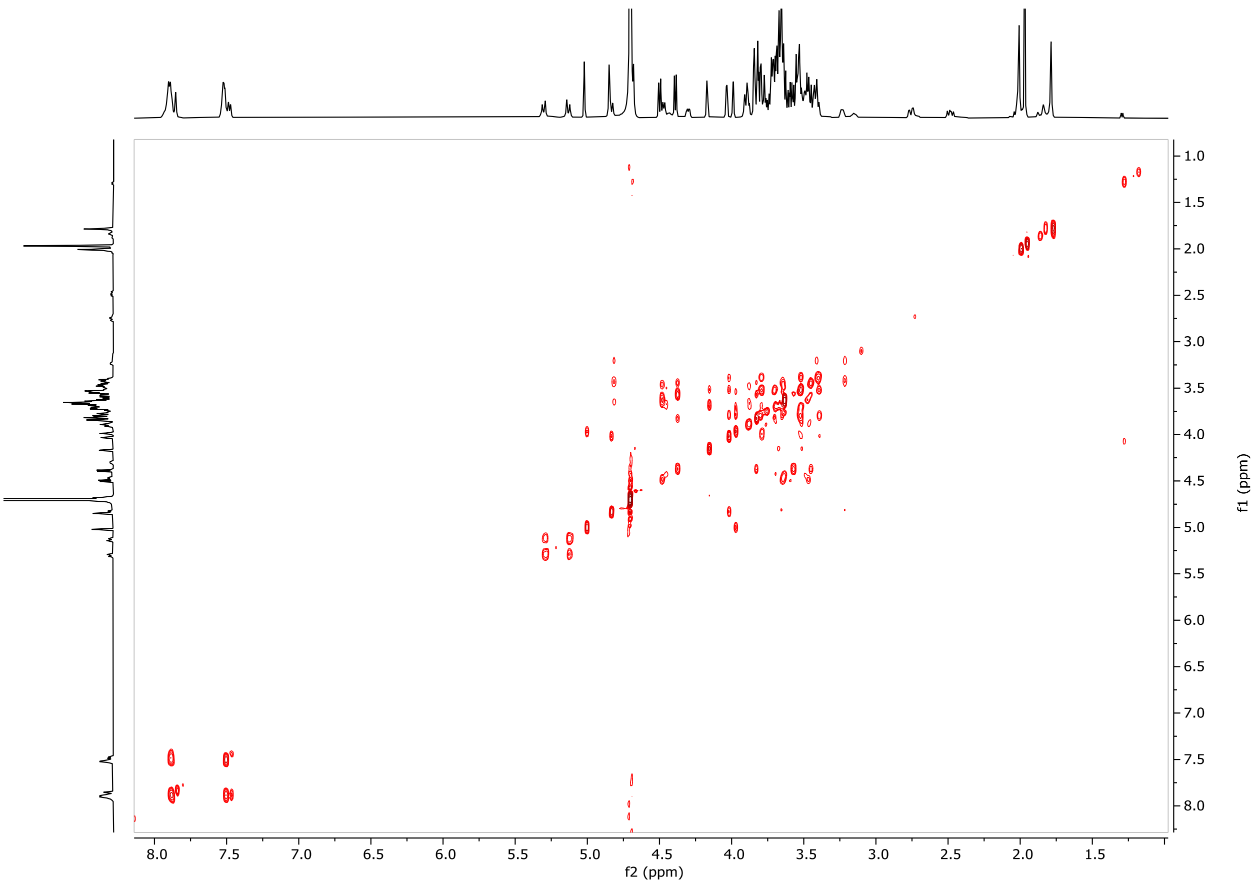

8.0
7.5
7.0
6.5
6.0
5.5
5.0
4.5
4.0
3.5
3.0
2.5
2.0
1.5
f2 (ppm)
1.0
1.5
2.0
2.5
3.0
3.5
4.0
4.5
5.0
5.5
6.0
6.5
7.0
7.5
8.0
f1 (ppm)


TOCSY

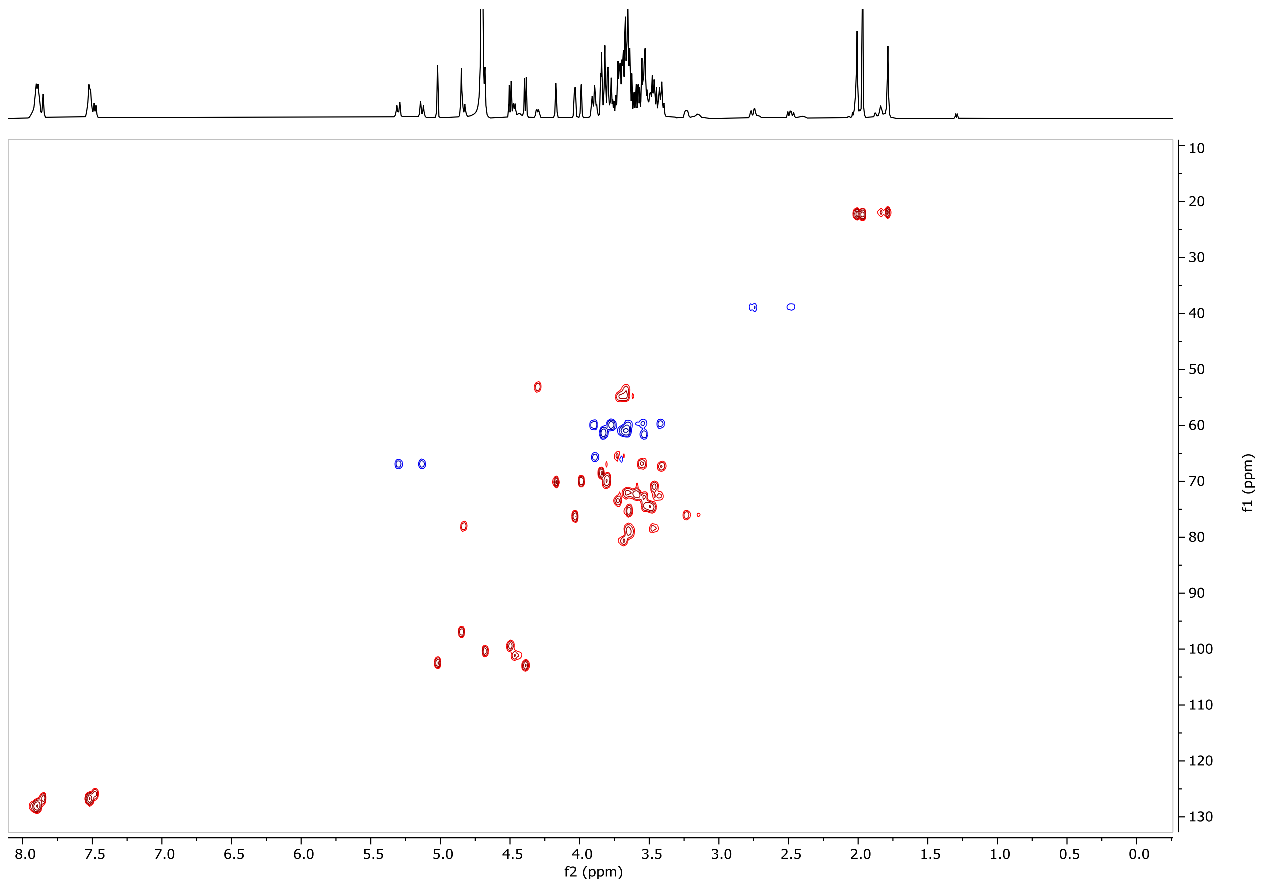


HSQC

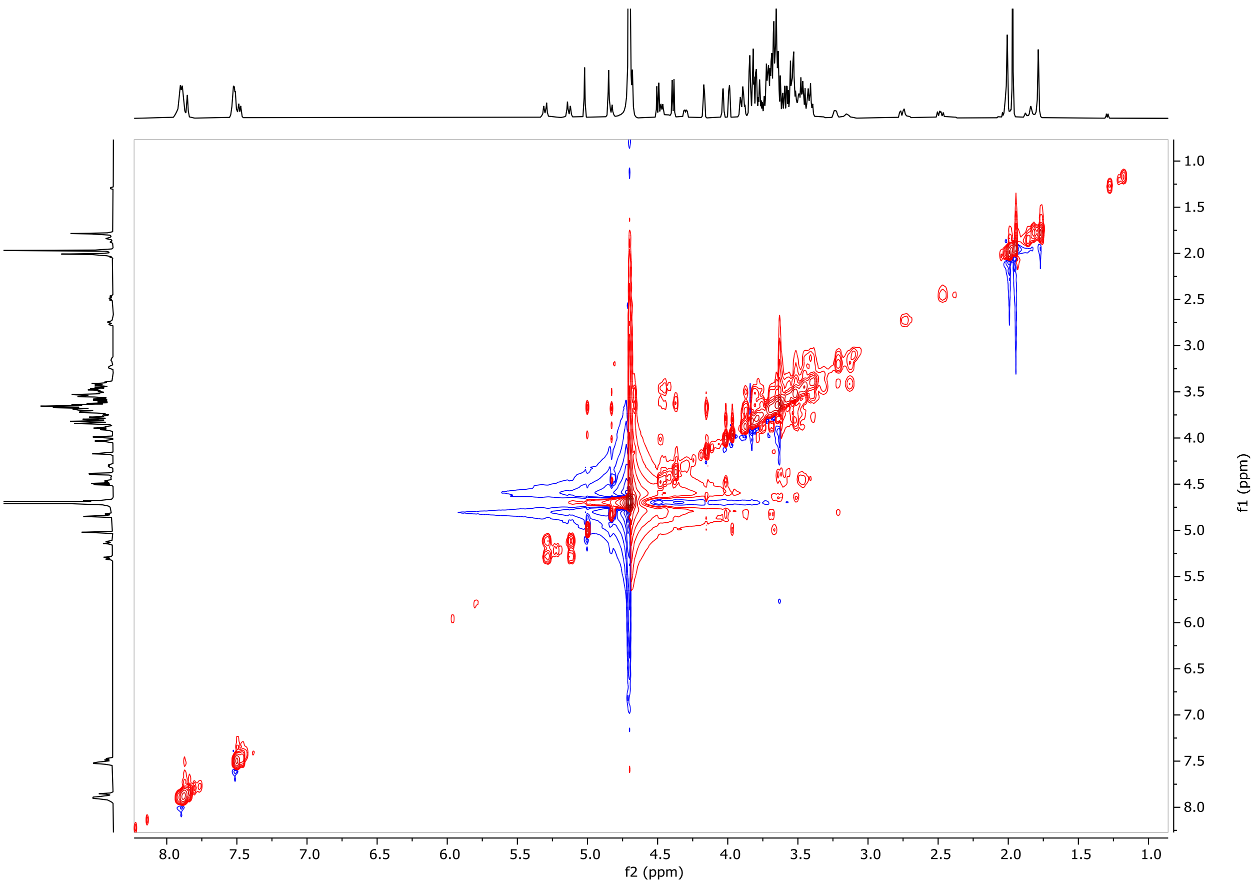


NOESY

#### Figure S4. LC-MS and NMR analysis of glycan 30

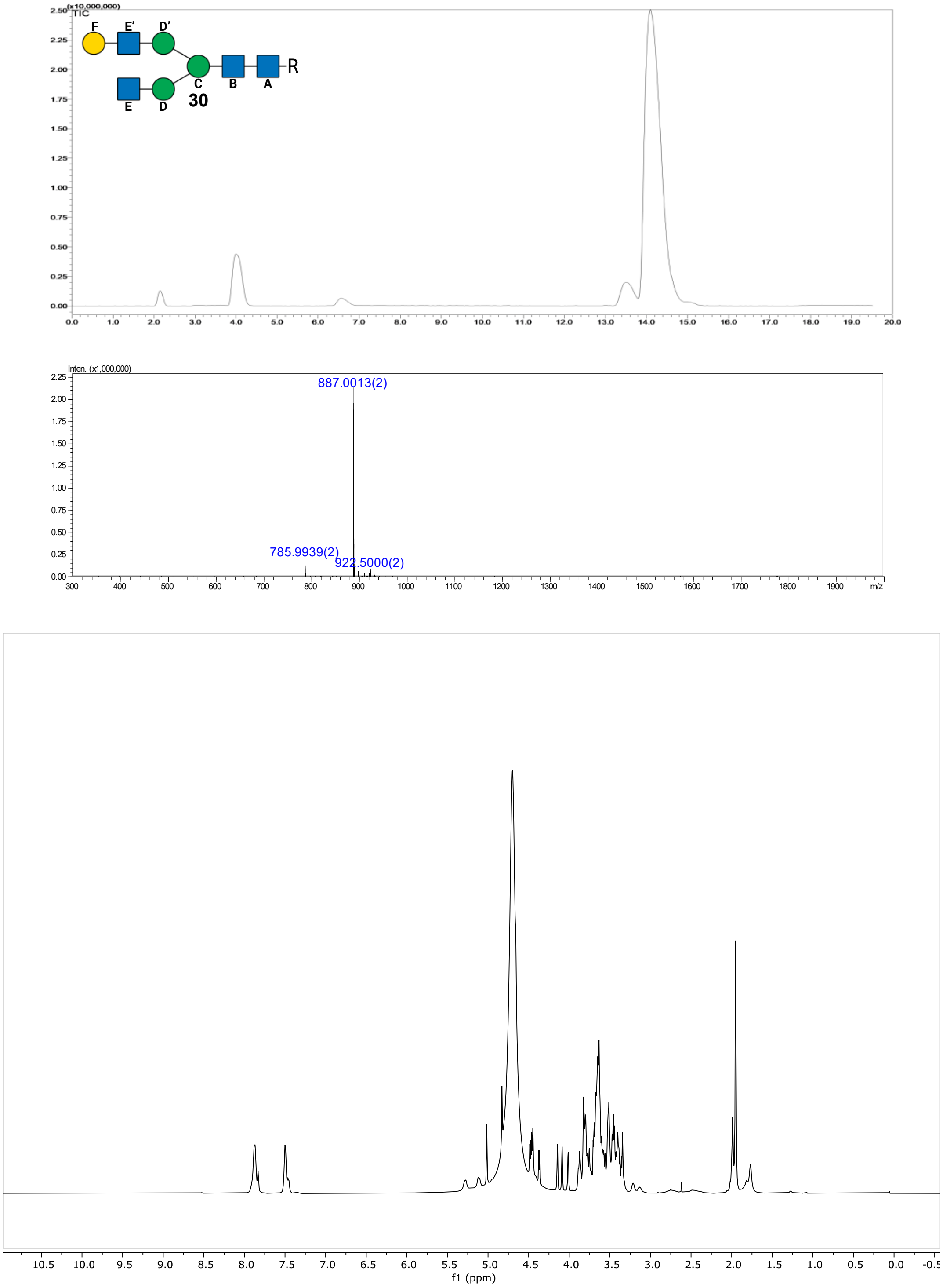


**Figure S4.** LC-MS and NMR analysis of glycan **30**. LC-MS glycan analysis of glycan **30** using a Waters XBridge BEH, Amide Column (3.5 μm, 2.1 x 150 mm). Over a linear gradient of 80-50% B vs A in 18 minutes at 25°C (A: 10 mM NH4HCOO in ultrapure water, pH3.5, B: ACN). Upper panel: TIC. Middle panel: ESI-Scan (+). Lower panel: The 1D $^{1}H$ NMR of glycan **30**, recorded at 298K in $D_2O$. Next pages: 2D $^{13}C$-$^{1}H$ COSY, TOCSY, HSQC and NOESY spectra of glycan **30**.

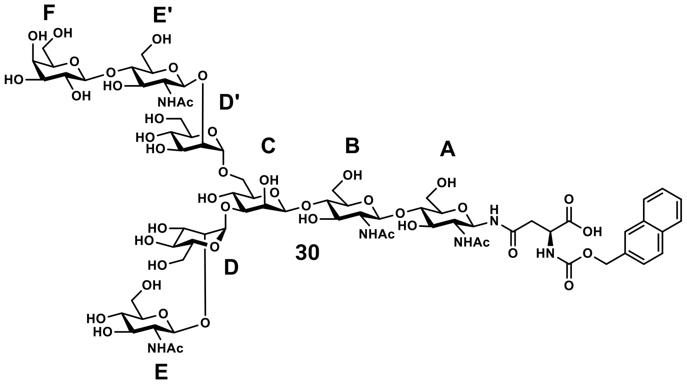


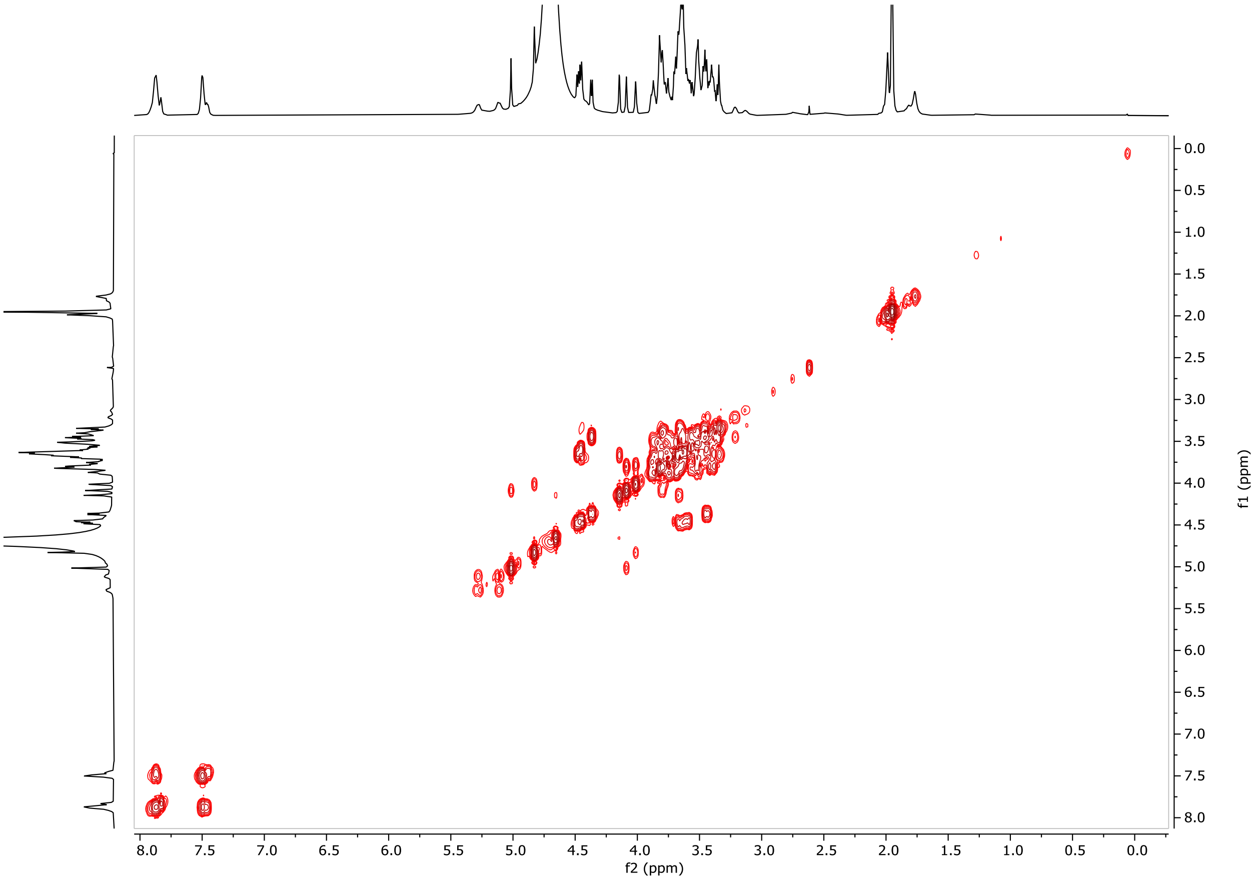


COSY

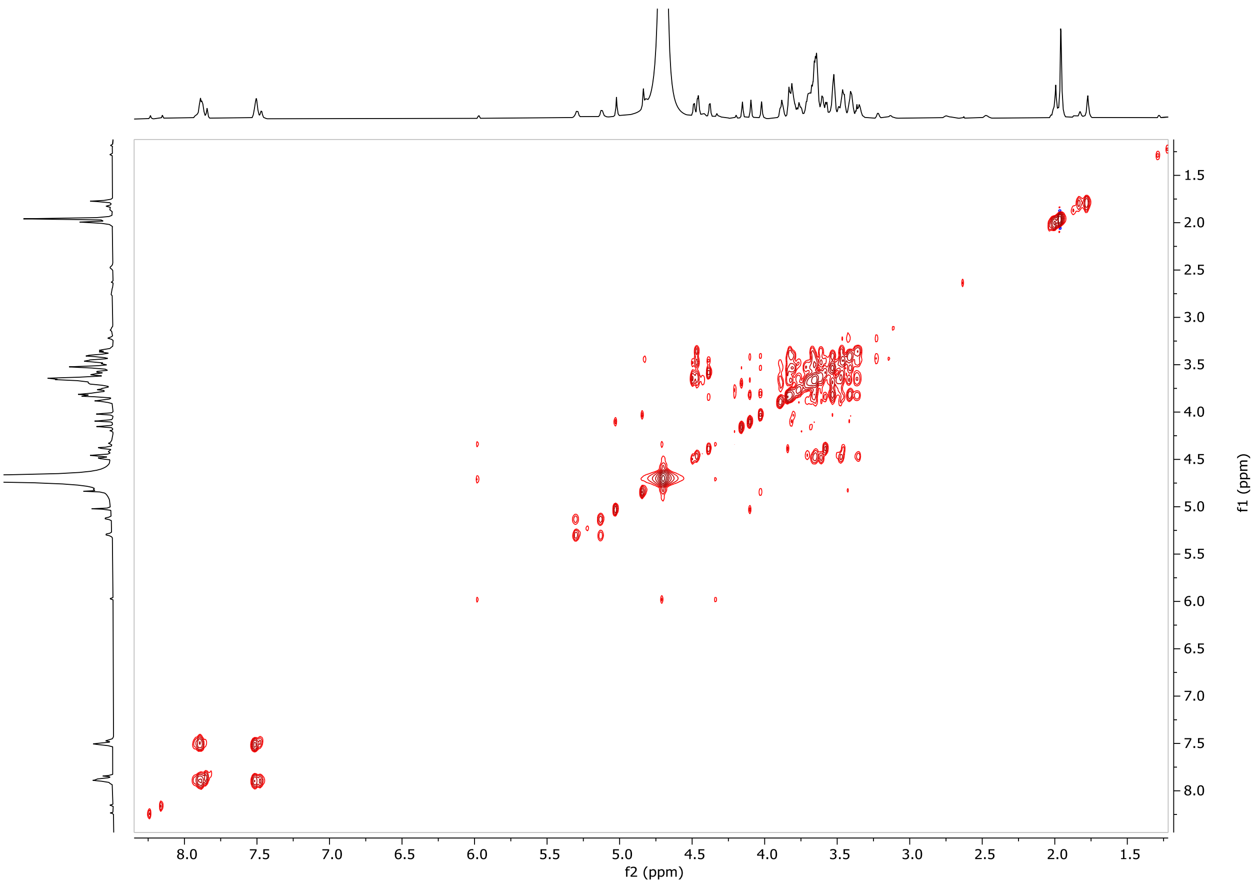


TOCSY

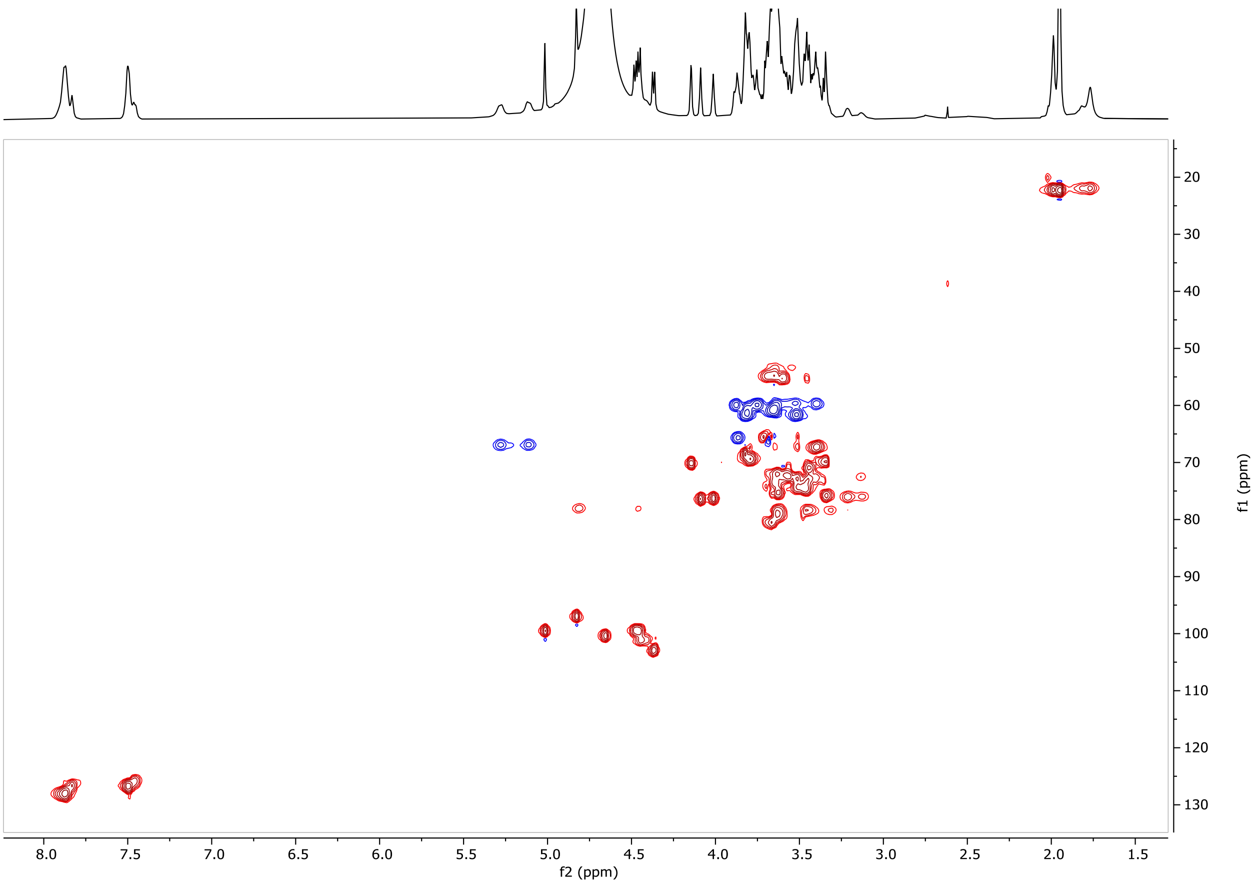


HSQC

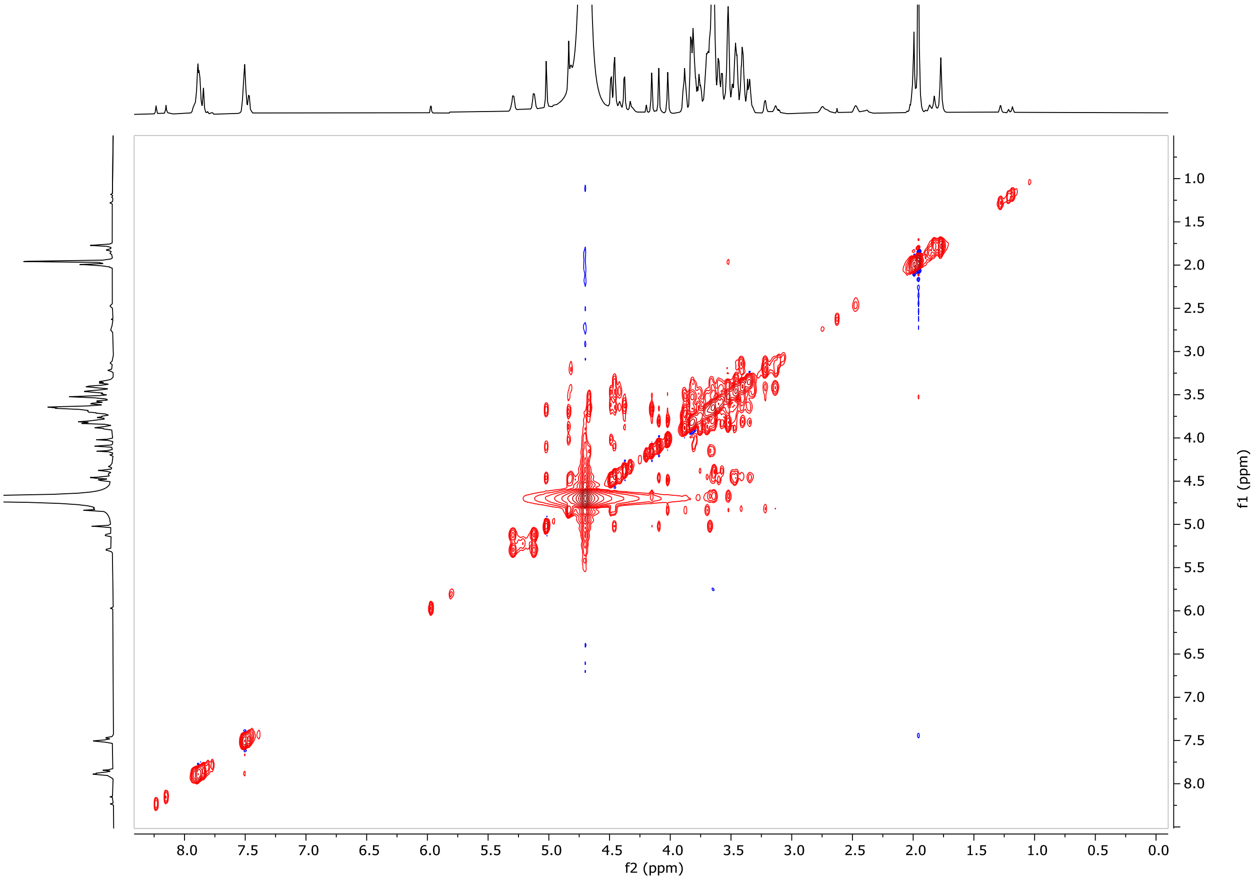


NOESY

#### Figure S5. NMR linkage analysis of glycans 28 and 30

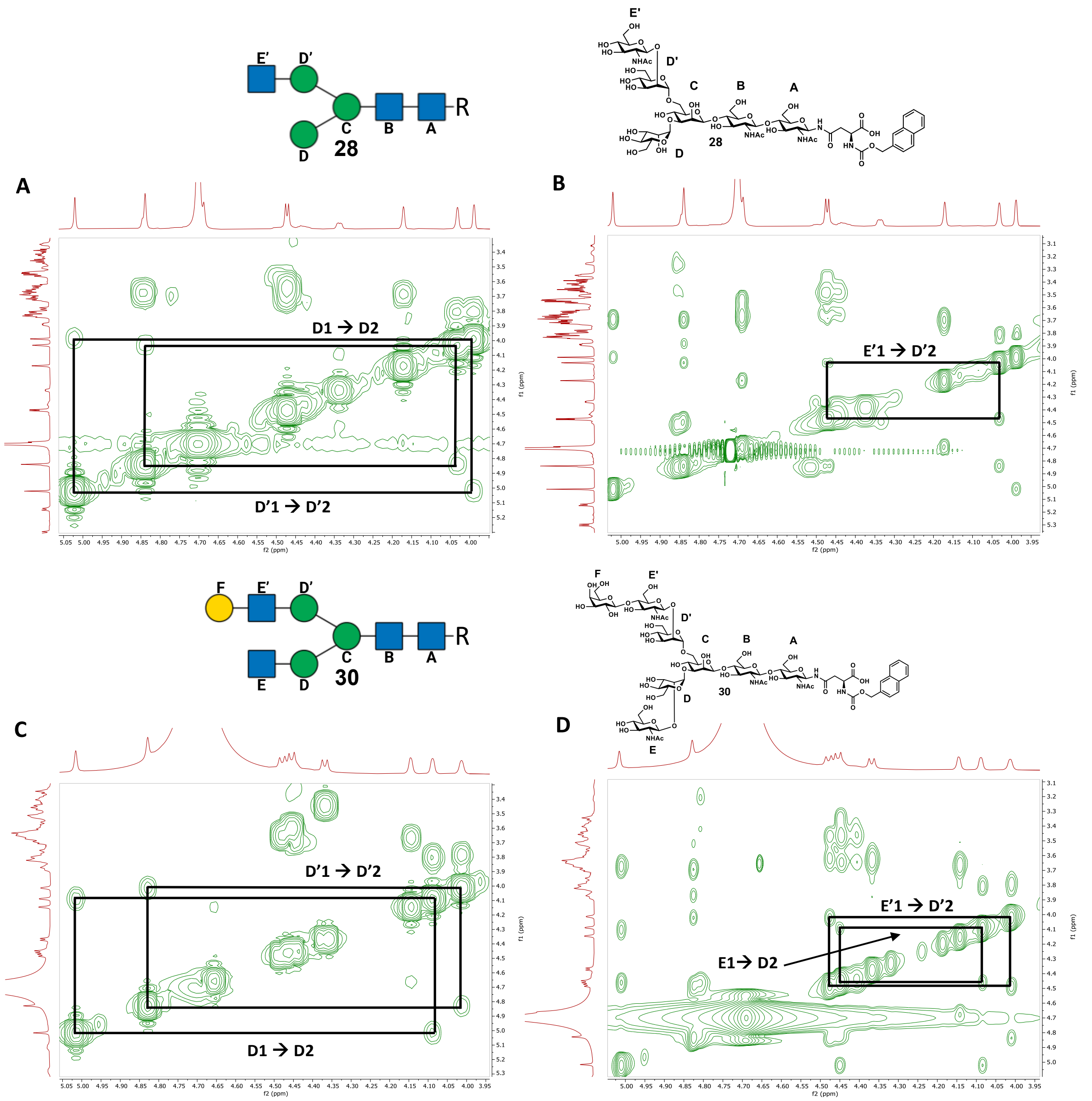


**Figure S5.** Linkage analysis of glycans **28** and **30** highlighting the (re-)installation of GlcNAc **E** to α1,3-mannose **D**. (A)&(C) COSY analysis to discern mannose H-2 shifts with and without the α1,3-mannose GlcNAc **E**. (B)&(D) NOESY spectrum to showcase the glycosidic NOEs between terminal GlcNAc H-1 and the H-2 of the α1,6-mannose (glycan **28**) as well as the α1,3-mannose (glycan **30**).

#### Figure S6. PGC-MS chromatogram of released glycan mAb 4

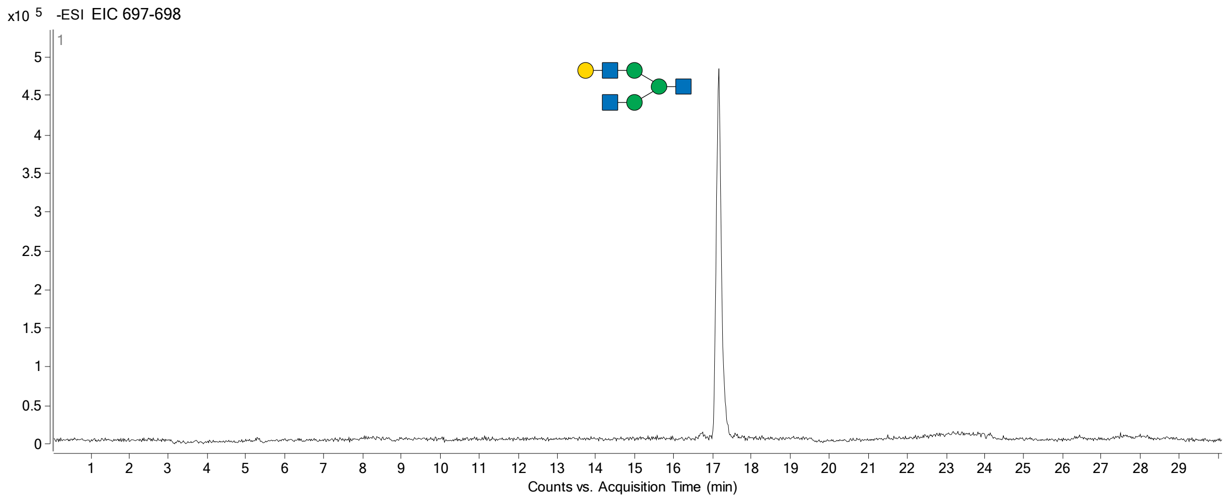


**Figure S6.** PGC-MS run of released and labelled glycan of mAb **4**. The extracted ion chromatogram shows a single peak in the mass range of 697-698 m/z. No structural isomers were detected.

#### Figure S7. Glycopeptide analysis of mAb 4

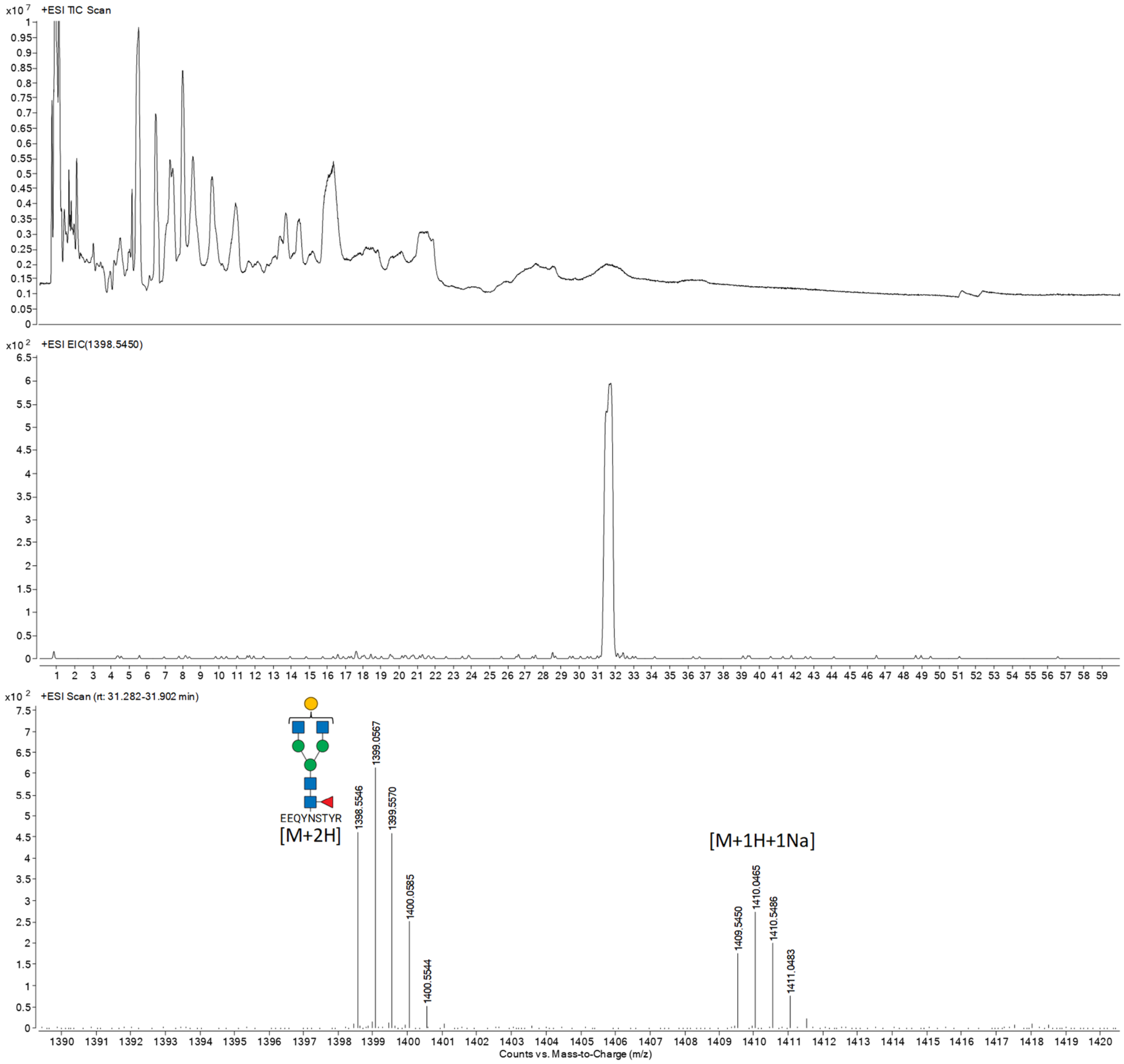


**Figure S7.** Trypsin digest of mAb **4** run on an AdvanceBio Glycan Mapping column, 2.1 × 150 mm, 2.7 µm, (683775913, Agilent) using an Agilent 1290 Infinity LC. Peptides were separated by running a linear gradient of 90-50% B vs A in 60 min at 25 °C (A: 0.1% FA in ultrapure water, B: ACN) with a flow rate of 0.3 mL/min. Top panel: total ion count (TIC). Middle panel: extracted ion chromatogram (EIC). Bottom panel: detected masses between 31.282-31.902 min. Using the parameters above, glycopeptides would elute > 25 min, by scanning beyond that timepoint a single glycoform could be identified. Findings were recorded in Table S1.

**Table S1. Detected glycopeptides of mAb 4 tryptic digest**

| Peptide | Glycan | Mono-isotopic mass | [M+2H] | Detected? |
|---|---|---|---|---|
| EEQYNSTYR | A0 | 2080.83 | 1041.42 | n.d. |
| 1188.5 g/mol | A1 | 2283.91 | 1142.96 | n.d. |
| | A1G1 | 2445.96 | 1223.98 | n.d. |
| | A2G1 | 2649.04 | 1325.52 | n.d. |
| | A2G2 | 2811.09 | 1406.55 | n.d. |
| | A0F | 2226.88 | 1114.44 | n.d. |
| | A1F | 2429.96 | 1215.98 | n.d. |
| | A1G1F | 2592.01 | 1297.01 | n.d. |
| | A2G1F | 2795.09 | 1398.55 | ✓ |
| | A2G2F | 2957.14 | 1479.57 | n.d. |
| TKPREEQYNSTYR | A0 | 2563.13 | 1282.57 | n.d. |
| 1670.8 g/mol | A1 | 2766.21 | 1384.11 | n.d. |
| | A1G1 | 2928.26 | 1465.13 | n.d. |
| | A2G1 | 3131.34 | 1566.67 | n.d. |
| | A2G2 | 3293.39 | 1647.70 | n.d. |
| | A0F | 2709.18 | 1355.59 | n.d. |
| | A1F | 2912.26 | 1457.13 | n.d. |
| | A1G1F | 3074.31 | 1538.16 | n.d. |
| | A2G1F | 3277.39 | 1639.70 | n.d. |
| | A2G2F | 3439.44 | 1720.72 | n.d. |

n.d. - not detected

**Figure S8. Sialylation optimization using mAb 4**

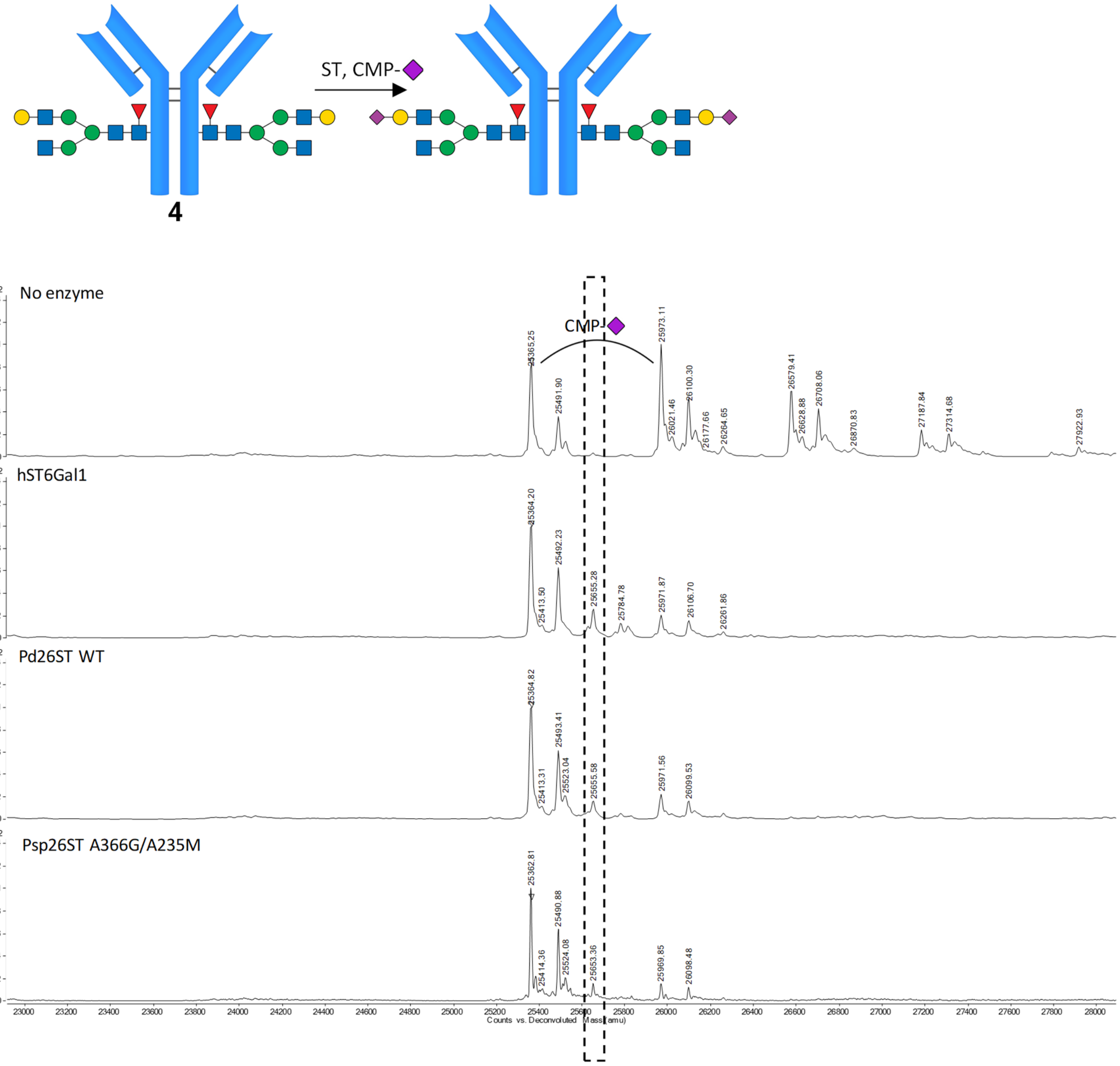


Product peak: 25656 Da

(Percentage from maximum height start material; 25364 Da)

| | |
|---|---|
| hST6Gal1 | 25.7% |
| Pd26ST WT | 16.4% |
| Psp26ST A366G/A235M | 15.5% |

**Figure S9. Sialylation optimization using mAb 19**

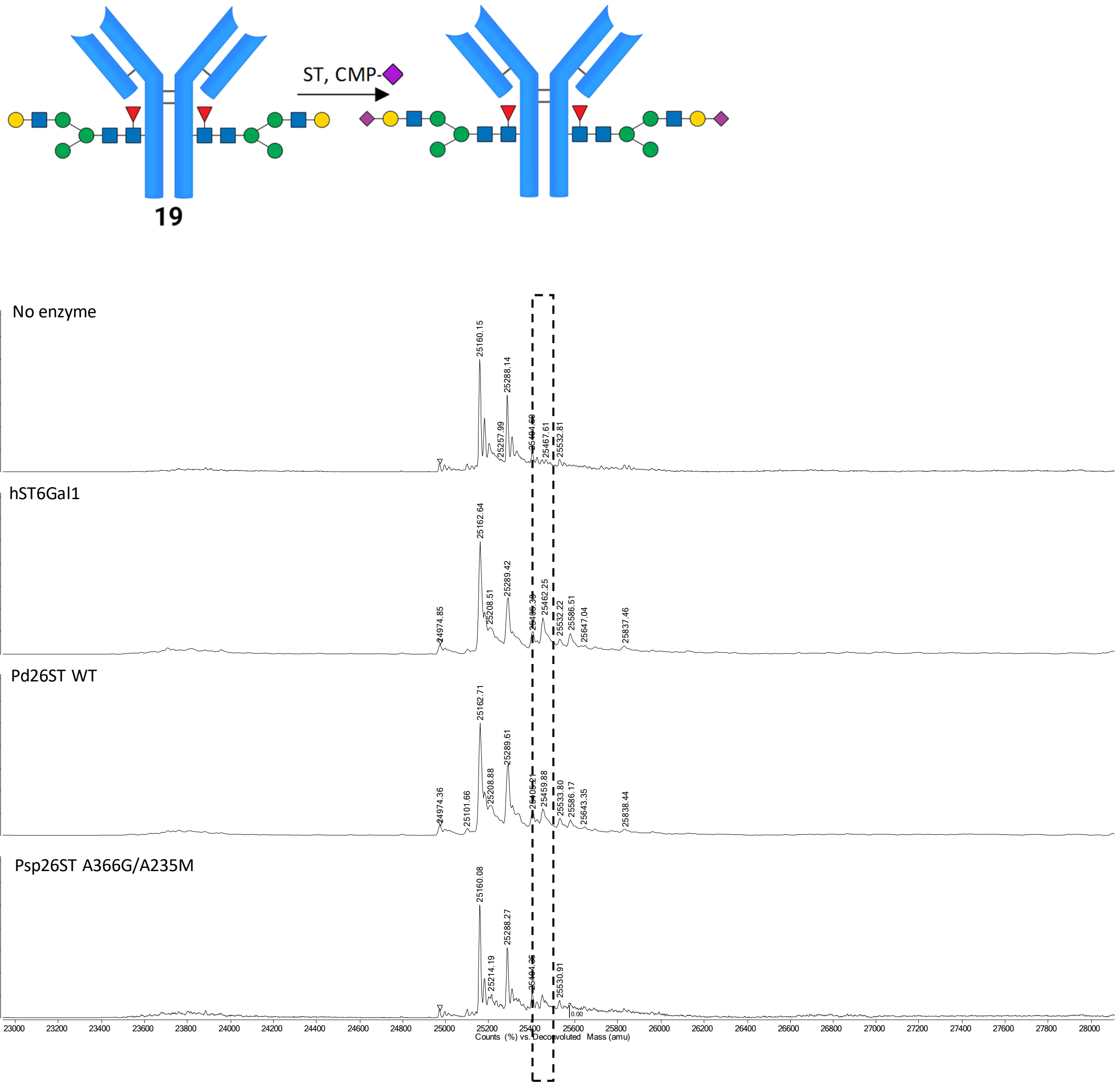


Product peak: 25451 Da

(Percentage from maximum height start material; 25160 Da)

| | |
|---|---|
| hST6Gal1 | 35.4% |
| Pd26ST WT | 24.4% |
| Psp26ST A366G/A235M | 18.9% |

**Figure S10. Complete enzymatical, asymmetrical glycoremodeling route**

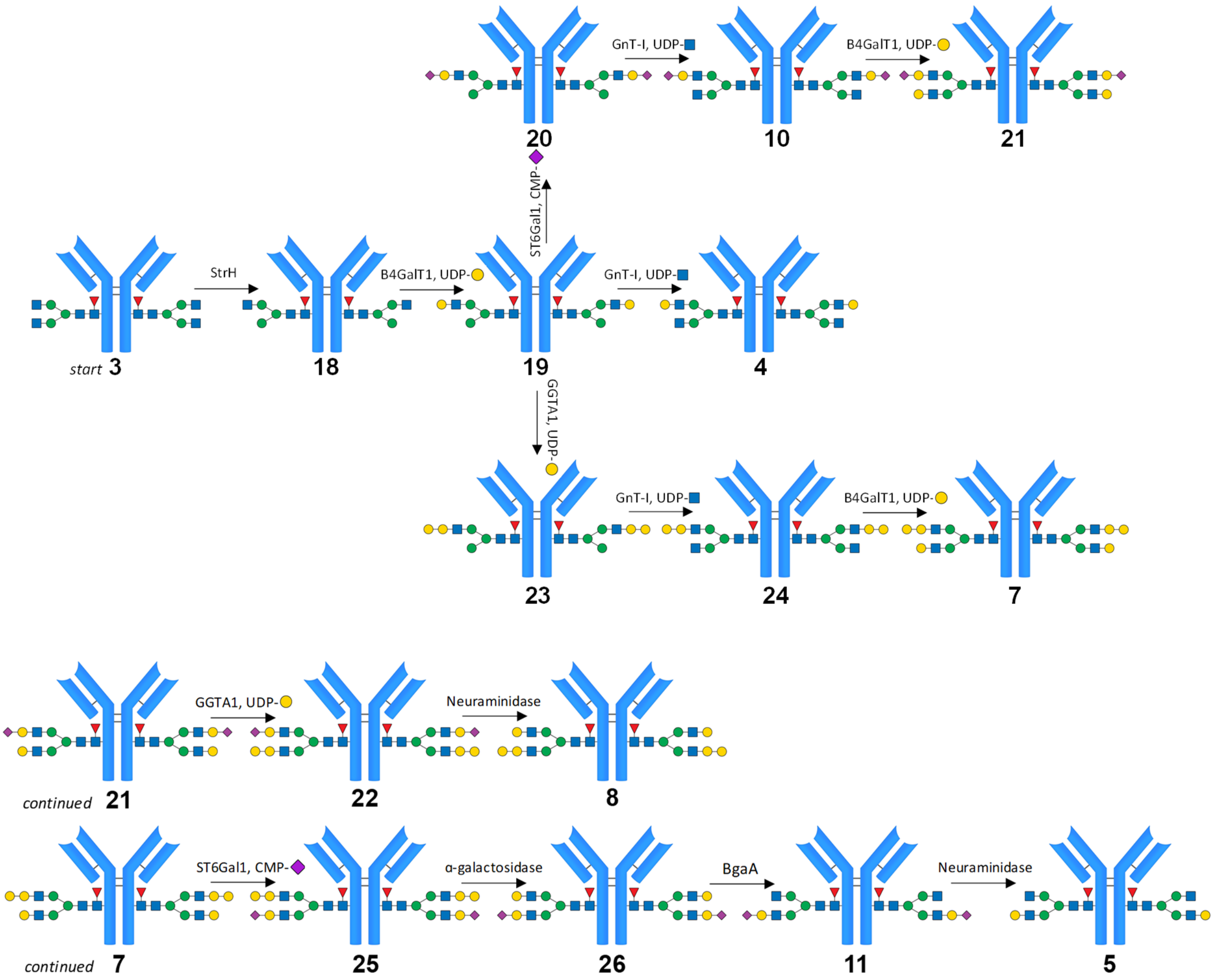

**Figure S11. Symmetrical glycoremodeling routes of infliximab and cetuximab**

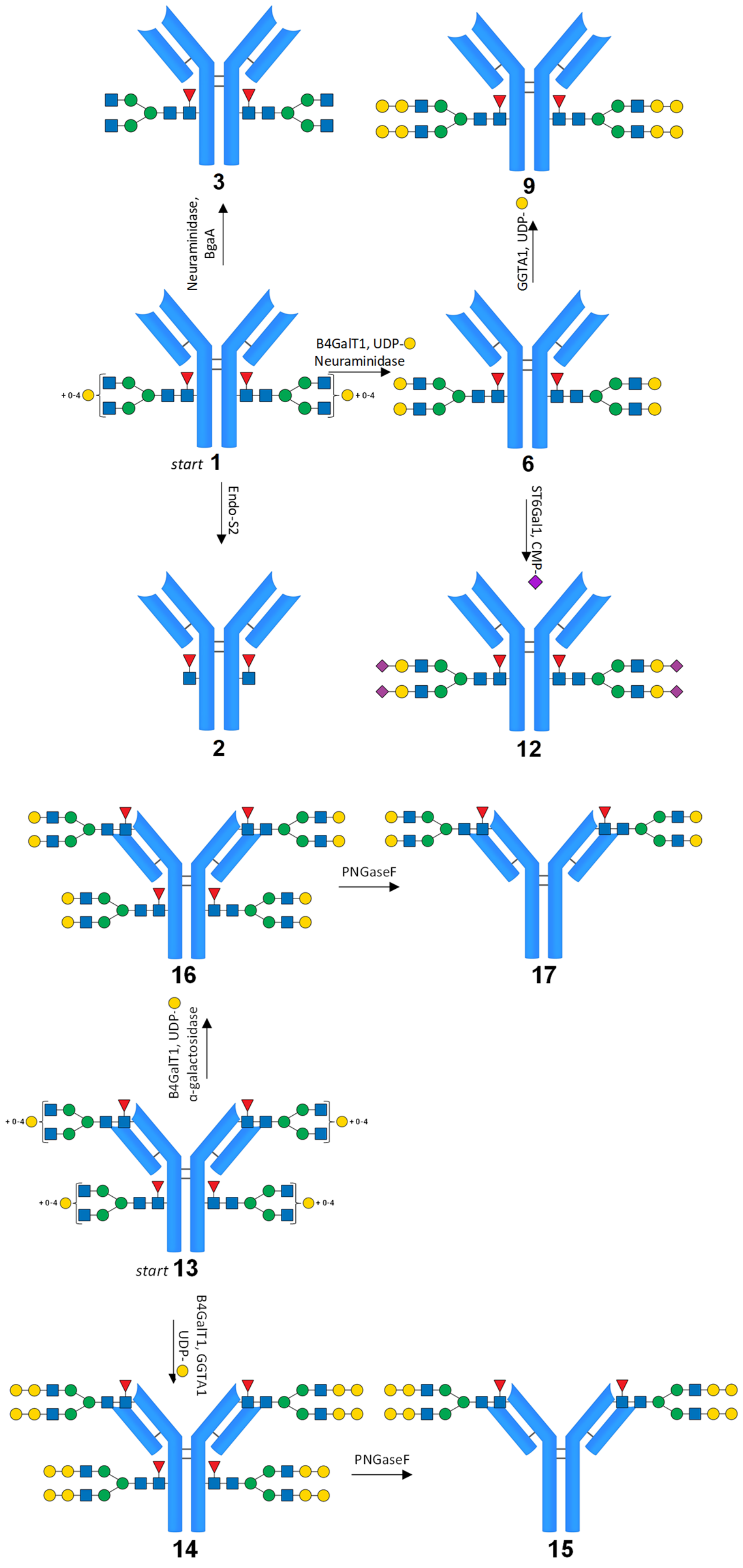


**Figure S11.** Symmetrical modification of the Fc glycan of infliximabs **1-3**, **6**, **9**, and **12** and symmetrical modification of the Fab and Fc glycan of cetuximab (**13-17**)

**Table S2. Theoretical masses versus observed masses infliximab**

| Infliximab | | lysine removed | | w/ lysine | |
|---|---|---|---|---|---|
| # | glycan | theoretical | observed | theoretical | observed |
| Fc | no glycan | 23754.7 | - | 23882.8 | - |
| 1 | A2G0F | 25202.7 | 25203.3 | 25330.8 | 25330.6 |
| | A2G1F | 25364.7 | 25364.8 | 25492.8 | 25492.4 |
| | A2G2F | 25526.7 | 25524.1 | 25654.8 | 25654.2 |
| 2 | GlcNAc-Fuc | 24104.7 | 24106.9 | 24232.8 | 24235.4 |
| 3 | A2F | 25202.7 | 25203.3 | 25330.8 | 25330.8 |
| 4 | A2G1F | 25365.7 | 25364.4 | 25493.8 | 25493.4 |
| 5 | A2G1F | 25365.7 | 25365.1 | 25493.8 | 25493.5 |
| 6 | A2G2F | 25526.7 | 25527.1 | 25654.8 | 25655.6 |
| 7 | A2G2G1F | 25689.7 | 25691.7 | 25817.8 | 25818.3 |
| 8 | A2G2G1F | 25689.7 | 25690.0 | 25817.8 | 25818.3 |
| 9 | A2G2G2F | 25850.7 | 25851.4 | 25978.8 | 25980.8 |
| 10 | A2G1S1F | 25657.7 | 25656.4 | 25785.8 | 25784.7 |
| 11 | A2G1S1F | 25657.7 | 25656.7 | 25785.8 | 25785.1 |
| 12 | A2G2S2F | 26110.7 | 26110.4 | 26238.8 | 26240.7 |
| Infliximab - intermediates | | | | | |
| 18 | A1F | 25000.7 | 25000.4 | 25128.8 | 25127.8 |
| 19 | A1G1F | 25162.7 | 25160.0 | 25290.8 | 25288.1 |
| 20 | A1G1S1F | 25455.7 | 25457.8 | 25583.8 | 25581.4 |
| 21 | A2G2S1F | 25819.7 | 25816.6 | 25947.8 | 25944.7 |
| 22 | A2G2G1S1F | 25981.7 | 25822.0 | 26109.8 | 25946.6 |
| 23 | A1G1G1F | 25324.7 | 25324.5 | 25452.8 | 25451.9 |
| 24 | A2G1G1F | 25527.7 | 25533.0 | 25655.8 | 25662.5 |
| 25 | A2G2G1S1F | 25981.7 | 25981.2 | 26109.8 | 26111.8 |
| 26 | A2G2S1F | 25819.7 | 25816.7 | 25947.8 | 25944.8 |

Infliximab Fc sequence after IdeS cleavage

Molecular weight; 23882.83 g/mol | C-term lysine removed; 23754.66 g/mol

```
                                                       230        240

                                                                   GP

       250        260        270        280        290        300

SVFLFPPKPK DTLMISRTPE VTCVVVDVSH EDPEVKFNWY VDGVEVHNAK TKPREEQYNS

       310        320        330        340        350        360

TYRVVSVLTV LHQDWLNGKE YKCKVSNKAL PAPIEKTISK AKGQPREPQV YTLPPSRDEL

       370        380        390        400        410        420

TKNQVSLTCL VKGFYPSDIA VEWESNGQPE NNYKTTPPVL DSDGSFFLYS KLTVDKSRWQ

       430        440

QGNVFSCSVM HEALHNHYTQ KSLSLSPGK
```

N-glycosylation site Non-aligned residues infliximab/cetuximab

**Table S3. Theoretical masses versus observed masses cetuximab**

| Cetuximab | | lysine removed | | w/ lysine | |
|---|---|---|---|---|---|
| # | glycan | theoretical | observed | theoretical | observed |
| Fc | no glycan | 23790.8 | - | 23918.9 | - |
| 13 | A2G0F | 25238.8 | 25234.3 | 25366.9 | 25363.3 |
| | A2G1F | 25400.8 | 25396.6 | 25528.9 | 25524.1 |
| | A2G2F | 25562.8 | 25557.2 | 25690.9 | 25684.3 |
| 14 | A2G2G2F | 25886.8 | 25888.3 | 26014.9 | 26012.0 |
| 15 | no glycan | 23790.8 | 23790.1 | 23918.9 | 23922.2 |
| 16 | A2G2F | 25562.8 | 25559.5 | 25690.9 | 25693.5 |
| 17 | no glycan | 23790.8 | 23793.1 | 23918.9 | 23908.4 |
| Cetuximab | | | | | |
| # | glycan | theoretical | observed | | |
| Fab | no glycan | 25470.5 | - | | |
| 13 | A2G0F | 26900.5 | - | | |
| | A2G1F | 27062.5 | 27060.4 | | |
| | A2G2F | 27224.5 | 27221.2 | | |
| | A2G2G1F | 27386.5 | 27382.2 | | |
| | A2G2G2F | 27548.5 | 27544.6 | | |
| 14 | A2G2G2F | 27548.5 | 27545.1 | | |
| 15 | A2G2G2F | 27548.5 | 27545.0 | | |
| 16 | A2G2F | 27224.5 | 27220.7 | | |
| 17 | A2G2F | 27224.5 | 27221.3 | | |

Cetuximab Fc sequence after IdeS cleavage and subsequent DTT treatment

Molecular weight; 23918.93 g/mol | C-term lysine removed; 23790.76 g/mol

```
                                                   230        240

                                                               GP

       250        260        270        280        290        300

SVFLFPPKPK DTLMISRTPE VTCVVVDVSH EDPEVKFNWY VDGVEVHNAK TKPREEQYNS

       310        320        330        340        350        360

TYRVVSVLTV LHQDWLNGKE YKCKVSNKAL PAPIEKTISK AKGQPREPQV YTLPPSREEM

       370        380        390        400        410        420

TKNQVSLTCL VKGFYPSDIA VEWESNGQPE NNYKTTPPVL DSDGSFFLYS KLTVDKSRWQ

       430        440

QGNVFSCSVM HEALHNHYTQ KSLSLSPGK
```

Cetuximab Fab sequence after IdeS cleavage and subsequent DTT treatment

Molecular weight; 25470.48 g/mol

```
        10         20         30         40         50         60
QVQLKQSGPG LVQPSQSLSI TCTVSGFSLT NYGVHWVRQS PGKGLEWLGV IWSGGNTDYN
        70         80         90        100        110        120
TPFTSRLSIN KDNSKSQVFF KMNSLQSNDT AIYYCARALT YYDYEFAYWG QGTLVTVSAA
       130        140        150        160        170        180
STKGPSVFPL APSSKSTSGG TAALGCLVKD YFPEPVTVSW NSGALTSGVH TFPAVLQSSG
       190        200        210        220        230
LYSLSSVVTV PSSSLGTQTY ICNVNHKPSN TKVDKRVEPK SCDKTHTCPP CPAPELLG
```

N-glycosylation site Non-aligned residues infliximab/cetuximab

**Figure S12. Deconvoluted MS spectrum mAb 1**

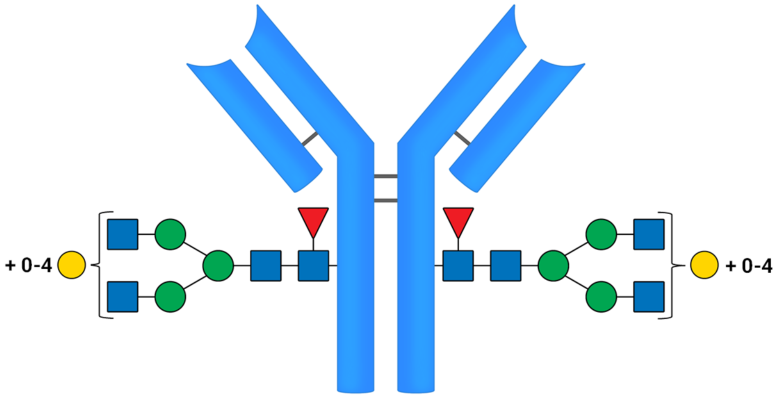


**1**

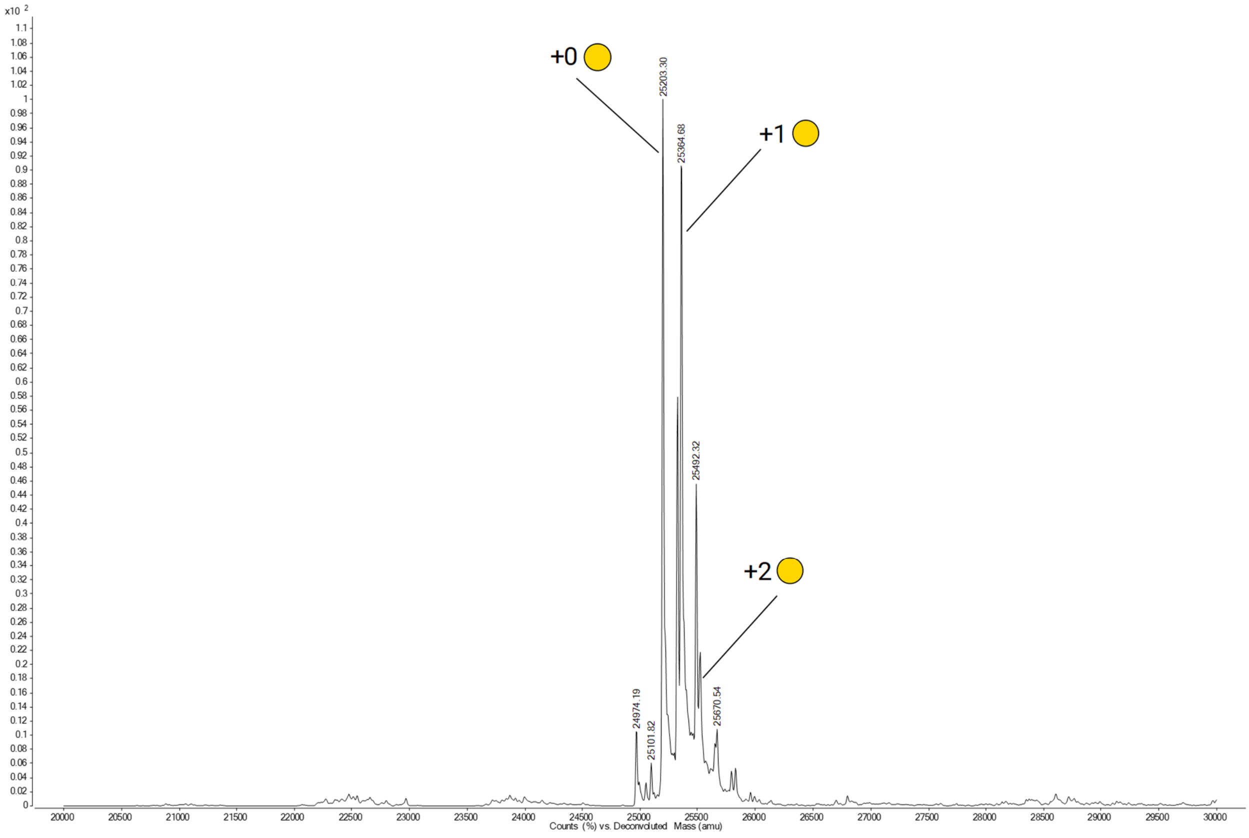

**Figure S13. Deconvoluted MS spectrum mAb 2**

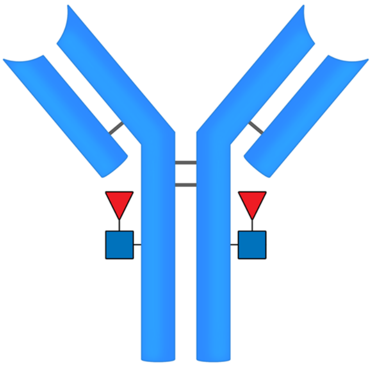

2

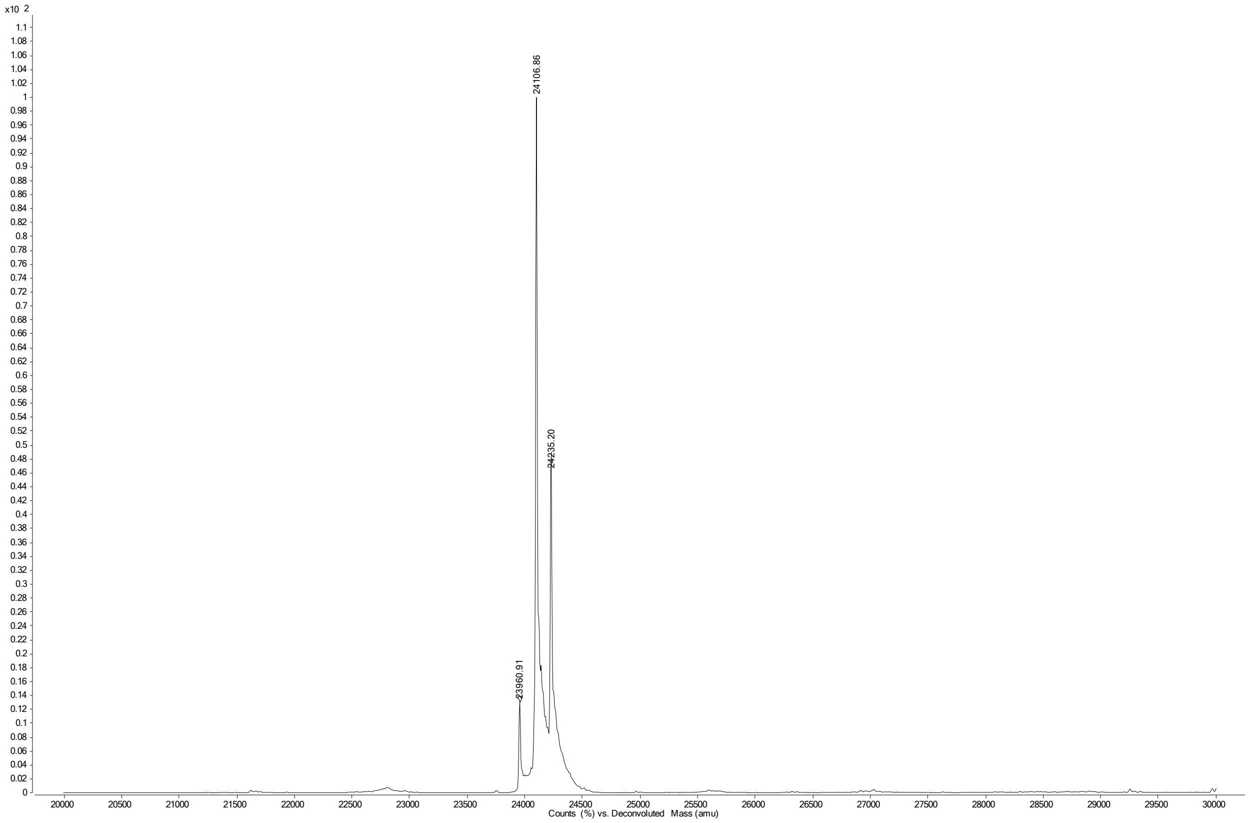

**Figure S14. Deconvoluted MS spectrum mAb 3**

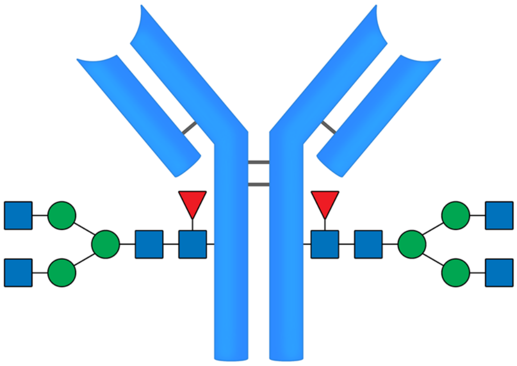

**3**

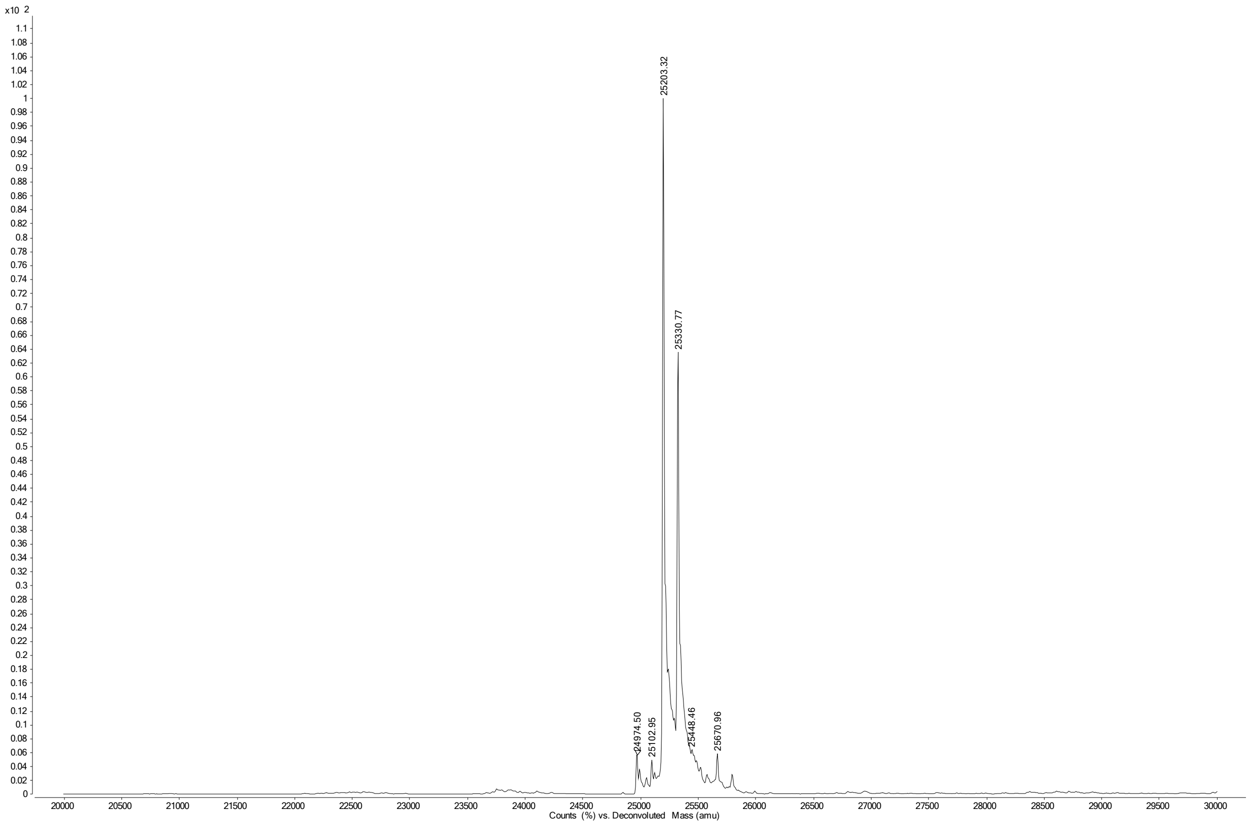

**Figure S15. Deconvoluted MS spectrum mAb 4**

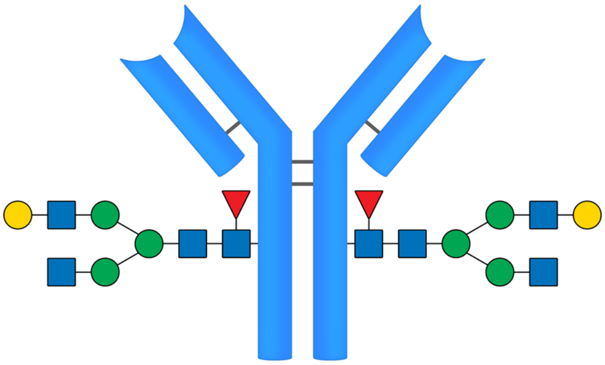

4

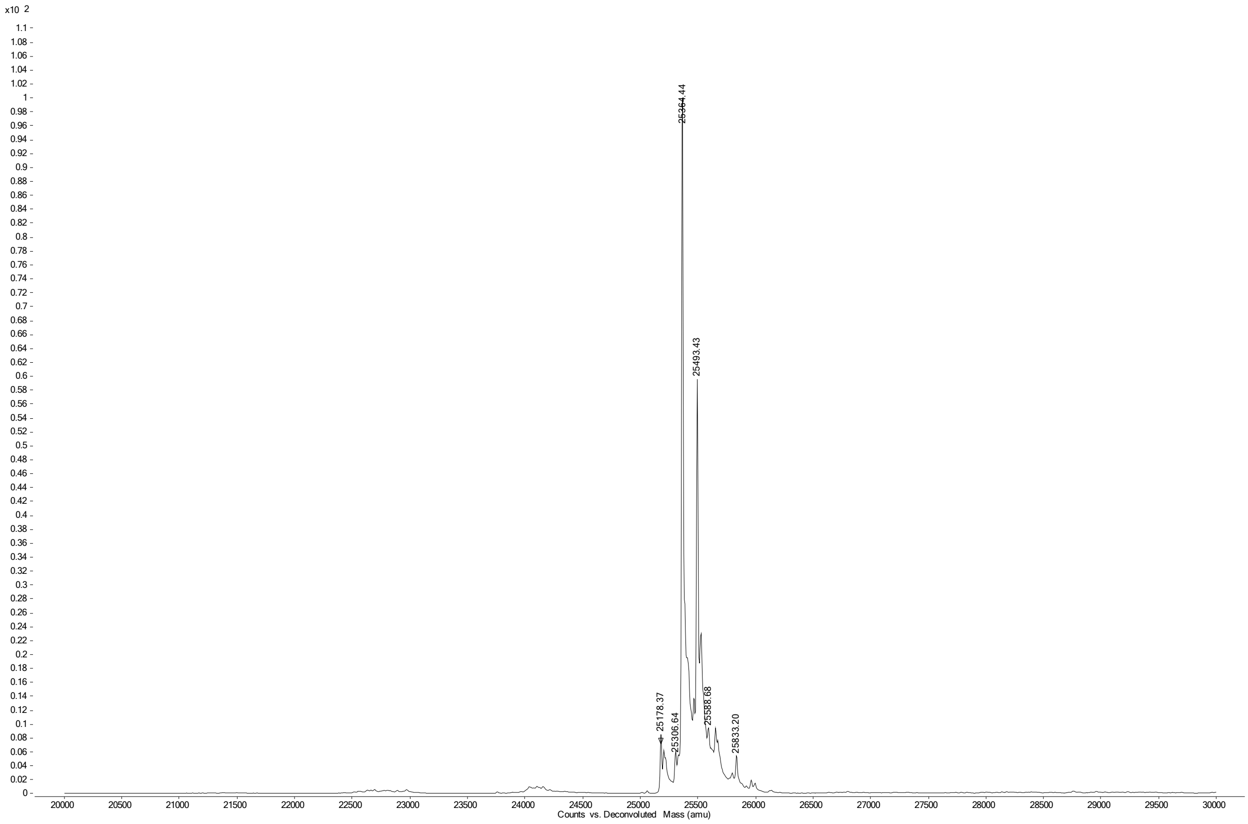

**Figure S16. Deconvoluted MS spectrum mAb 5**

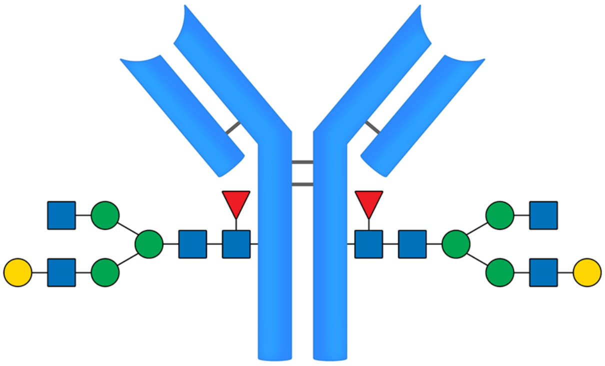

5

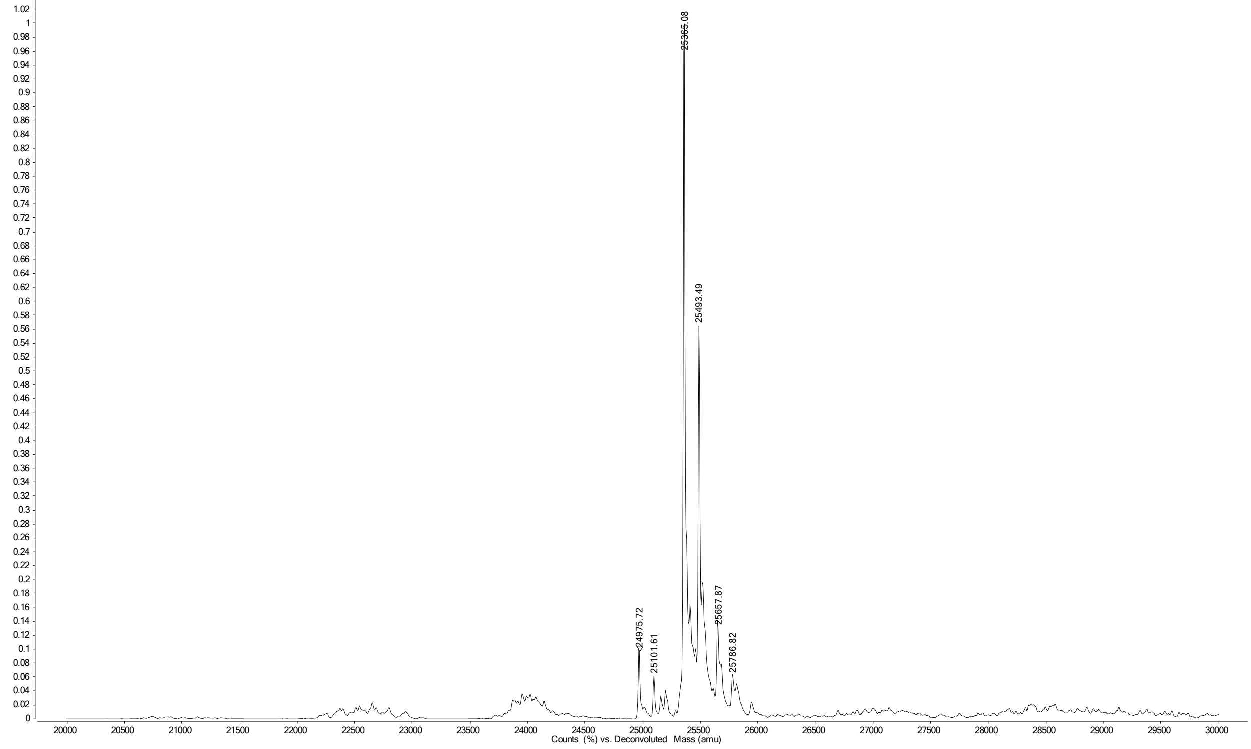

**Figure S17. Deconvoluted MS spectrum mAb 6**

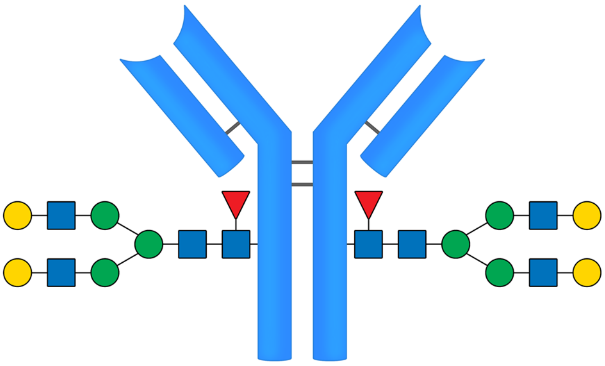

**6**

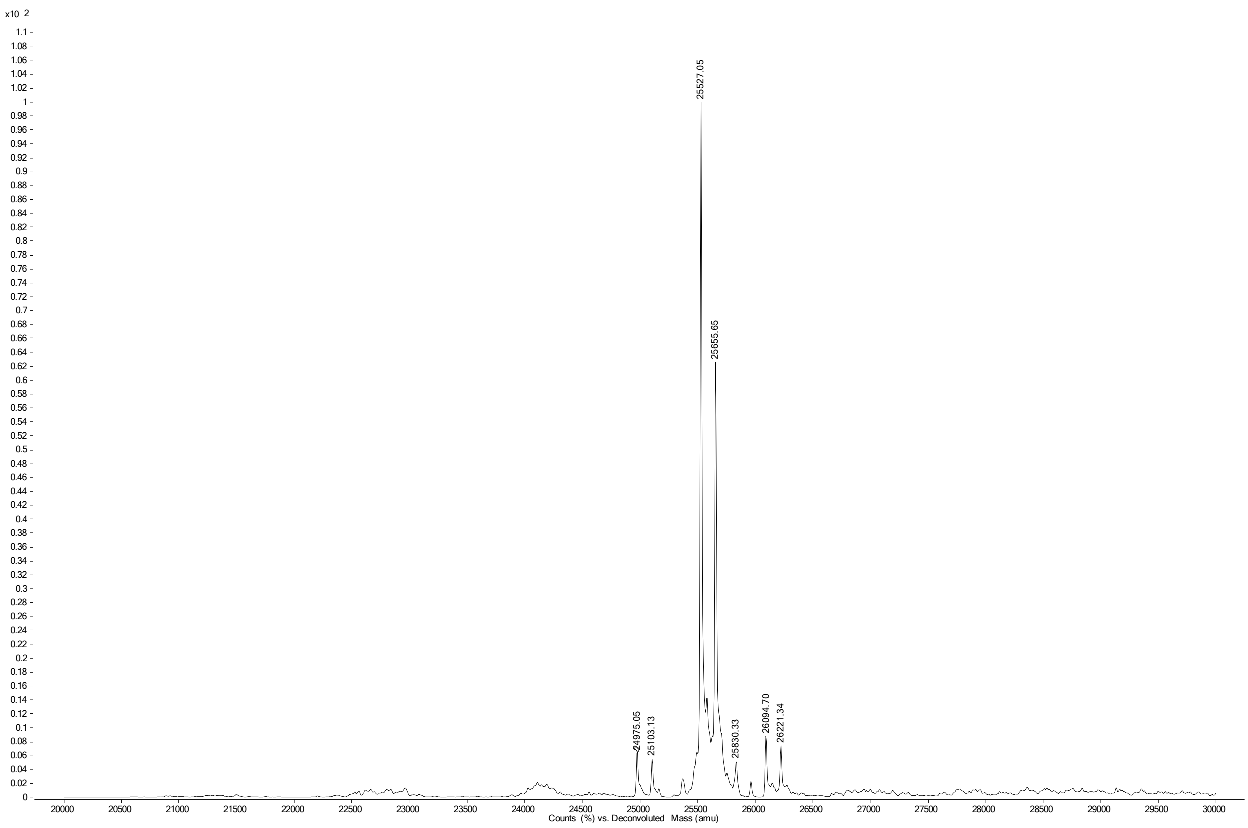

**Figure S18. Deconvoluted MS spectrum mAb 7**

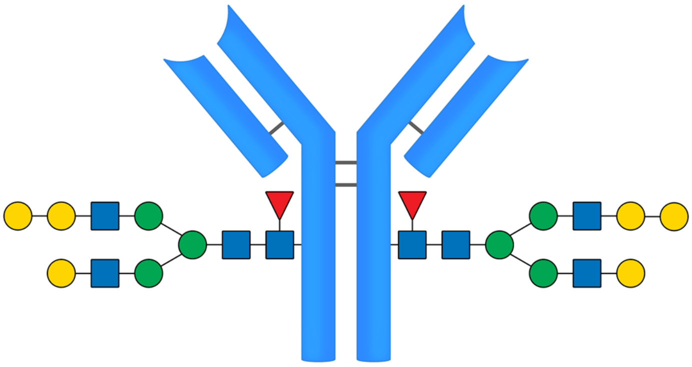

7

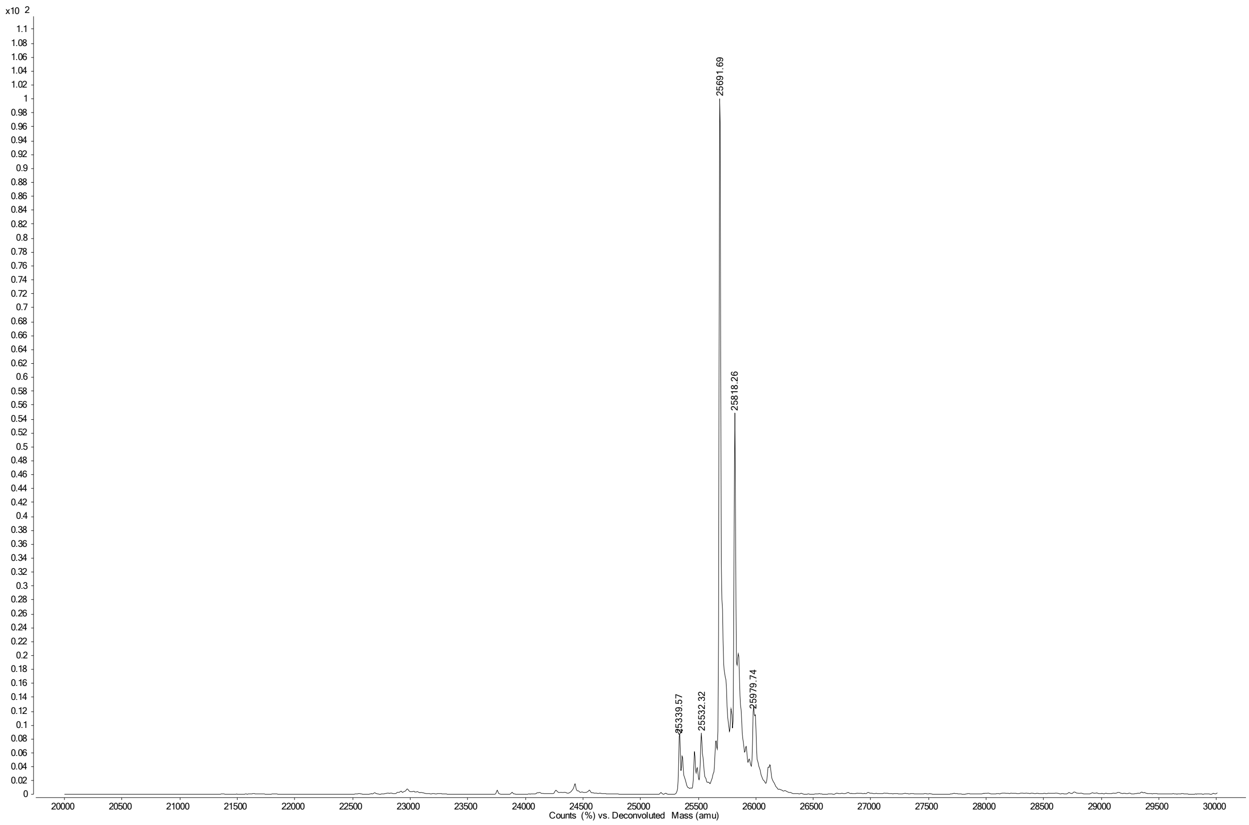

**Figure S19. Deconvoluted MS spectrum mAb 8**

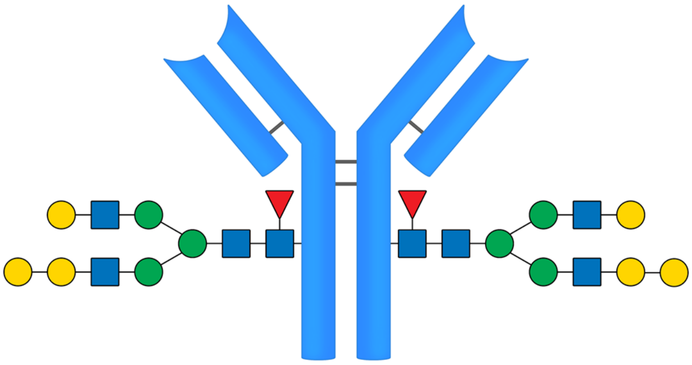

**8**

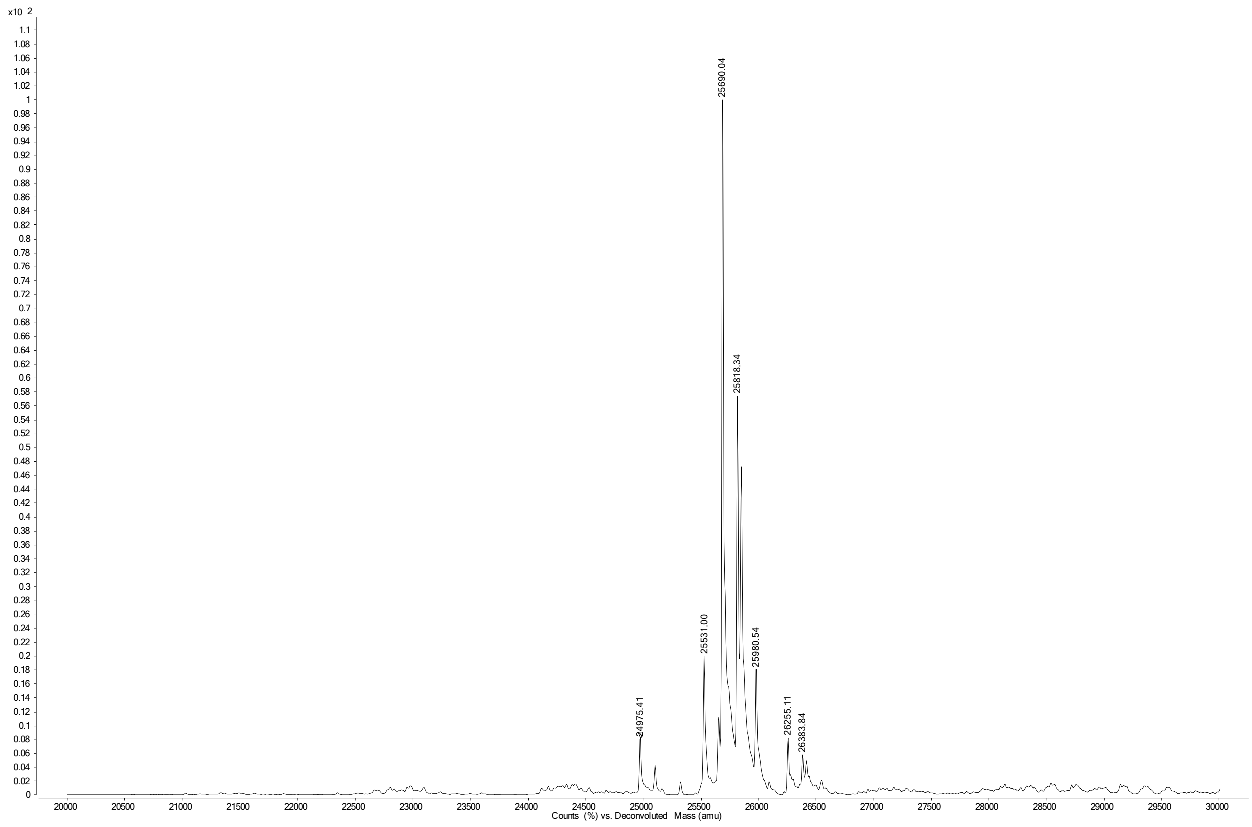

**Figure S20. Deconvoluted MS spectrum mAb 9**

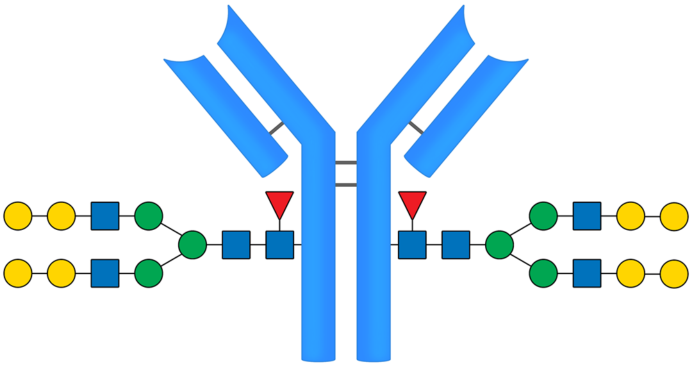

**9**

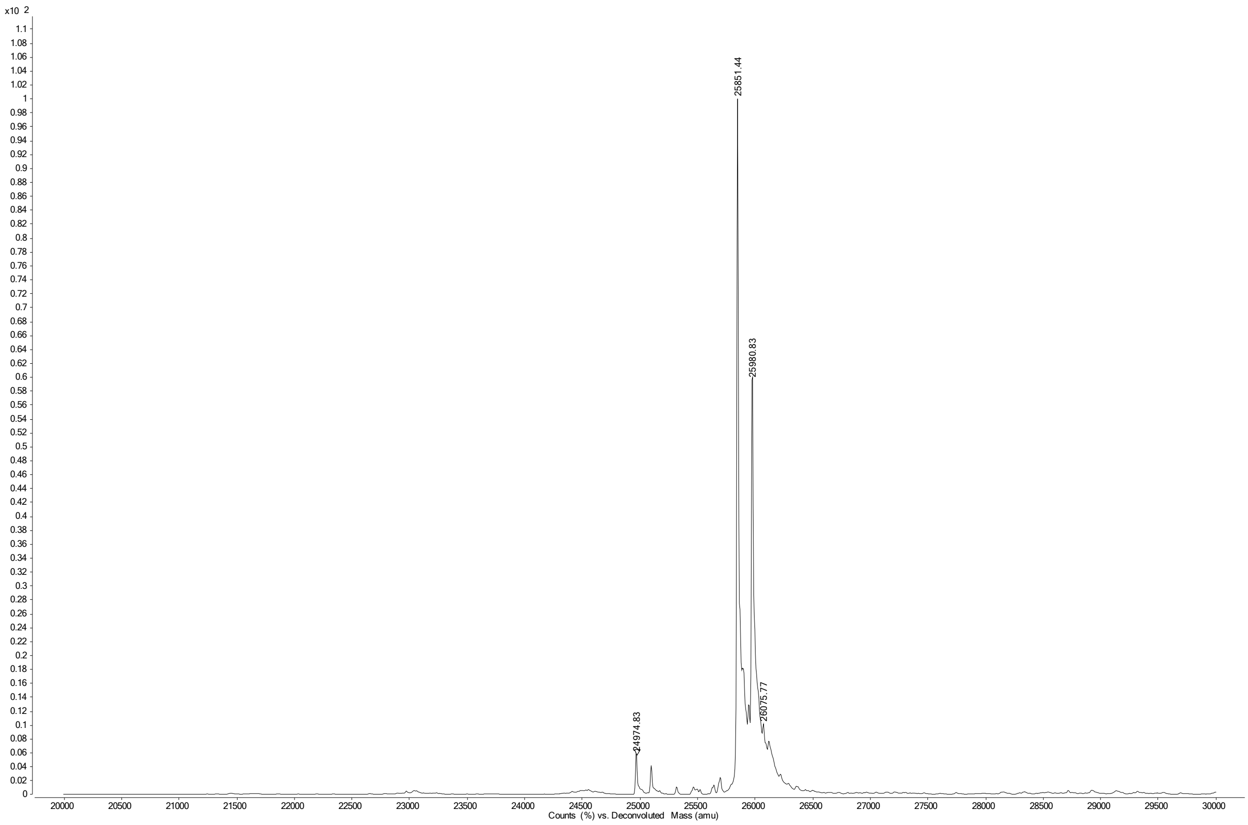

**Figure S21. Deconvoluted MS spectrum mAb 10**

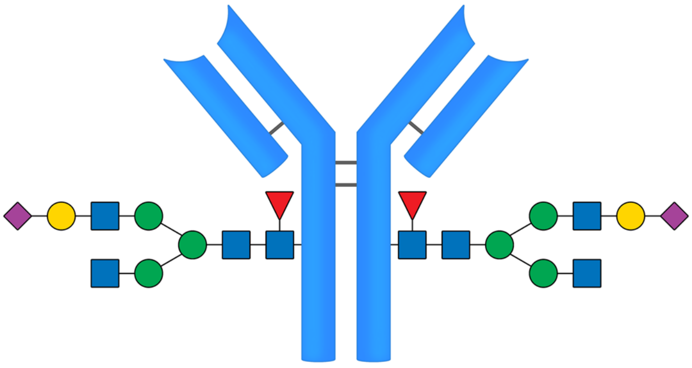

**10**

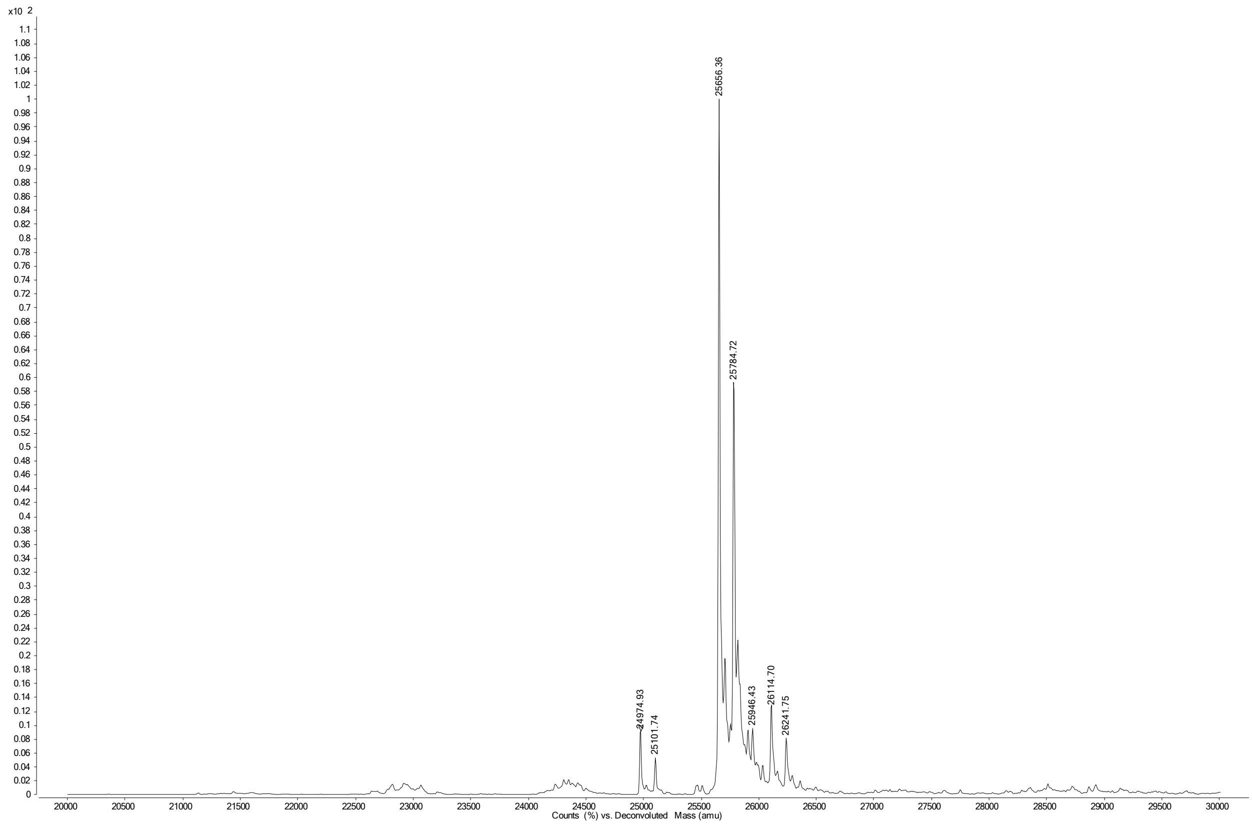

**Figure S22. Deconvoluted MS spectrum mAb 11**

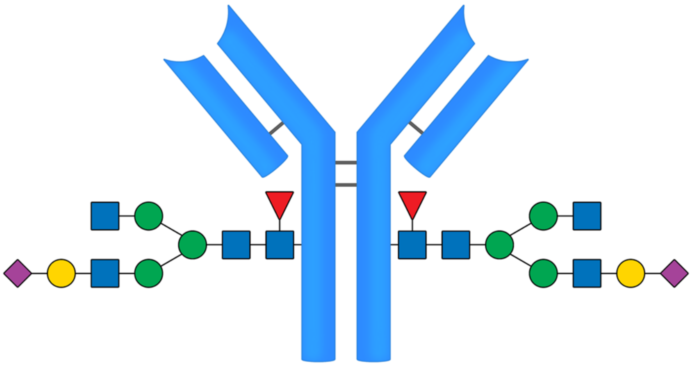

11

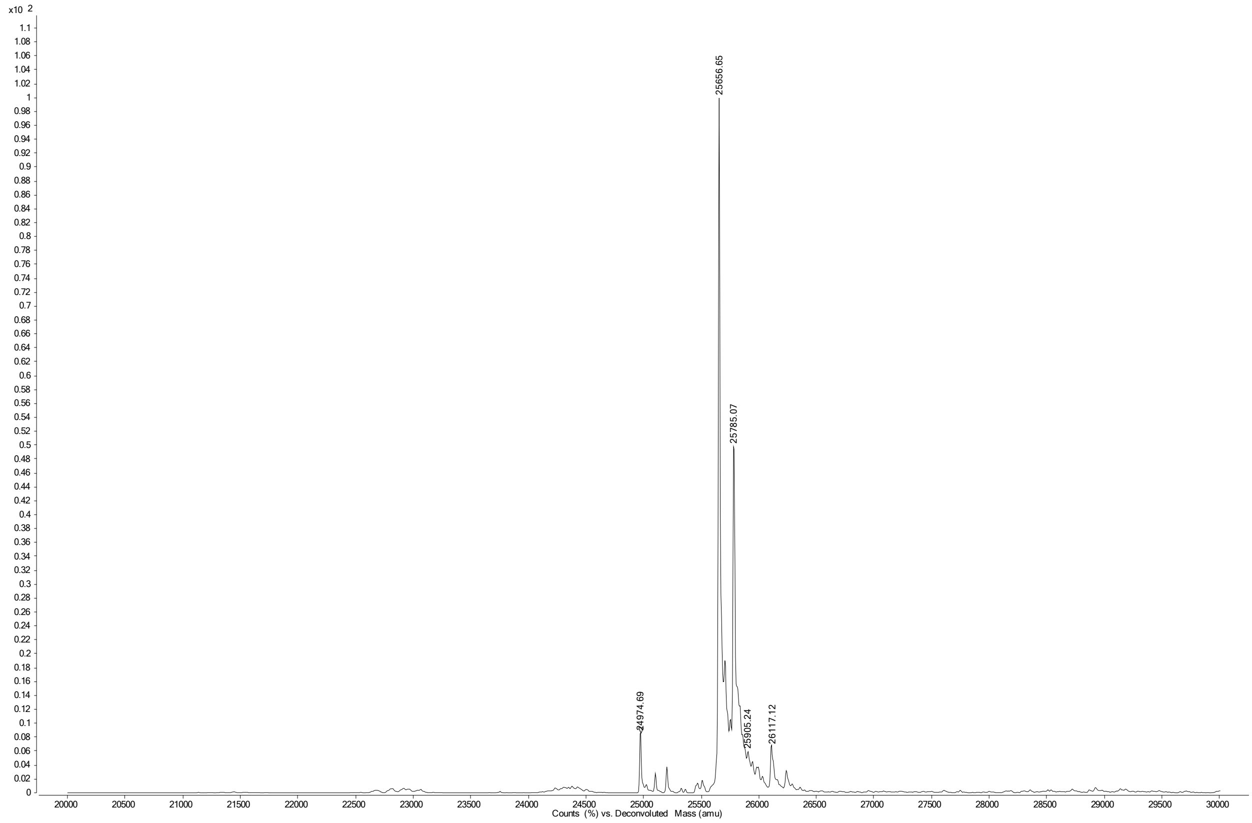

**Figure S23. Deconvoluted MS spectrum mAb 12**

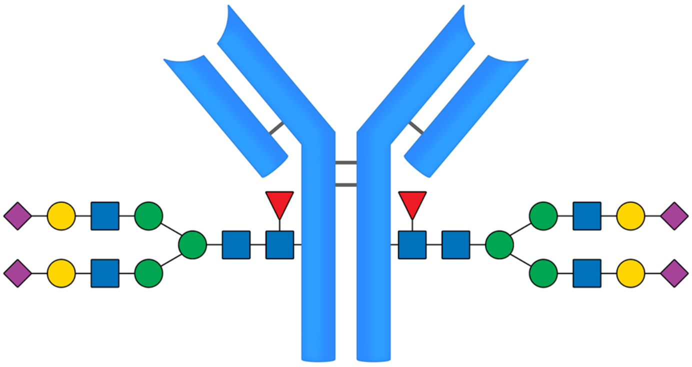

12

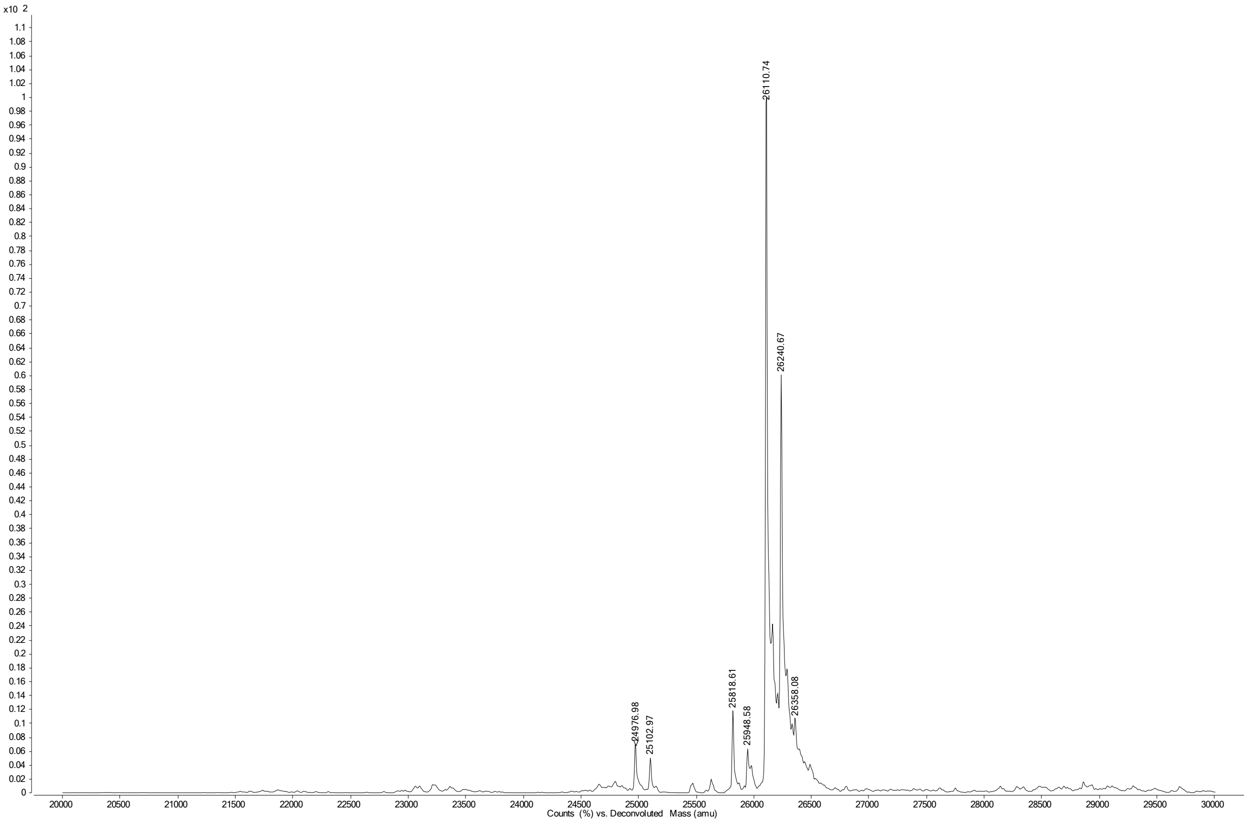

**Figure S24. Deconvoluted MS spectrum mAb 13**

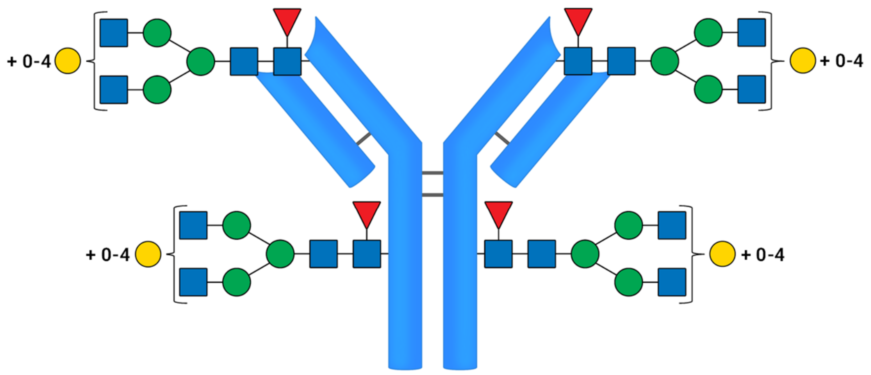


**13**

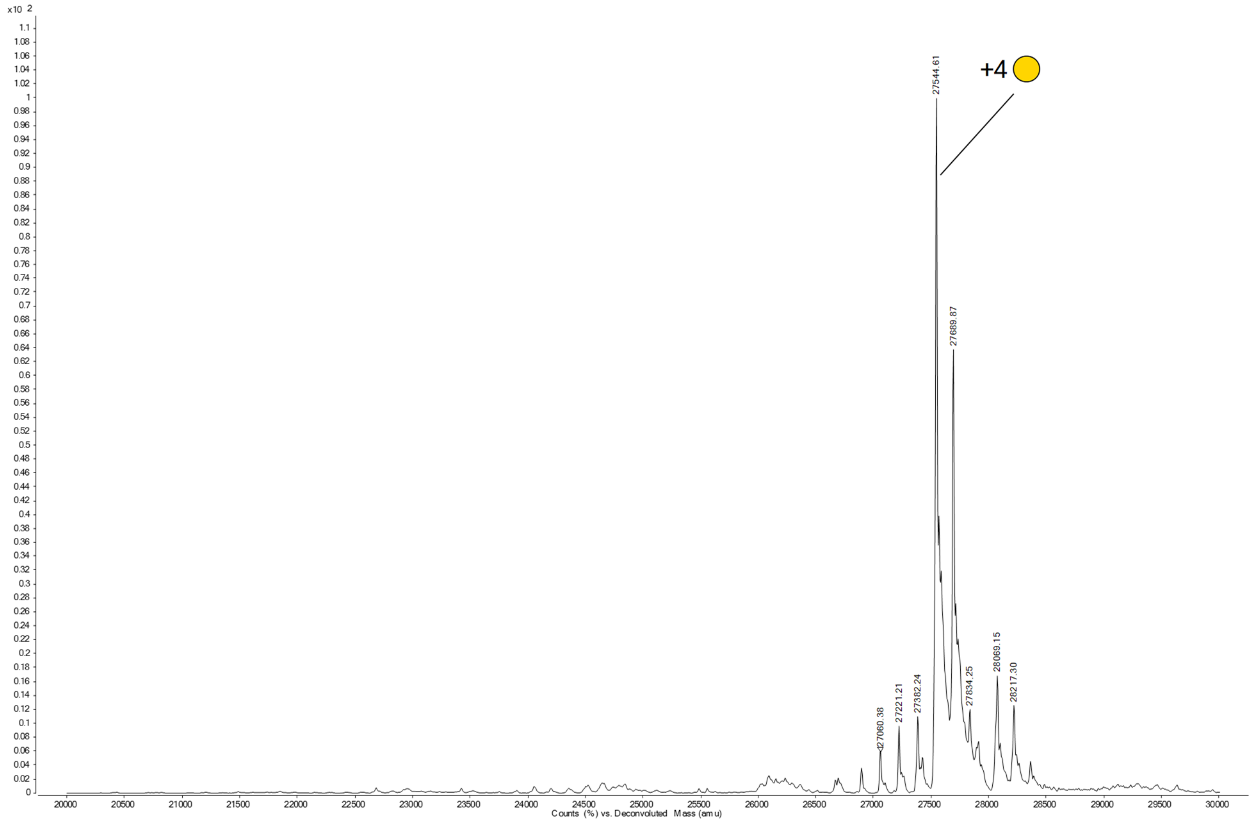


Single Fab domain

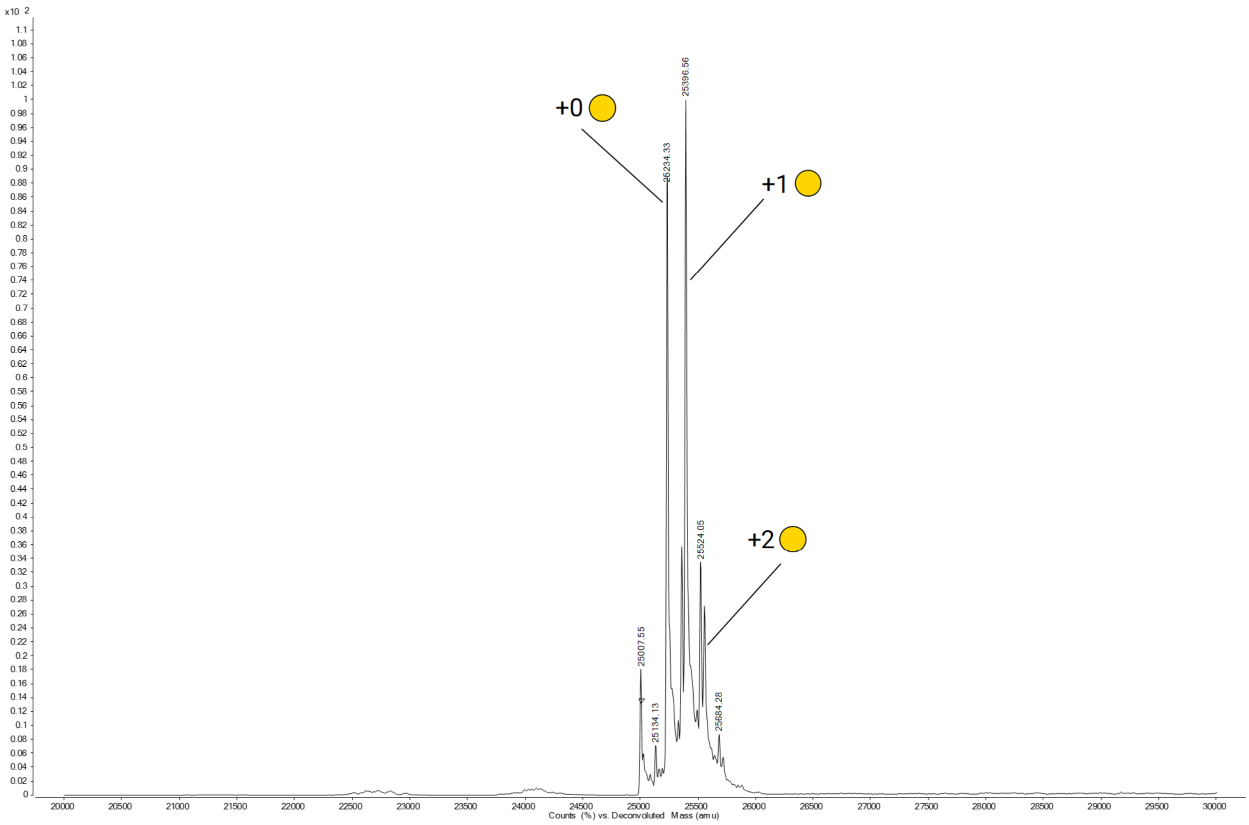


Single Fc domain

**Figure S25. Deconvoluted MS spectrum mAb 14**

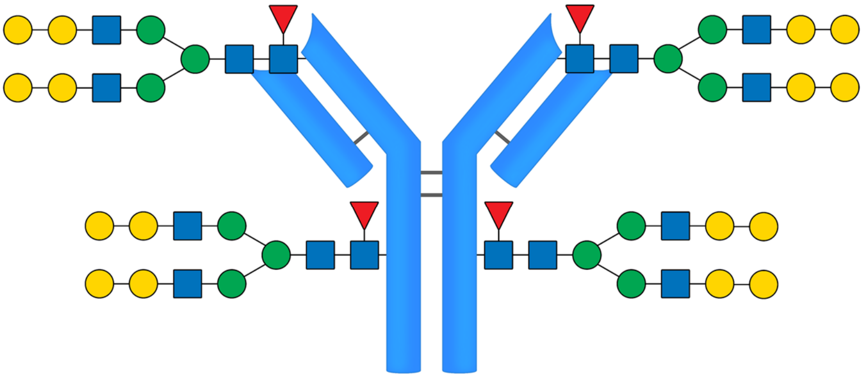

14

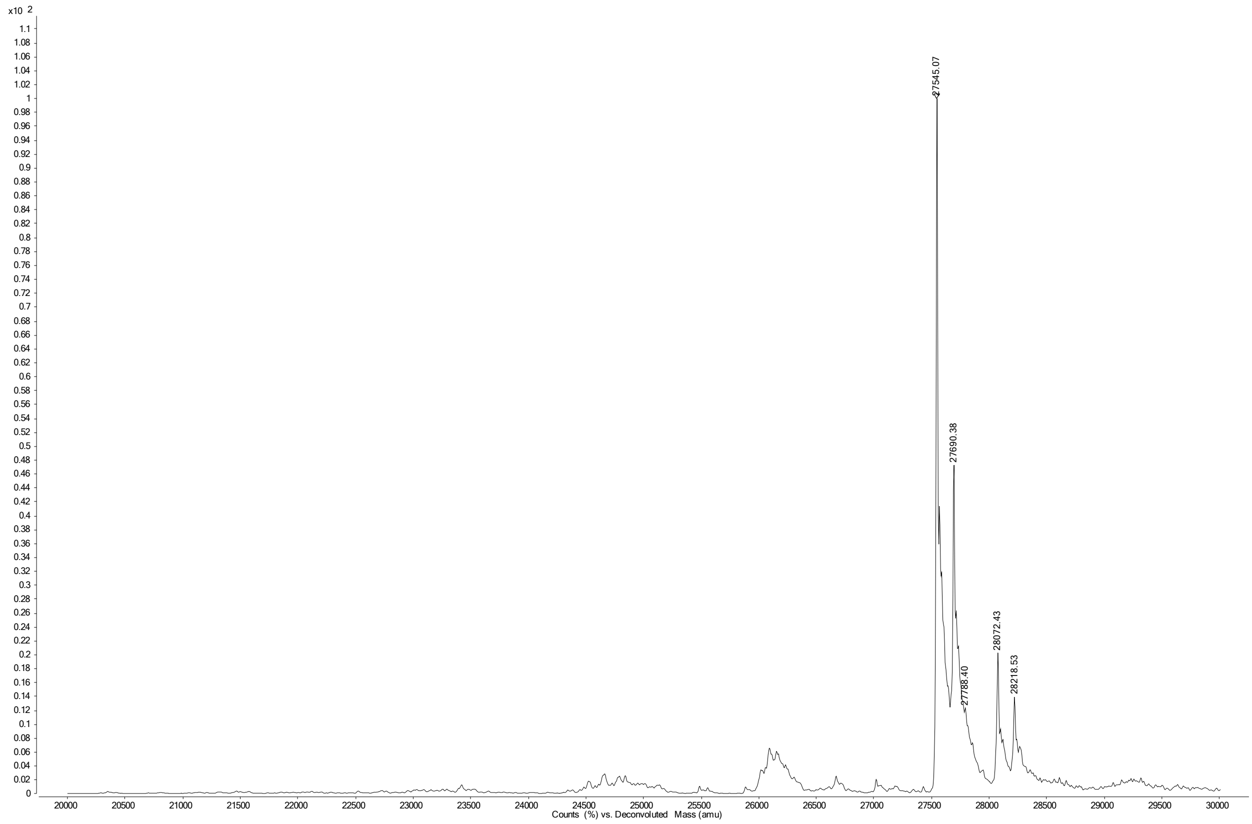


Single Fab domain

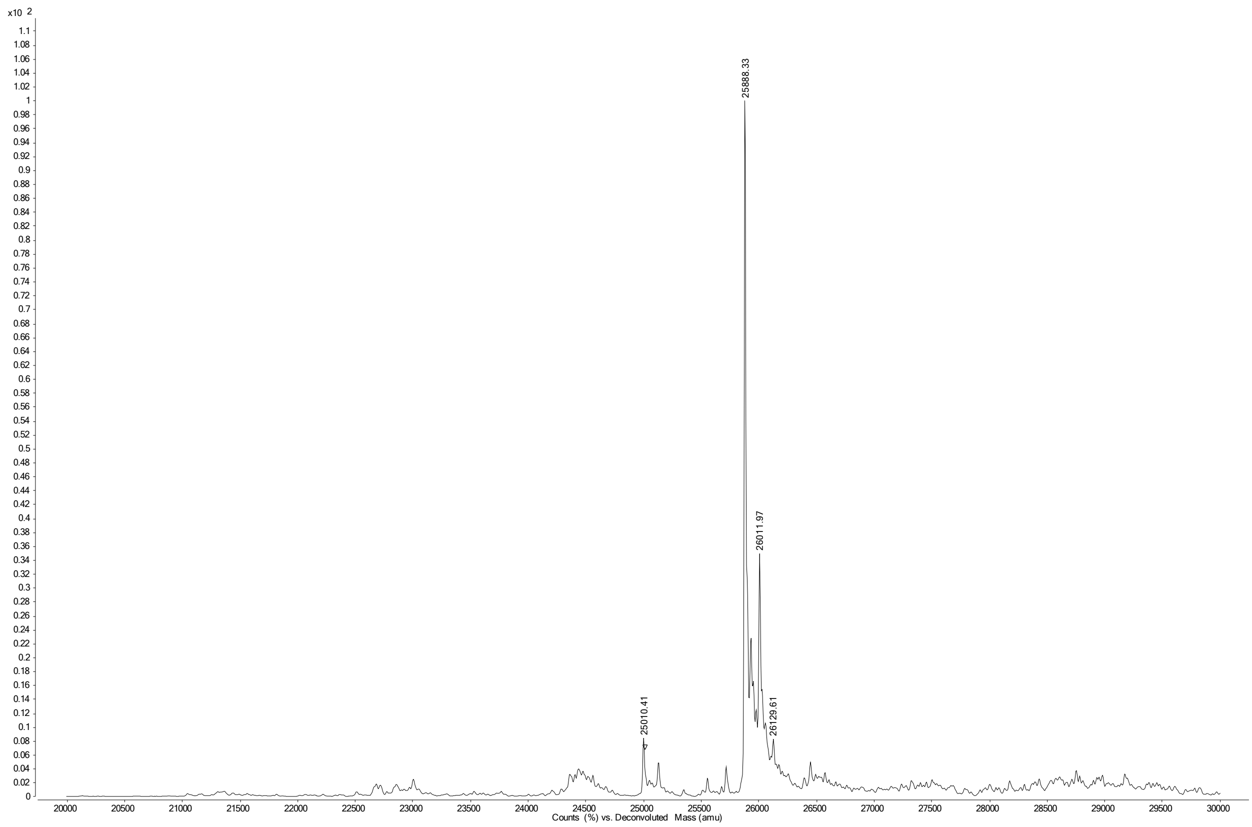


Single Fc domain

**Figure S26. Deconvoluted MS spectrum mAb 15**

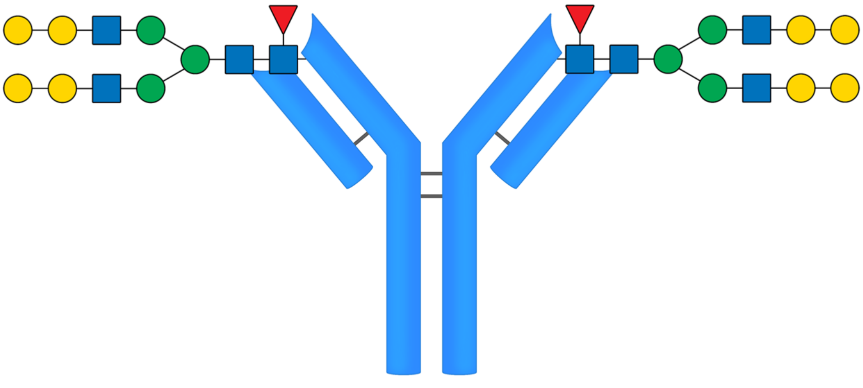


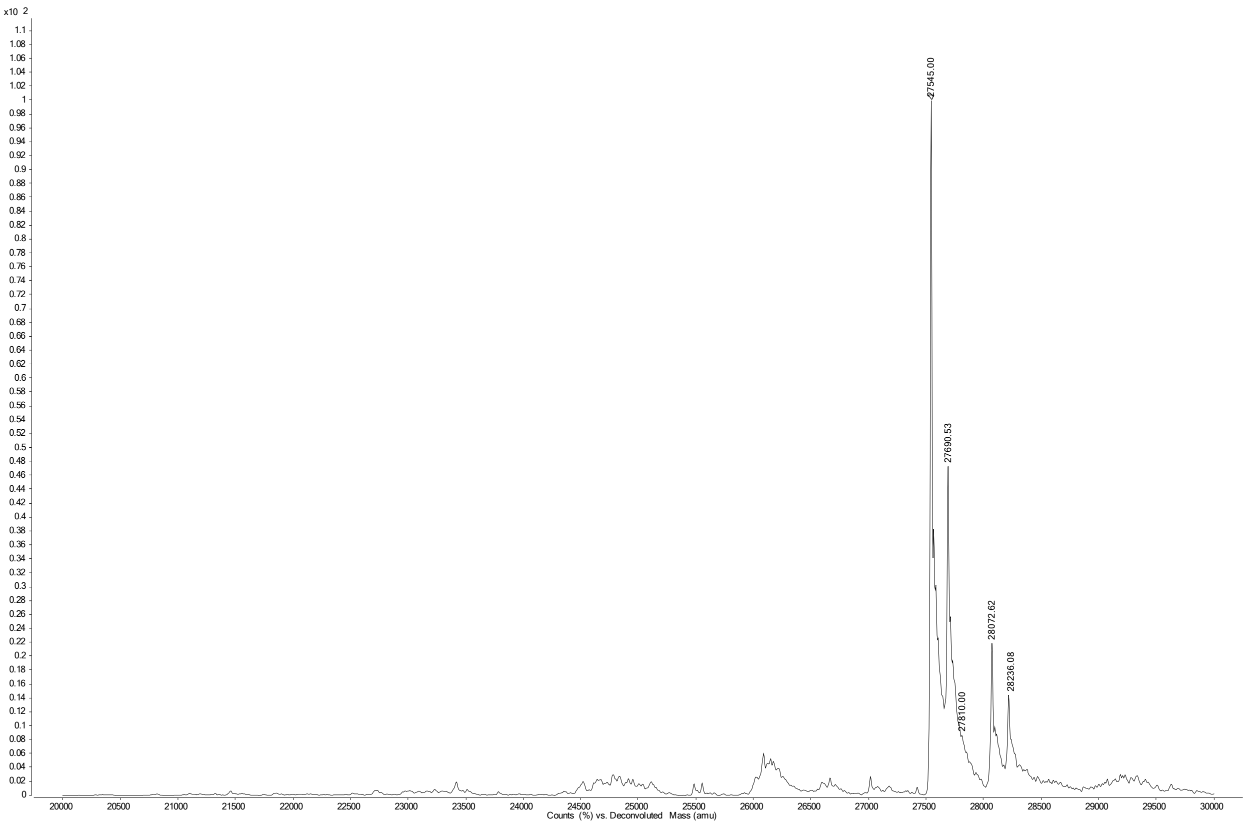


Single Fab domain

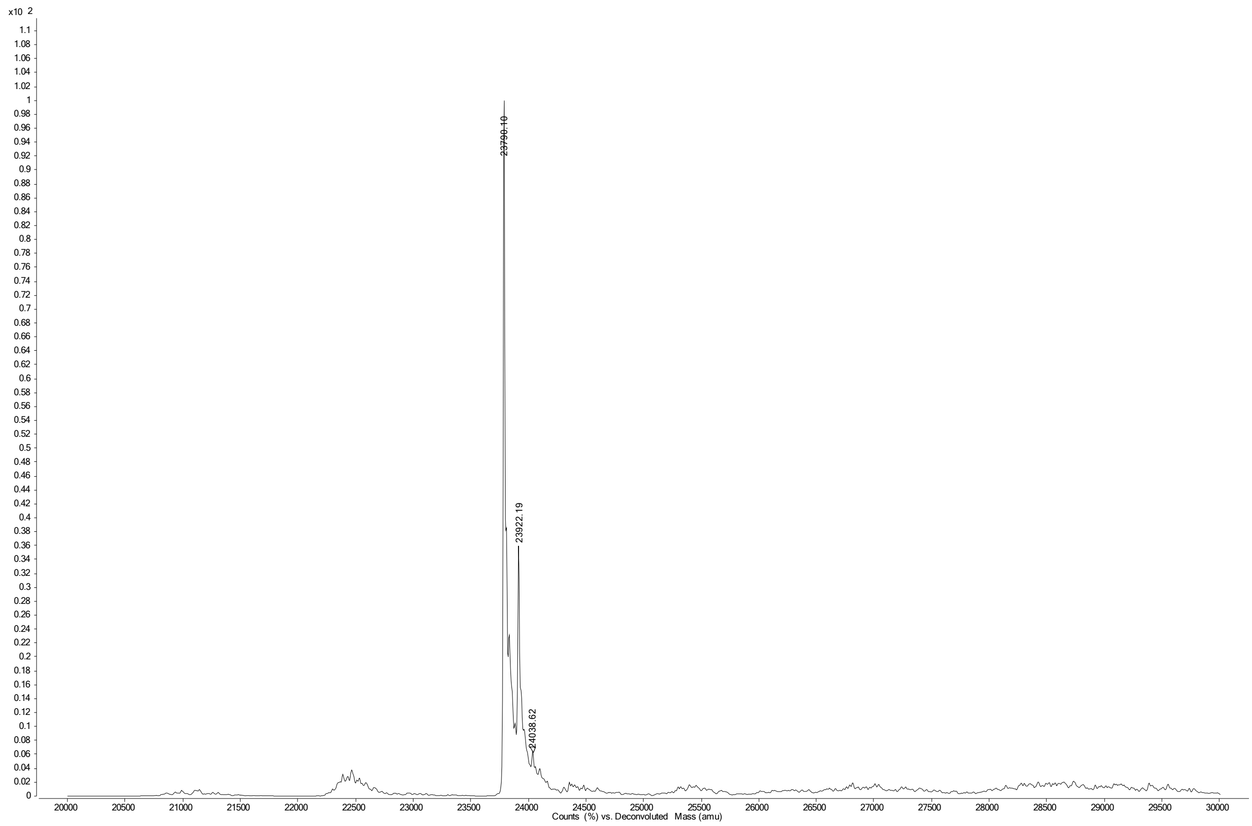


Single Fc domain

**Figure S27. Deconvoluted MS spectrum mAb 16**

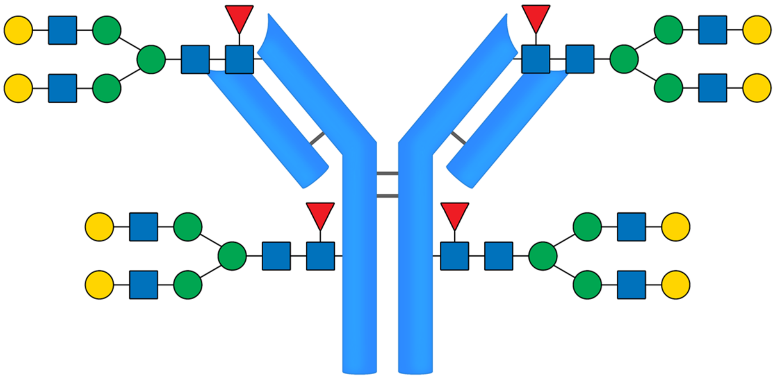

16

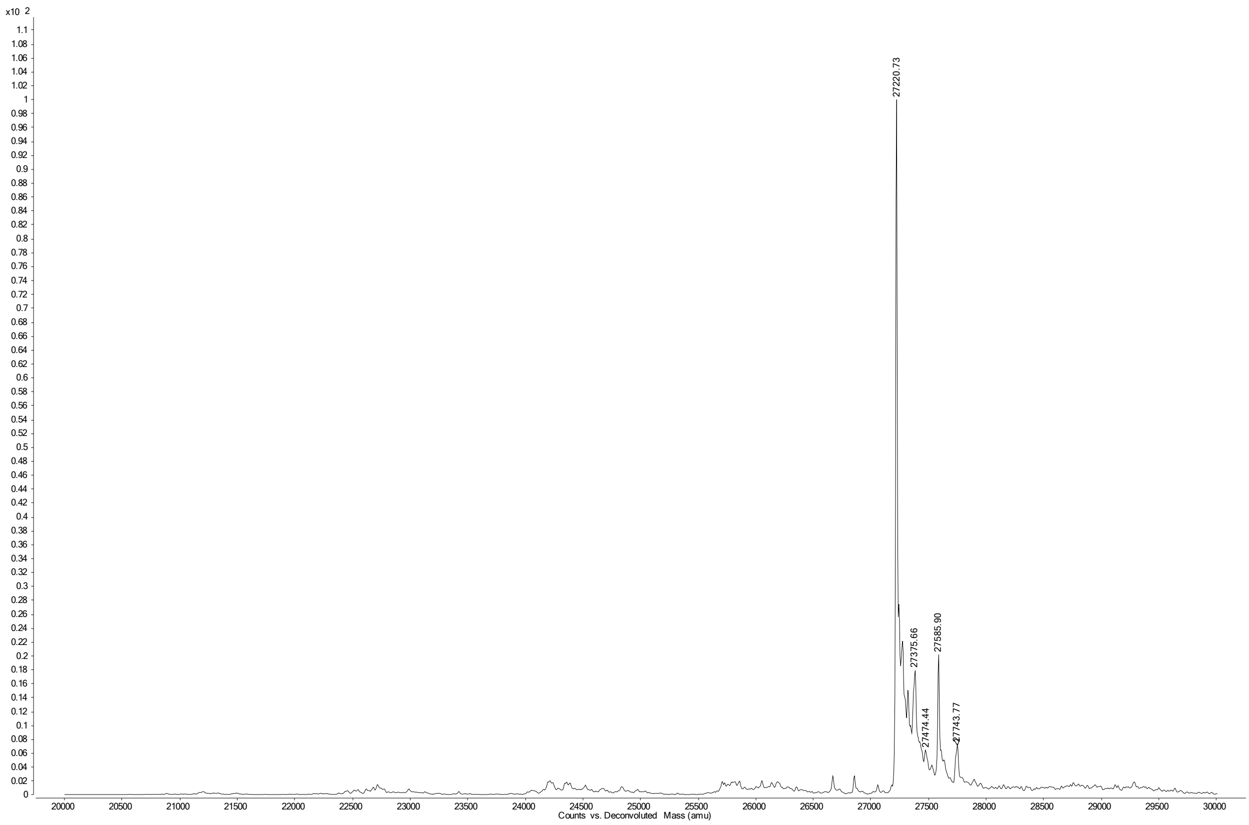


Single Fab domain

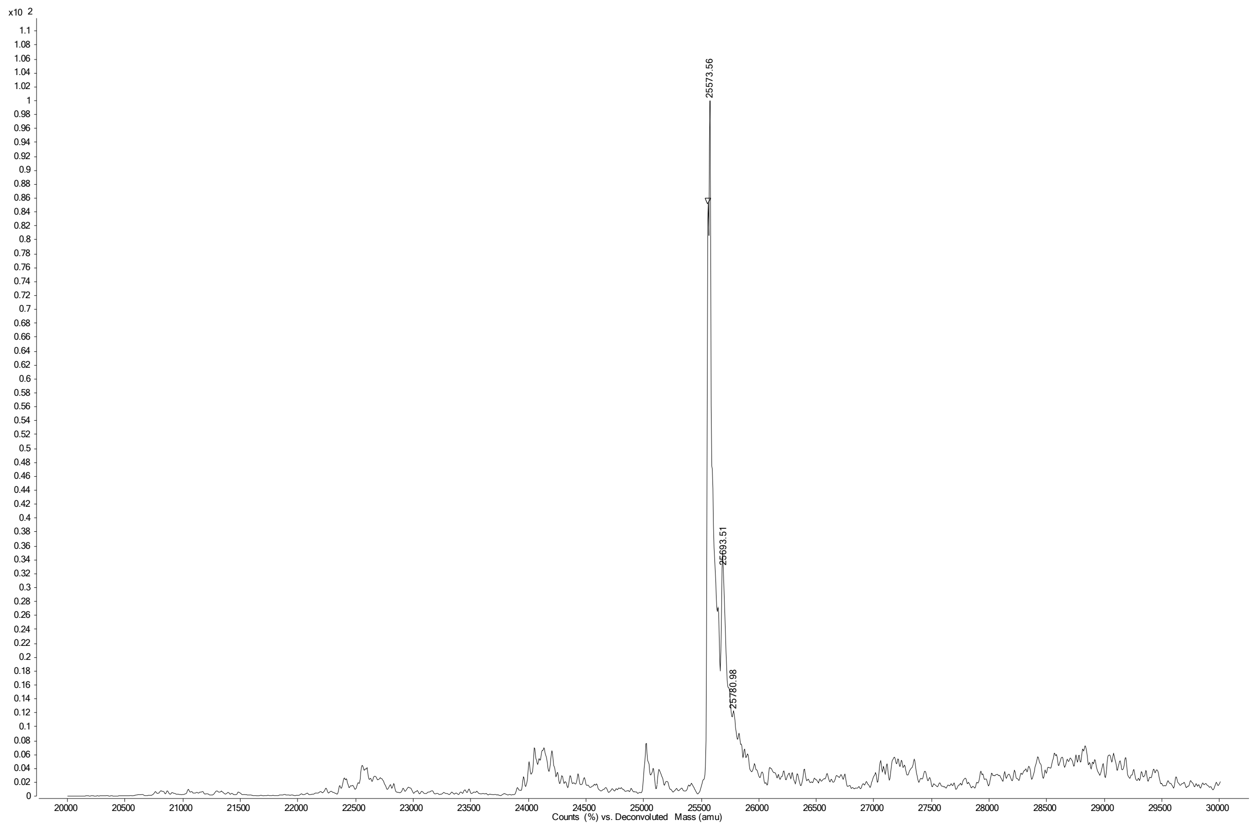


Single Fc domain

**Figure S28. Deconvoluted MS spectrum mAb 17**

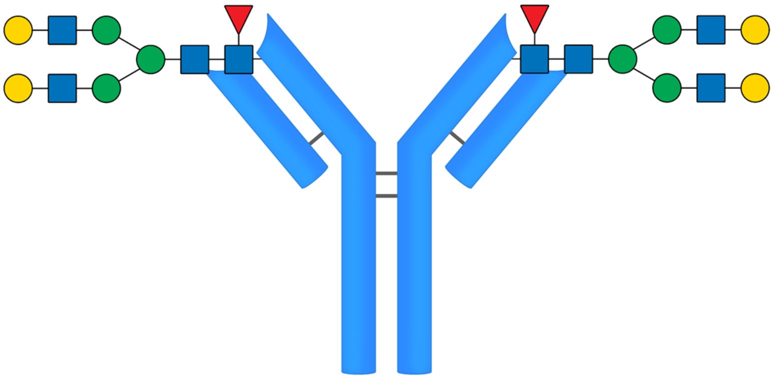

17

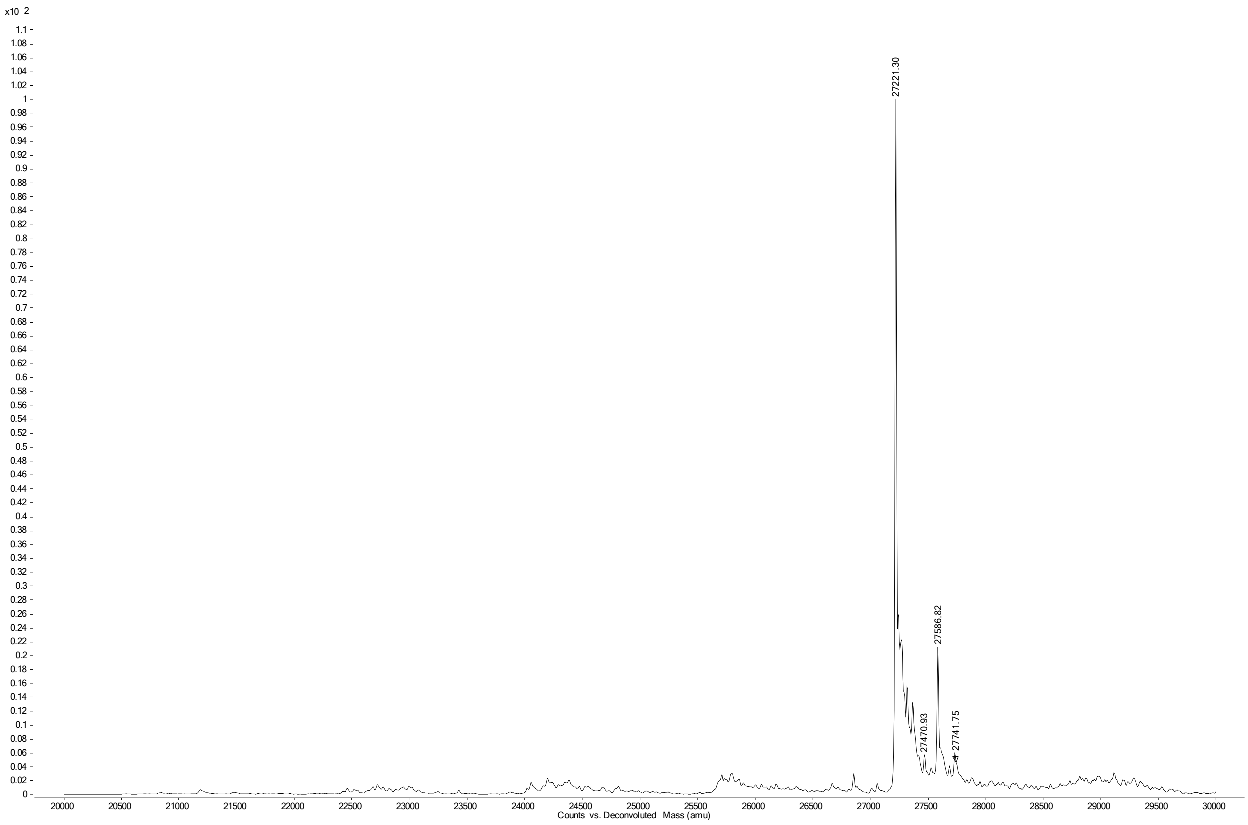


Single Fab domain

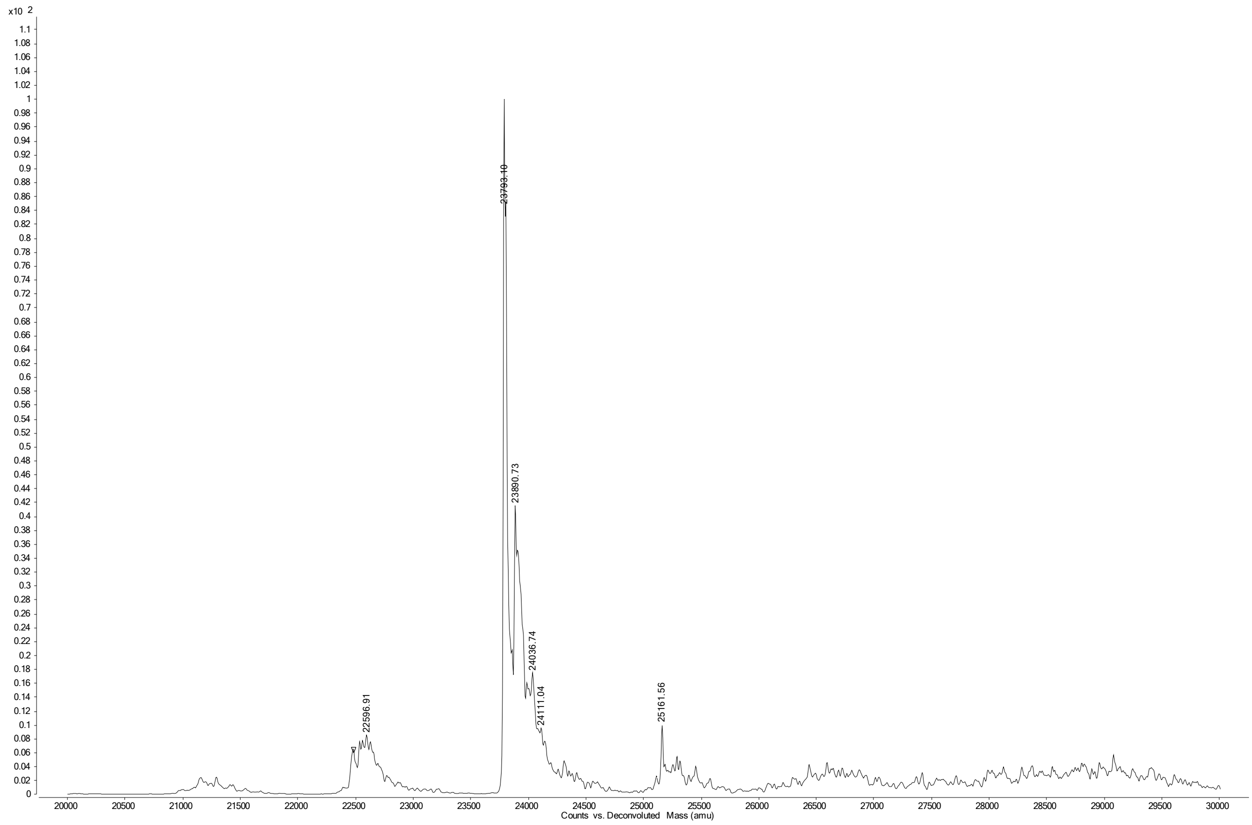


Single Fc domain

**Figure S29. Deconvoluted MS spectrum intermediate mAb 18**

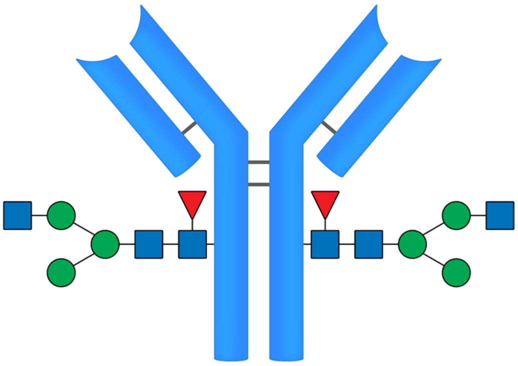

18

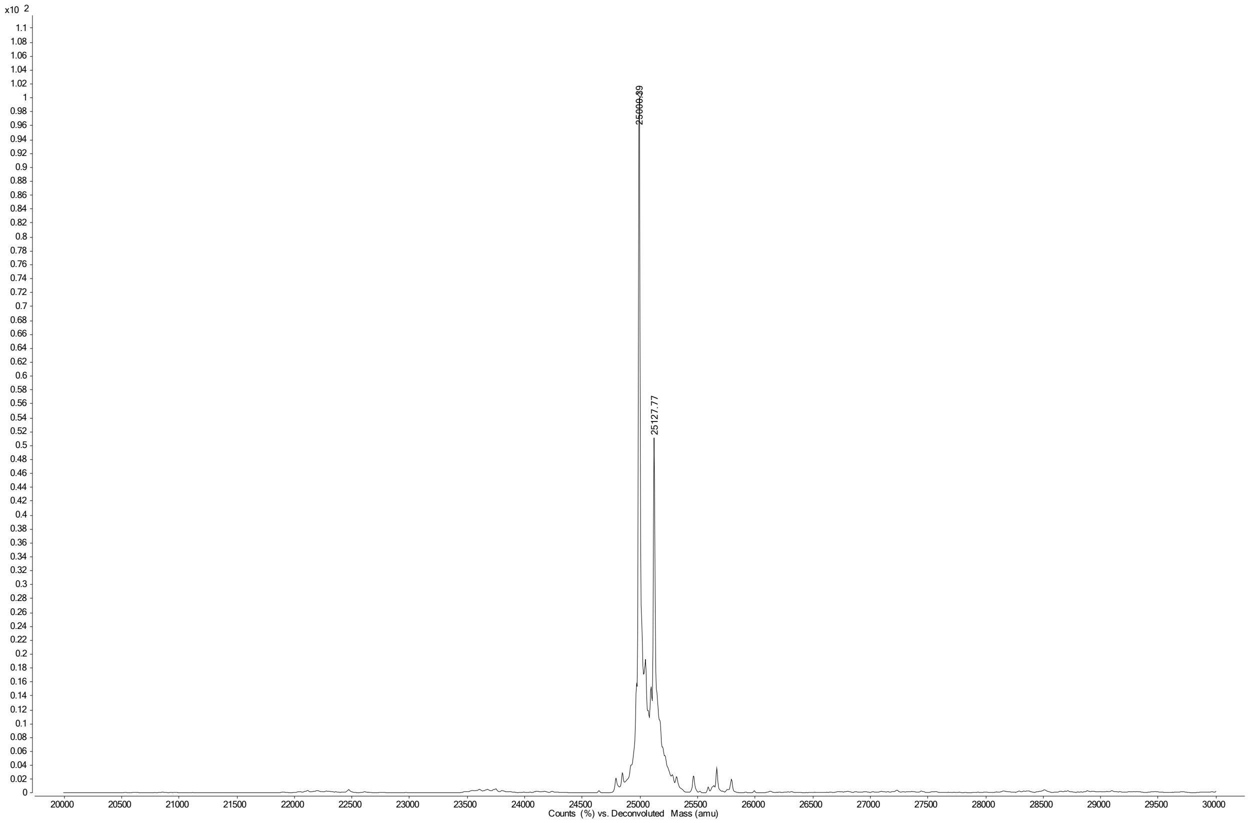

**Figure S30. Deconvoluted MS spectrum intermediate mAb 19**

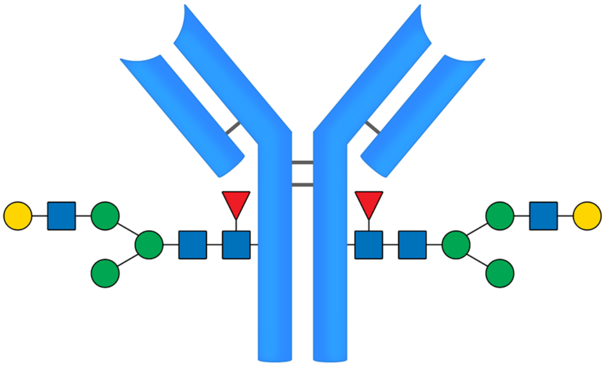

**19**

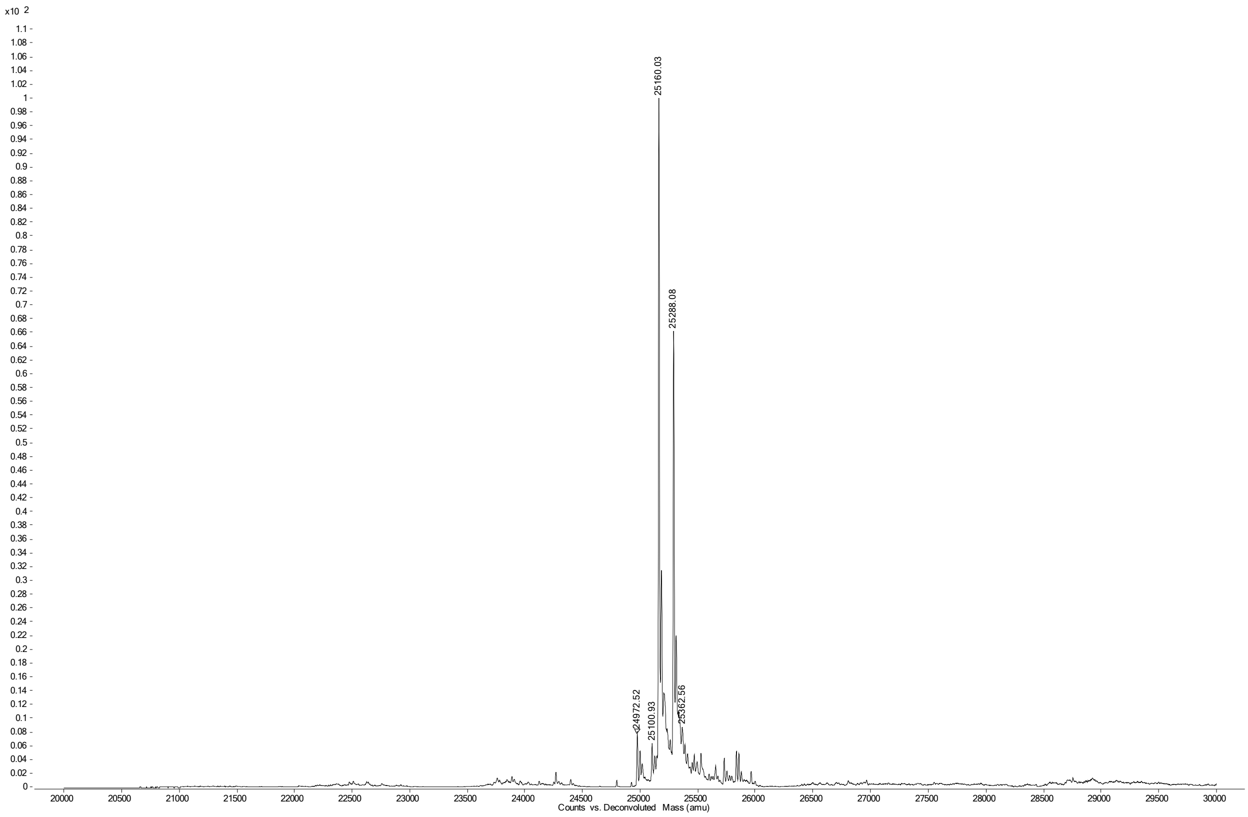

**Figure S31. Deconvoluted MS spectrum intermediate mAb 20**

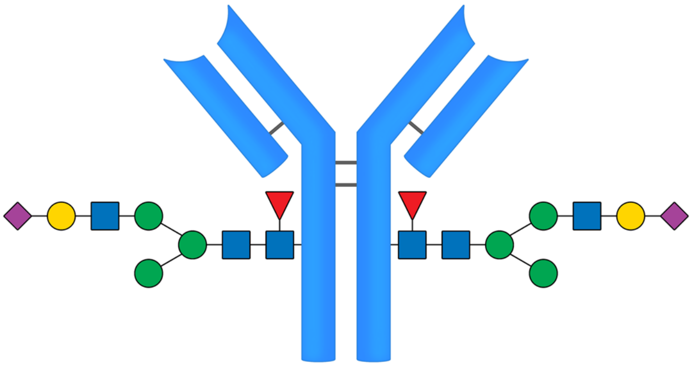

20

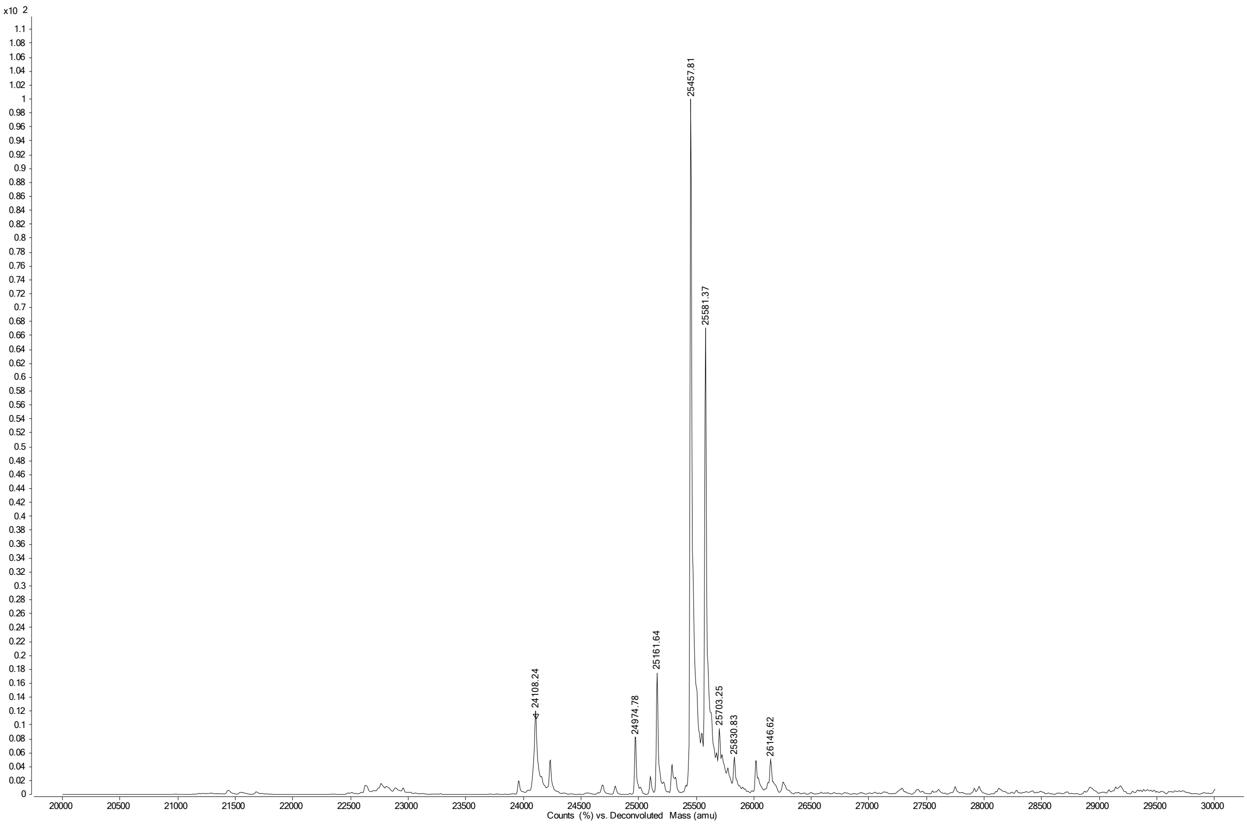

**Figure S32. Deconvoluted MS spectrum intermediate mAb 21**

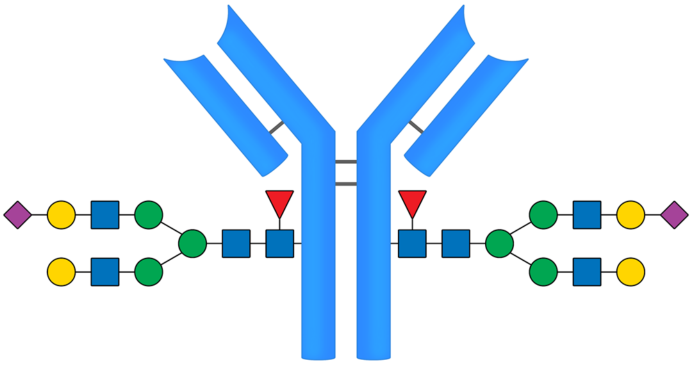

21

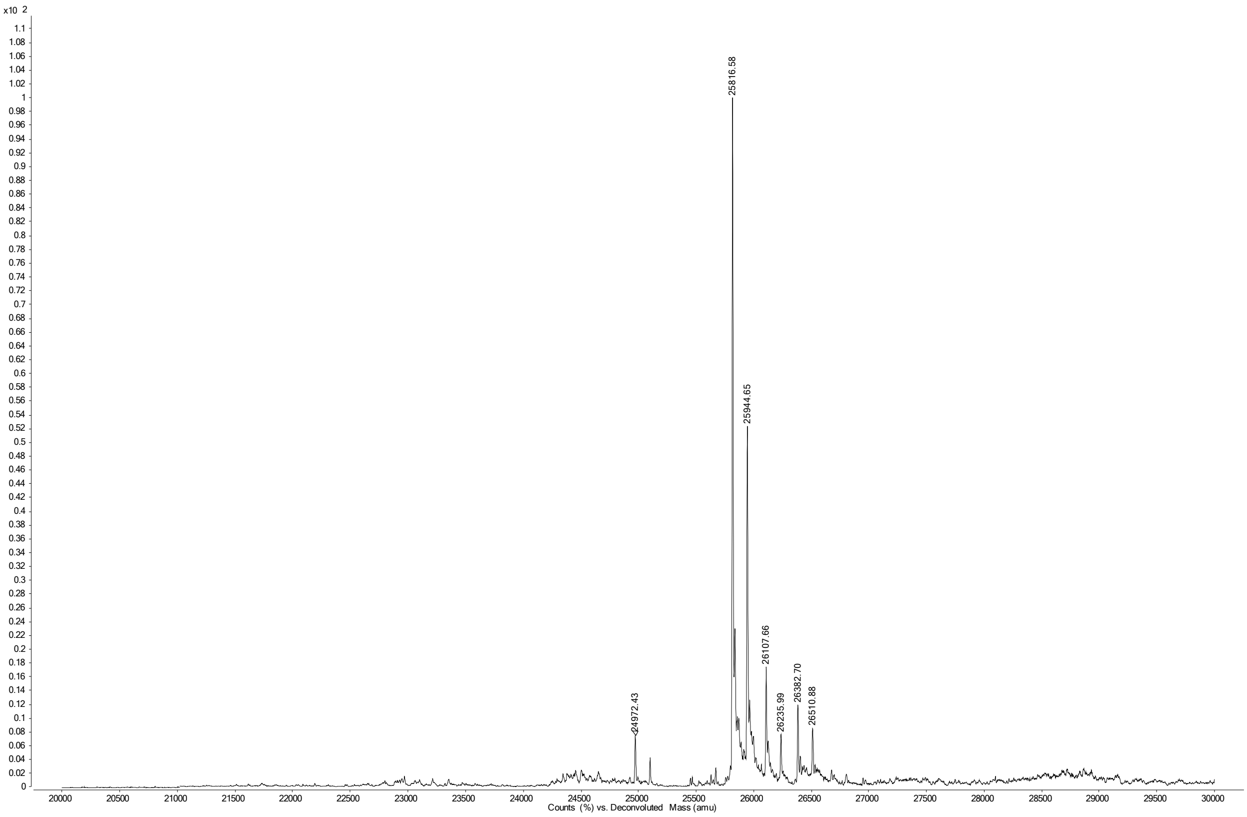

**Figure S33. Deconvoluted MS spectrum intermediate mAb 22**

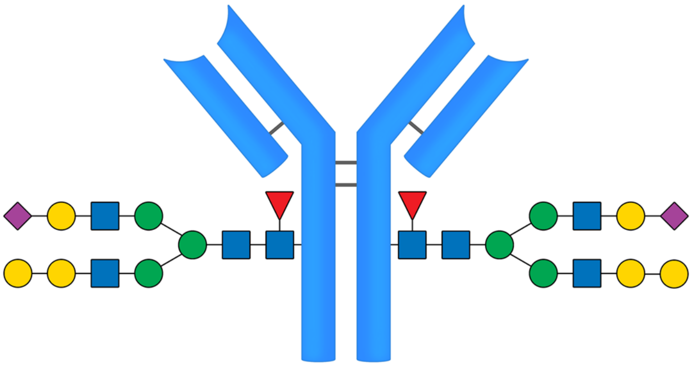

**22**

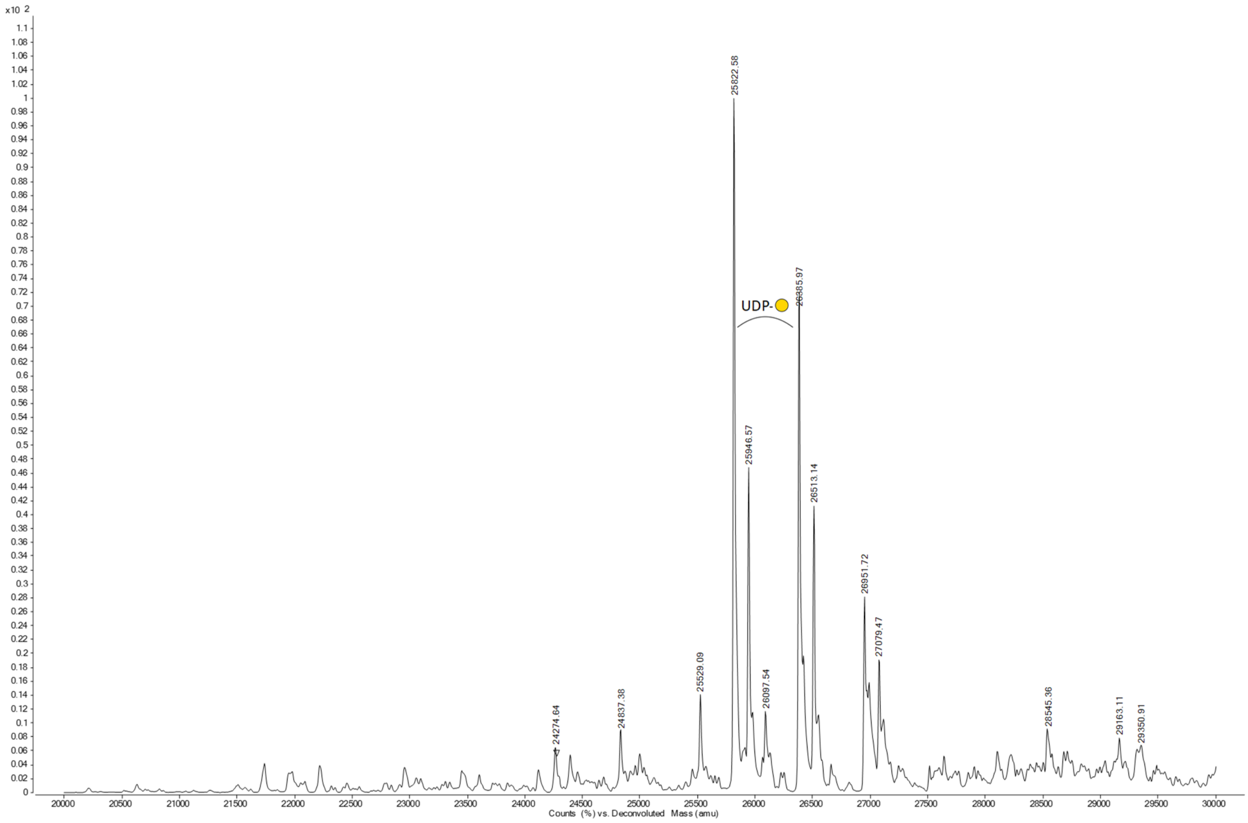

**Figure S34. Deconvoluted MS spectrum intermediate mAb 23**

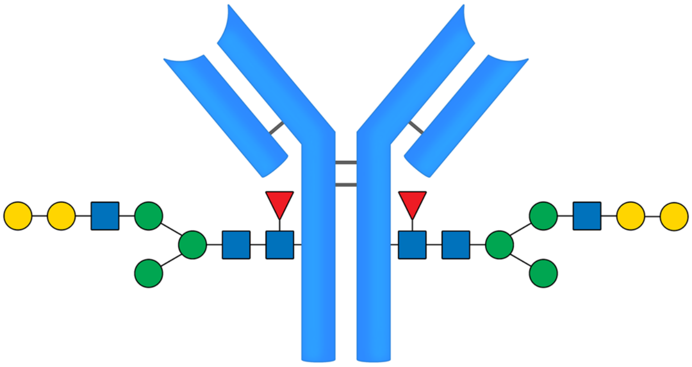

**23**

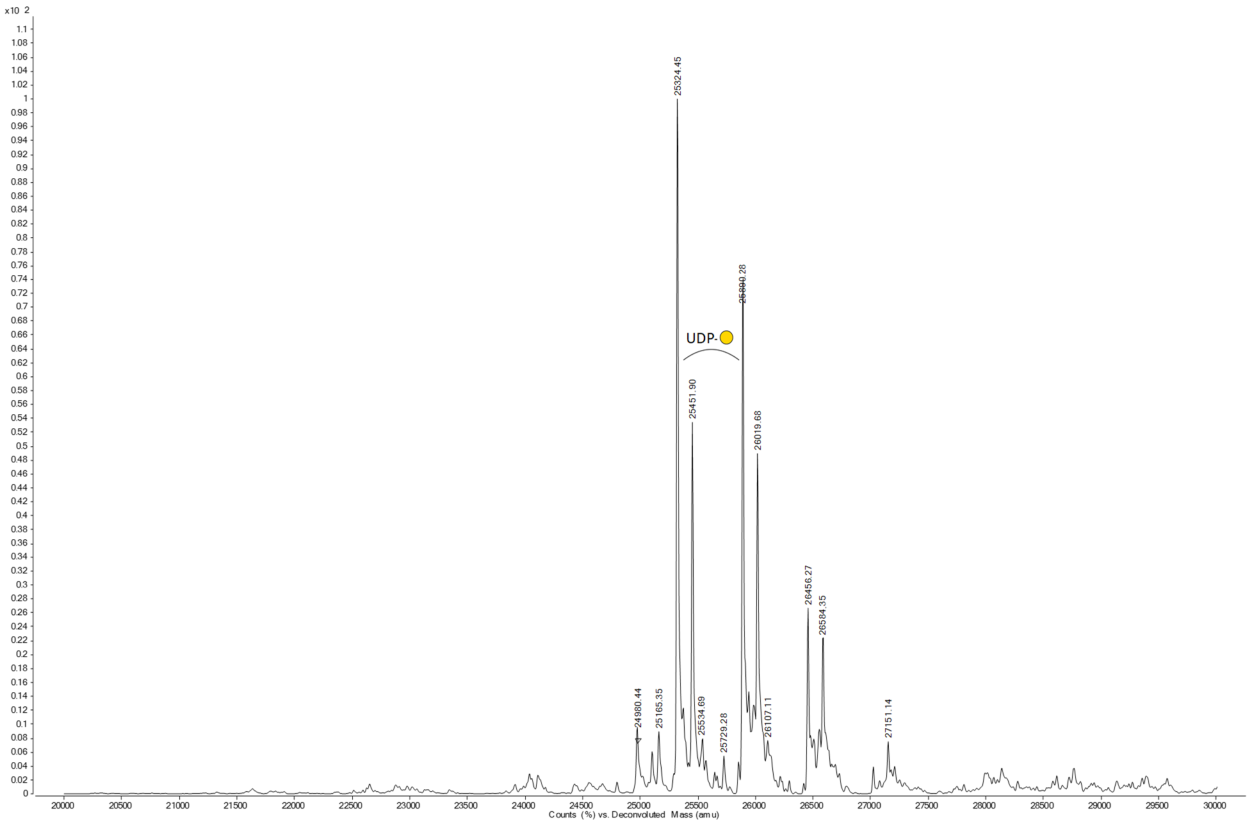

**Figure S35. Deconvoluted MS spectrum intermediate mAb 24**

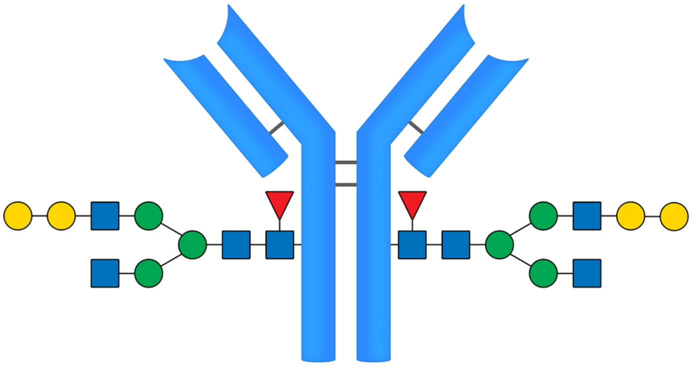

24

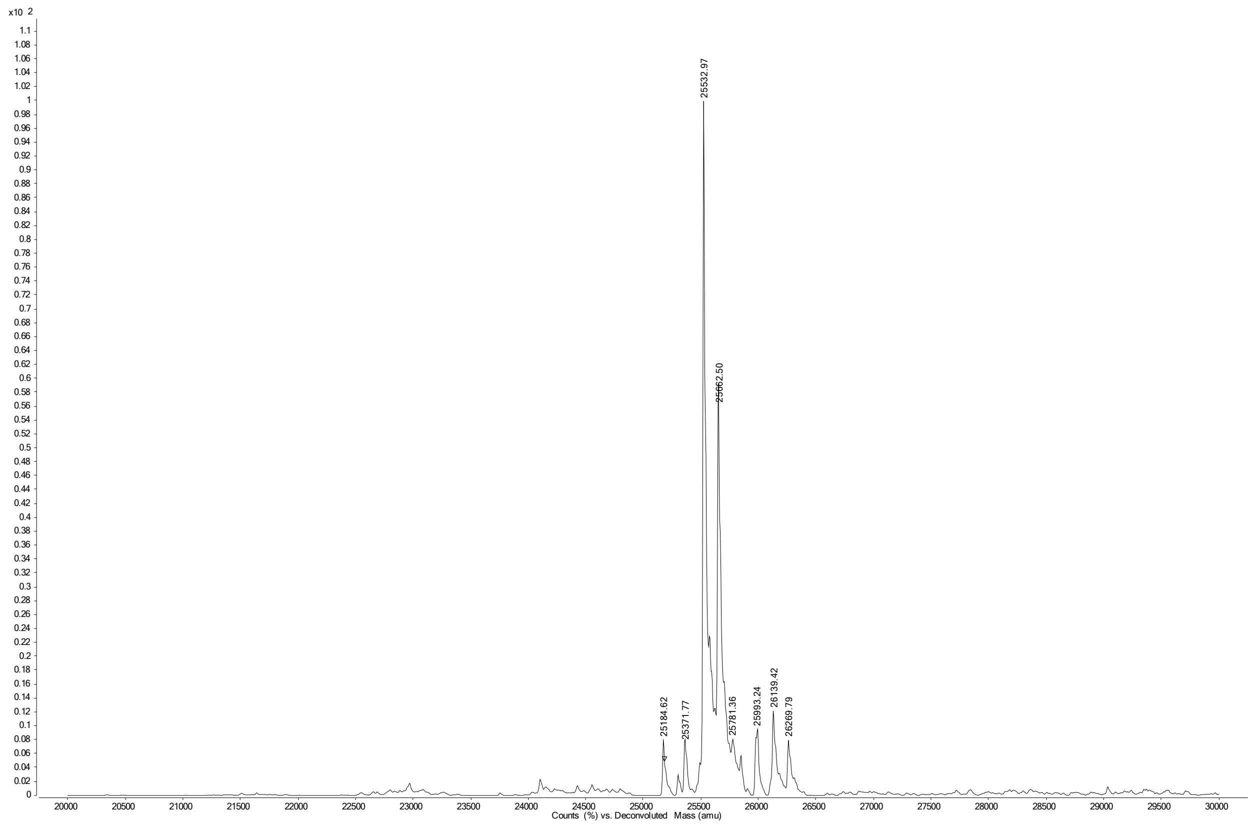

**Figure S36. Deconvoluted MS spectrum intermediate mAb 25**

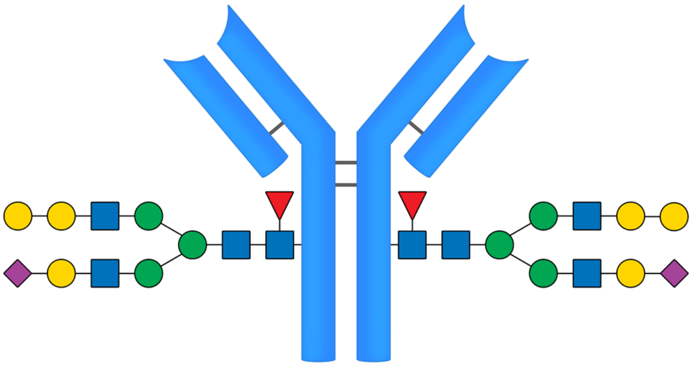

**25**

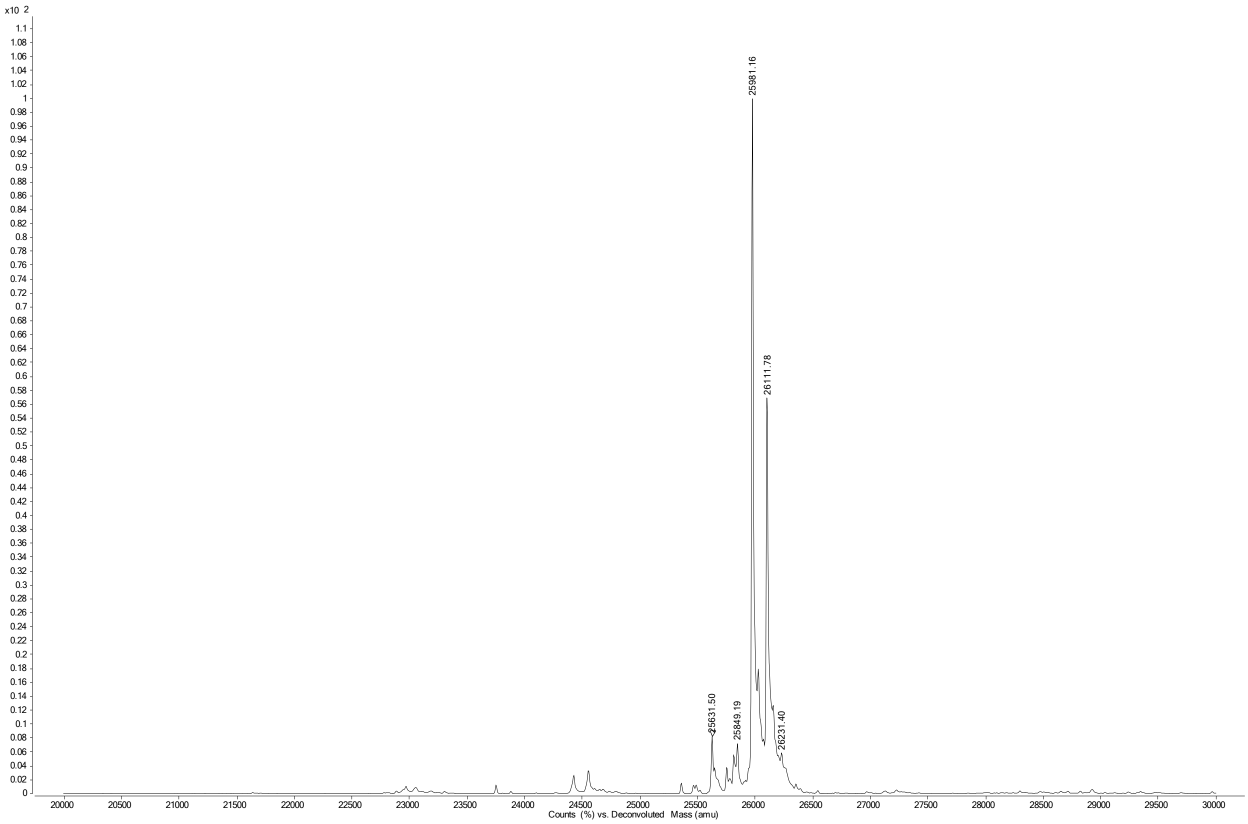

**Figure S37. Deconvoluted MS spectrum intermediate mAb 26**

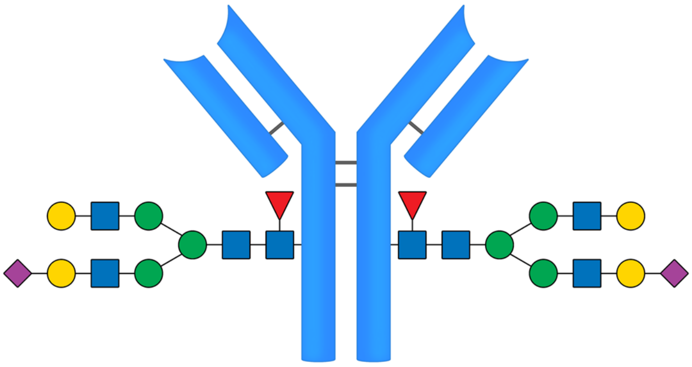

**26**

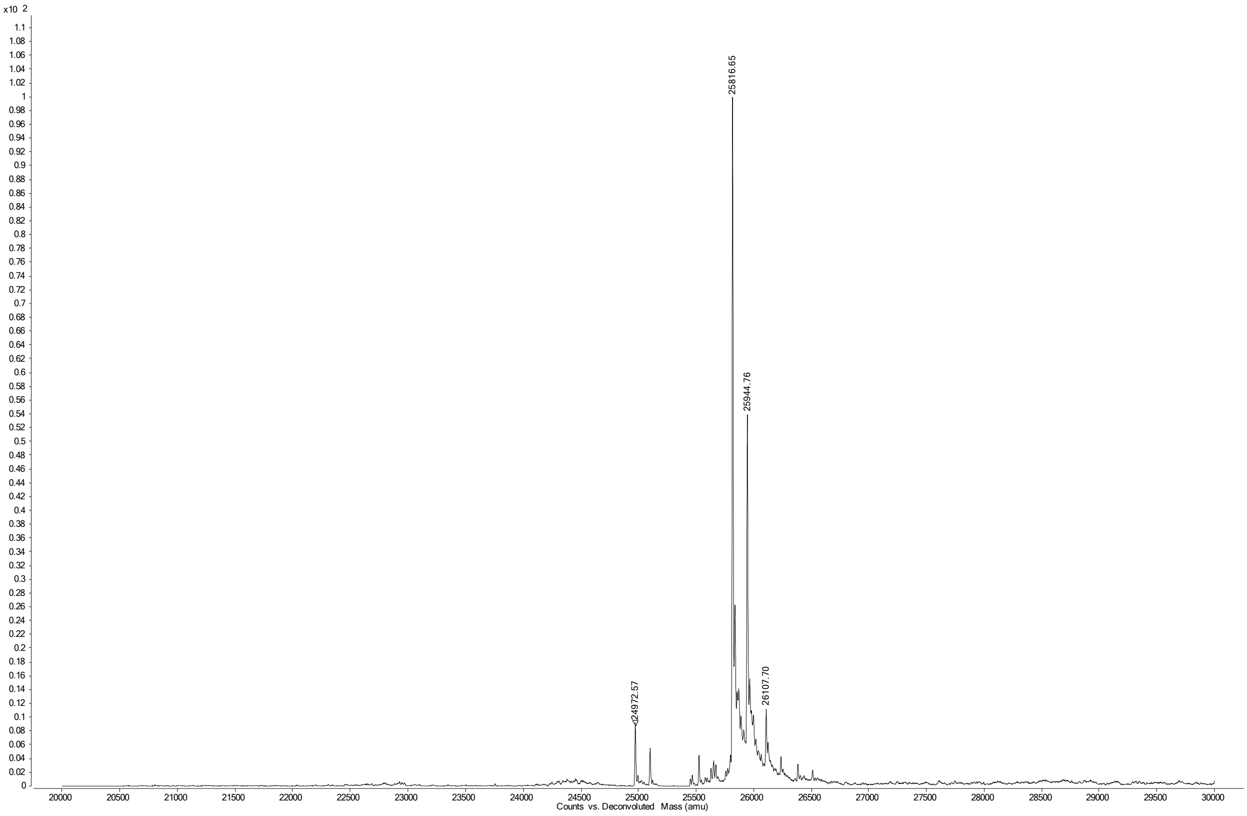

**Figure S38. SDS-PAGE gel - Coomassie stain of mAbs 1-17**

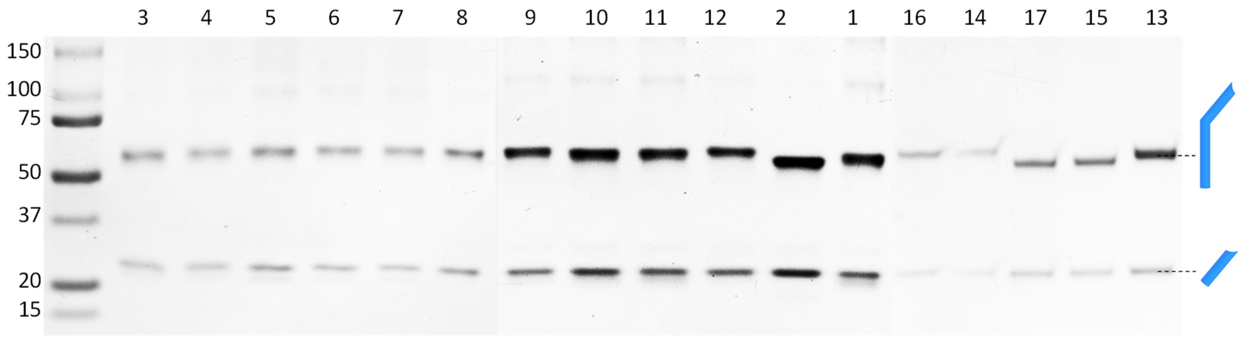


**Figure S38.** Infliximab (**1-12**) and Cetuximab (**13-17**) variants run on 4–20% Mini-PROTEAN® TGX™ Precast Protein Gel, 12-well under reducing conditions (in Laemmli sample buffer + 5% β-mercaptoethanol). Ladder: Precision Plus Protein™ All Blue Prestained Protein Standard #1610373. Stain: Coomassie (GelCode™ Blue Stain Reagent)

#### Figure S39. SPR FcγR binding sensorgrams

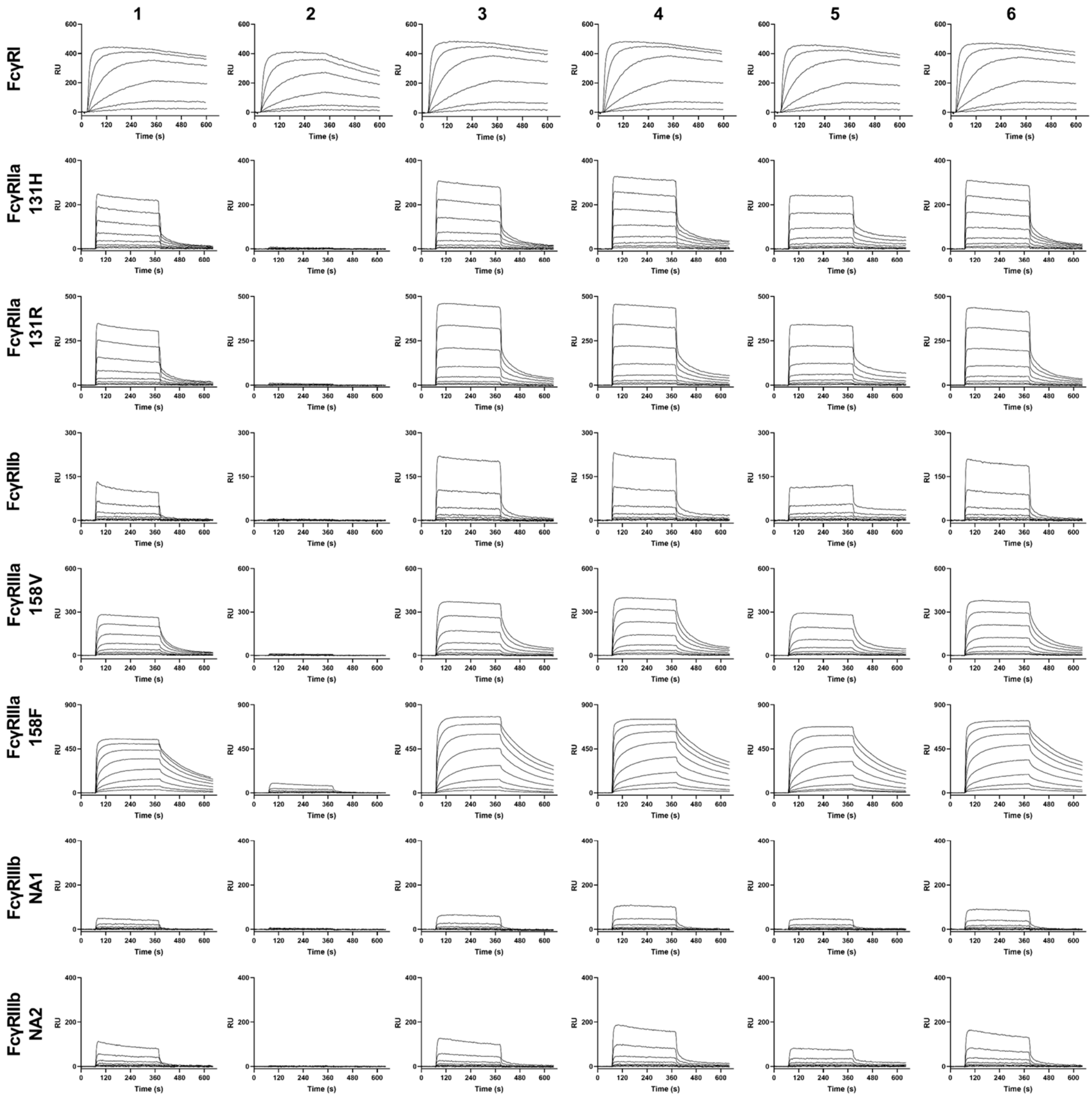

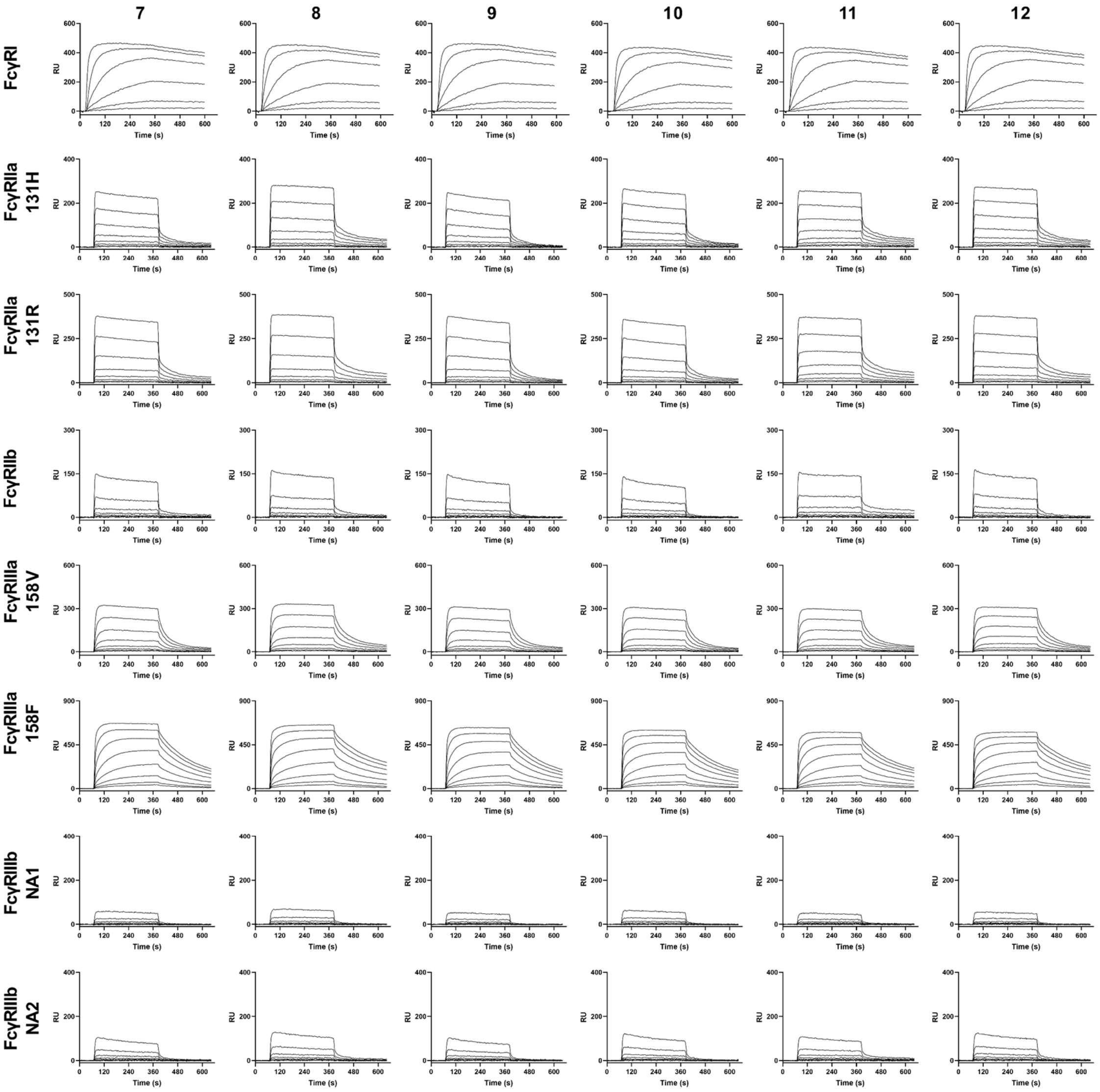
7
8
9
10
11
12
FcγRI
FcγRIIa 131H
FcγRIIa 131R
FcγRIIb
FcγRIIIa 158V
FcγRIIIa 158F
FcγRIIIb NA1
FcγRIIIb NA2
RU
Time (s)

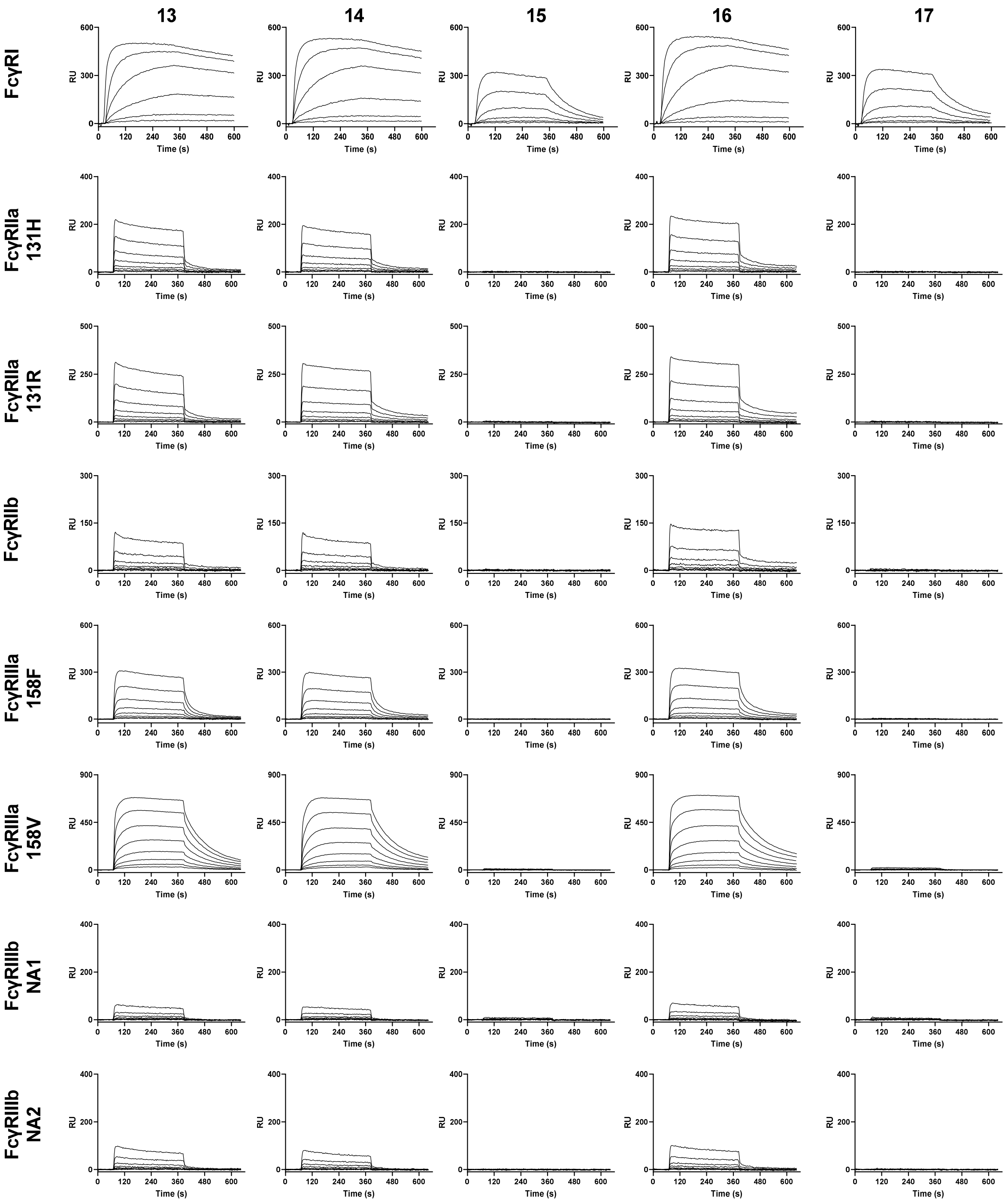
13
14
15
16
17
FcγRI
FcγRIIa 131H
FcγRIIa 131R
FcγRIIb
FcγRIIIa 158F
FcγRIIIa 158V
FcγRIIIb NA1
FcγRIIIb NA2
RU
Time (s)
